# Disproportionate regulation of trade in hunting trophies

**DOI:** 10.1101/2024.11.11.622931

**Authors:** Daniel W.S. Challender, David Mallon, Michael ’t Sas-Rolfes, Marianne Courouble, Amy Dickman, Darragh Hare, Adam G. Hart, Roseline L. Mandisodza-Chikerema, Dilys Roe, Joseph E. Mbiawa, Michael Hoffmann

**Author notes:** Corresponding author |.

## Abstract

An increasing number of countries in the Global North have enacted, or are considering, import bans on hunting trophies with the aim of protecting wildlife. We interrogate key arguments used to characterize trophy hunting and associated trade by proponents of such bans, using data from the Convention on International Trade in Endangered Species of Wild Fauna and Flora (CITES) over 25 years (2000-2024), the International Union for Conservation of Nature (IUCN) Red List of Threatened Species, and empirical research. Contrary to claims by proponents of bans, we show that trophy hunting is not a major threat to any CITES-listed species traded as a hunting trophy in 2000-2024. Arguments used to support bans often mischaracterize trophy hunting and its impacts and fail to recognize the benefits it can, and does, have for wildlife and people. Countries that have enacted or are considering import bans account for 64% of global trade in hunting trophies of CITES-listed species taken from the wild. Combined, these policies would restrict approximately one-fifth of this trade, likely with significant implications for conservation. We question whether these legislative actions are proportionate and discuss policy solutions. Wildlife trade solutions may appear simple, but evidence-based, just policies are needed to conserve biodiversity and avoid harming people.

## Introduction

People have hunted wildlife throughout history for food, clothing, household uses, trade, cultural purposes, for purported medicinal products, and recreation. In more recent times, recreational hunting for trophies by external or overseas hunters has become a more prevalent activity, with reportedly positive impacts on local livelihoods and populations of hunted species (Allen et al., 2023; Di Minin et al., 2021). Following the killing of ‘Cecil’, a lion, in 2015, legal hunting for trophies (hereafter, “trophy hunting”) has received increasing scrutiny from governments and non-governmental organizations (NGOs) and become the subject of contentious public policy debate in a growing number of countries (Ares & Sturge, 2023; Challender et al., 2024). Trophy hunting typically involves limited offtake where hunters pay a fee to hunt individual animals sometimes with particular characteristics (e.g. horn length), retaining all or part of the animal (IUCN SSC, 2012). International trade in hunting trophies from many species is regulated under the Convention on International Trade in Endangered Species of Wild Fauna and Flora (CITES), to ensure that it is sustainable, based on non-detriment findings (NDFs) (CITES, 2026). This hunting is also regulated (sub)nationally in virtually all countries in which it occurs (Roe et al., 2026).

Intense scrutiny of trophy hunting in the last decade has been led by coalitions of animal protection organizations (especially in the Global North) focused on highlighting the practice as an issue of public concern and political interest and one requiring urgent government intervention (Ban Trophy Hunting n.d.; Born Free Foundation et al., 2022). Often opposed to the killing of individuals animals for trophies regardless of outcomes for wildlife or people, these organizations have advocated for legislation to further restrict international trade in hunting trophies (Born Free Foundation et al., 2022). This has included the use of media campaigns and celebrity ambassadors, collaborating with receptive politicians, and coordinating transnational networks (Ban Trophy Hunting, n.d.). Since 2015, import bans or similar restrictions have been enacted or announced in Australia, Belgium, Finland, Germany and the Netherlands. Politicians and governments in France, Italy, Poland, Spain, the United Kingdom (UK) and the United States (US) have also considered, or are considering, import restrictions, some of which would restrict international trade in trophies from thousands of species (Supp. Mat. 1). The European Parliament has encouraged Member States to implement import bans, despite many such countries allowing extensive trophy hunting domestically (Supp. Mat. 1). Canada has also restricted the import of hunting trophies if they include rhinoceros (Rhinocerotidae spp.) horn or elephant (Elephantidae spp.) ivory (Canada Gazette, 2023). This policy proliferation suggests that trophy hunting is problematic and requires urgent policy attention or is politically sensitive; this is reflected in the arguments and moral sentiments behind bans (Born Free Foundation et al., 2022).

Here, we critique key arguments used to characterize trophy hunting and associated trade in trophies by proponents of import bans and discuss the implications for public policymaking on international trade in hunting trophies. These arguments include that: i) trophy hunting threatens or negatively impacts hunted species, ii) benefits don’t reach indigenous peoples and local communities, or are meagre, iii) trophy hunting doesn’t contribute to conservation and should be replaced with photo-tourism; and iv) there is overwhelming public support for import bans. We use data from CITES over 25 years (2000-2024; the latest available data) and the International Union for Conservation of Nature (IUCN) Red List of Threatened Species (hereafter, “Red List”), the global authority on extinction risk, and empirical research. Our methods largely follow Challender et al. (2024) and are detailed in Supp. Mat. 1 and 2. This article should be of interest to academics and conservation practitioners as well as policymakers and politicians in countries that have enacted, or are considering, tighter regulations on the trade in hunting trophies and countries where trophy hunting takes place.

## Arguments put forward for banning trade

### Trophy hunting threatens or negatively impacts hunted species

Proponents of import bans imply that trophy hunting threatens hunted species (Table 1). Between 2000 and 2024, international trade in hunting trophies from CITES-listed species in the wild involved an estimated 586,000 trophies and an estimated 447,000 whole organism equivalents (WOEs, i.e. individual animals) from 340 species and subspecies (hereafter, “species”; see Supp. Mat. 3). Data from the Red List highlights that most (69%, *n =* 236) of these species are not threatened with extinction and among those that are, trophy hunting is not likely to be a major threat to any (Fig. 1a-c., Supp. Mat. 3). Based on the Red List, trophy hunting is likely or possibly a local threat, or has been in the past, to some populations of 8 species (Fig. 1b) but this does not contribute to them being of elevated conservation concern (Challender et al., 2024; IUCN, 2024).

**Fig. 1.**
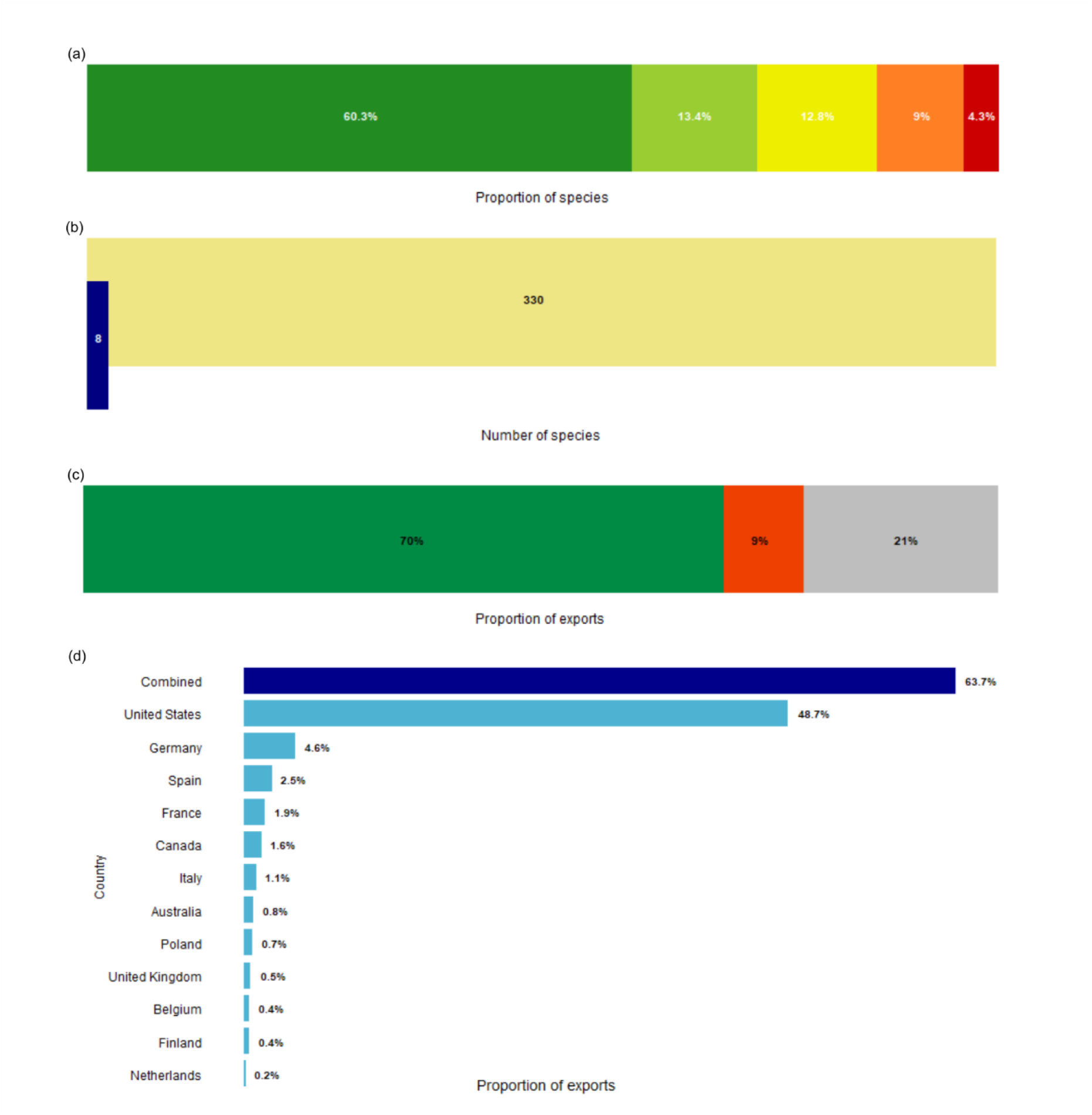
Hunting trophy trade. (**a**) Proportion of the 340 CITES-listed species sourced from the wild and traded internationally as hunting trophies (2000-2024) by Red List category: Critically Endangered (CR; red), Endangered (EN; orange), Vulnerable (VU; yellow), Near Threatened (NT; lime green), Least Concern (LC; mid-green). *n* = 320, excluding species and subspecies that are Data Deficient (DD; *n* = 2) or not assessed on the Red List (*n* = 18). (**b**) Number of these species for which trophy hunting is not a major threat (pale yellow, *n* = 330), excluding species and subspecies that lack data on threats on the Red List (*n* = 10); of these, the number for which trophy hunting is likely or possibly a local threat or has been in the past to some populations (dark blue, *n* = 8). (**c**) Proportion of hunting trophy exports (2015-2024) from countries where populations of the hunted species are considered stable, increasing or abundant (green), are decreasing (orange-red) or the status is unknown (gray) inferred from the Red List. (**d**) Proportion of direct global trade in hunting trophies from CITES-listed species in the wild (2015-2024) imported to countries that have enacted, or are considering, import bans on hunting trophies (light blue) and overall for these countries (navy blue).

**Table 1.**
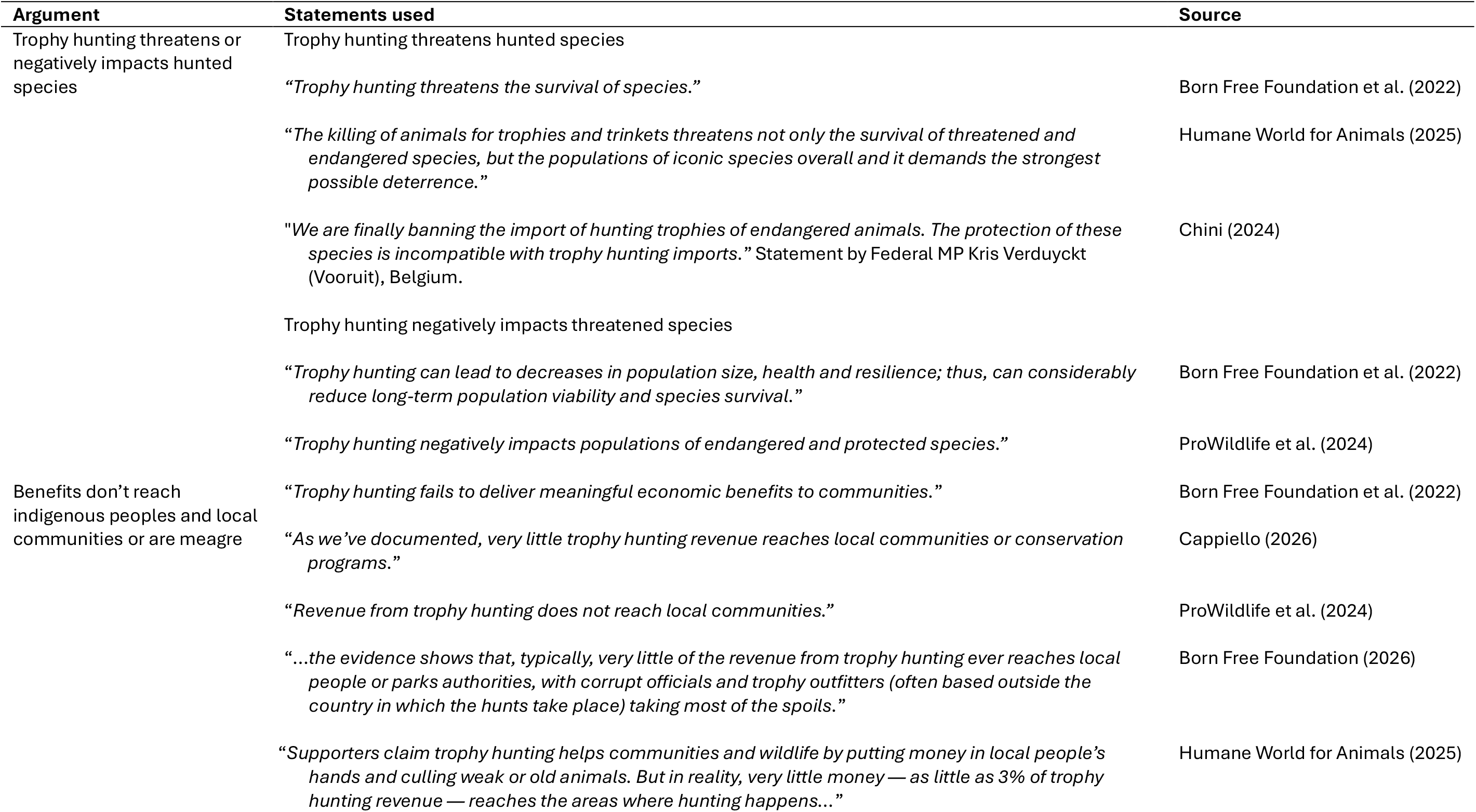
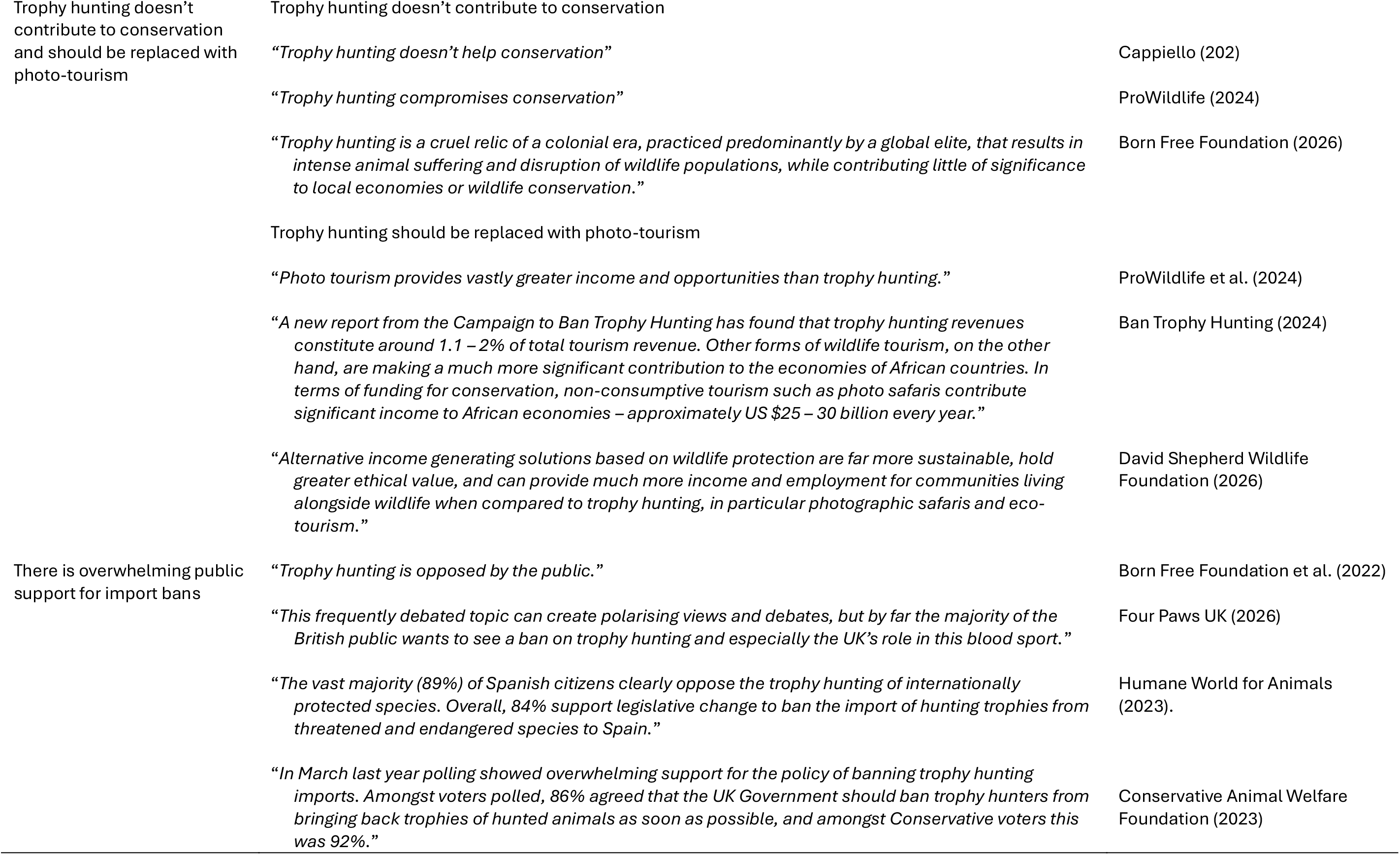
Key arguments used by proponents of import bans on hunting trophies.

Based on CITES trade data for the last decade (2015-2024) and available information in Red List assessments, trophy hunting has occurred at levels unlikely to be detrimental for most species. In this period, 198 CITES-listed species were traded internationally as hunting trophies, involving 211,000 trophies and an estimated 154,000 individual animals (Supp. Mat. 3). Ninety-eight percent of this trade involved 48 species only, including those with the highest number of trophy exports (Supp Mat. 3). For 29 of these species for which there are relevant data in Red List assessments, calculation of a relative exploitation index (REI)—based on the number of animals (WOEs) traded as trophies annually and population estimates or the number of mature individuals—indicates that for most of them exploitation pressure from trophy hunting is low relative to estimated abundance (Table 2; Supp. Mat. 2). For most of these species (*n* = 23), the number of animals involved annually was equivalent to <1% of abundance estimates. For example, for blue duiker (*Philantomba monticola*) and American alligator (*Alligator mississippiensis*), relative pressure on populations from trophy hunting was negligible. For 76% (*n* = 150) of species traded in this period, trophy exports involved fewer than 20 animals a year, and for 49% (*n* = 97), less than 1 animal a year (Supp. Mat. 3), suggesting that offtake is unlikely to be detrimental. This assessment does not consider species’ life history, density dependence, age structure, environmental variability, other threats, or other variables key to determining the sustainability of offtake (Law et al., 2021; Sutherland, 2001). It also does not consider localised governance failures and thus potential instances of unsustainable offtake and trade (Di Minin et al., 2021). However, all these exports should have been authorized based on a positive NDF by the CITES Scientific Authorities in exporting countries, a key indicator of ecological sustainability.

**Table 2.** Twenty-nine CITES-listed species traded internationally as hunting trophies (2015-2024), which comprise 80% of this trade based on number of trophies. Table includes estimated number of trophies, mean (± SD) number of trophies traded/year, estimated WOEs, mean (± SD) WOEs traded/year, and mean annual relative exploitation index (REI; %), ranked by number of trophies.

| Rank | Species | Trophies | Mean ± SD trophies/year | WOEs | Mean ± SD WOE/year | Mean annual REI (%) 2015-2024 (range) |
| --- | --- | --- | --- | --- | --- | --- |
| 1 | American black bear<br>*Ursus americanus* | 42030 | 4203 ± 1838.03 | 23681 | 2368.1 ± 1838.03 | 0.26 (0.06-0.39) |
| 2 | Nile Crocodile<br>*Crocodylus niloticus* | 36100 | 3610 ± 4664.91 | 26022 | 2602.2 ± 4664.91 | 4.34 (0.64-14.64) |
| 3 | African Savannah Elephant<br>*Loxodonta africana* | 14816 | 1481.6 ± 1803.74 | 3812.87 | 381.29 ± 1803.74 | 0.09 (0.04-0.14) |
| 4 | Hartmann's Mountain Zebra<br>*Equus zebra hartmannae* | 14397 | 1439.7 ± 380.44 | 14221.25 | 1422.13 ± 380.44 | 4.28 (2.20-5.76) |
| 5 | Giraffe<br>*Giraffa camelopardalis* | 12745 | 1274.5 ± 2250.04 | 5562 | 556.2 ± 2250.04 | 1.76 (0.36-3.25)<br>(2019-2024 only) |
| 6 | Common hippopotamus<br>*Hippopotamus amphibius* | 11882 | 1188.2 ± 298.21 | 4701.86 | 470.19 ± 298.21 | 0.38 (0.30-0.61) |
| 7 | Grey Wolf<br>*Canis lupus* | 5946 | 594.6 ± 194.55 | 5396 | 539.6 ± 194.55 | 0.42 (0.17-0.66) |
| 8 | Leopard<br>*Panthera pardus* | 4832 | 483.2 ± 123.16 | 4691.5 | 469.15 ± 123.16 | 0.2 (0.15-0.31) |
| 9 | Brown bear<br>*Ursus arctos* | 3611 | 361.1 ± 210.52 | 3439 | 343.9 ± 210.52 | 0.17 (0.08-0.39) |
| 10 | Ocellated Turkey<br>*Meleagris ocellata* | 3483 | 348.3 ± 162.2 | 3364 | 336.4 ± 162.2 | 0.96 (0.06-1.49) |
| 11 | Siberian Ibex<br>*Capra sibirica* | 3145 | 314.5 ± 158.69 | 3145 | 314.5 ± 158.69 | 0.15 (0.02-0.29) |
| 12 | Lechwe<br>*Kobus leche* | 1830 | 183 ± 53.13 | 1819 | 181.9 ± 53.13 | 0.09 (0.06-0.14) |
| 13 | Bighorn Sheep<br>*Ovis canadensis* | 1400 | 140 ± 35.94 | 1400 | 140 ± 35.94 | 0.2 (0.12-0.25) |
| 14 | Lion<br>*Panthera leo* | 1357 | 135.7 ± 57.57 | 1191 | 119.1 ± 57.57 | 0.51 (0.41-0.7) |
| 15 | Southern White Rhinoceros<br>*Ceratotherium simum simum* | 1254 | 125.4 ± 48.86 | 1219.75 | 121.98 ± 48.86 | 0.68 (0.32-1.14) |
| 16 | Wild Goat<br>*Capra hircus aegagrus* | 1168 | 116.8 ± 51.1 | 1167 | 116.7 ± 51.1 | 0.33 (0.02-0.55) |
| 17 | Blue Duiker<br>*Philantomba monticola* | 1168 | 116.8 ± 53.1 | 1168 | 116.8 ± 53.1 | 0 (0.00-0.00) |
| 18 | American Alligator<br>*Alligator mississippiensis* | 858 | 85.8 ± 173.5 | 772 | 77.2 ± 173.5 | 0 (0.00-0.00) |
| 19 | Cheetah<br>*Acinonyx jubatus* | 730 | 73 ± 22.15 | 723 | 72.3 ± 22.15 | 1.02 (0.46-1.48) |
| 20 | Great Curassow<br>*Crax rubra* | 699 | 69.9 ± 36.73 | 692 | 69.2 ± 36.73 | 0.03 (0.00-0.05) |
| 21 | Marco Polo Argali<br>*Ovis ammon polii* | 653 | 65.3 ± 60.51 | 653 | 65.3 ± 60.51 | 1.69 (1.27-2.39)<br>(2021-2024 only) |
| 22 | Wild Sheep<br>*Ovis ammon* | 632 | 63.2 ± 44.56 | 632 | 63.2 ± 44.56 | 0.15 (0.04-0.37) |
| 23 | Western Tur<br>*Capra caucasica* | 512 | 51.2 ± 43.46 | 512 | 51.2 ± 43.46 | 1.63 (0.82-2.47)<br>(2018-2024 only) |
| 24 | Bobcat<br>*Lynx rufus* | 415 | 41.5 ± 18.69 | 414 | 41.4 ± 18.69 | 0 (0.00-0.00) |
| 25 | Polar Bear<br>*Ursus maritimus* | 414 | 41.4 ± 33.72 | 280.5 | 28.05 ± 33.72 | 0.11 (0.02-0.25) |
| 26 | Created Guan<br>*Penelope purpurascens* | 402 | 40.2 ± 23.51 | 401 | 40.1 ± 23.51 | 0.01 (0.00-0.03) |
| 27 | Aoudad<br>*Ammotragus lervia* | 400 | 40 ± 22.69 | 400 | 40 ± 22.69 | 0.08 (0.00-0.33) |
| 28 | Mountain Zebra<br>*Equus zebra* | 357 | 35.7 ± 112.19 | 353 | 35.3 ± 112.19 | 0.34 (0.00-1.00)<br>(2022-2024 only) |
| 29 | European Wildcat<br>*Felis silvestris* | 320 | 32 ± 20.1 | 317 | 31.7 ± 20.1 | 0.02 (0.00-0.05) |

Similarly, based on available information in Red List assessments, an estimated 72% of trade in hunting trophies in the period 2015-2024 was from countries where populations of the hunted species are inferred to be stable, increasing or abundant (Fig. 1c., Supp. Data 1). For example, African savannah elephant (*Loxodonta africana*) trophies involved an estimated 3,813 animals, 95% of which were from southern African countries with stable or (hyper-)abundant populations, some of which are now looking to implement managed culling programs. Crucially, for species for which trophy hunting is likely or possibly a threat to some local populations or was in the past (see Supp. Mat. 2 for definitions), including lion (*Panthera leo*) and leopard (*P. pardus*), this threat is trivial relative to major threats including illegal killing and habitat loss, which are much more significant in terms of population impact, and which trophy hunting can, and does, help mitigate (CITES, 2025; Nicholson et al., 2025; Stein et al., 2025).

Proponents of import bans argue that trophy hunting negatively impacts populations of hunted species (Table 1) but fail to recognise the benefits it can provide, despite compelling evidence. In some instances, trophy hunting may negatively impact populations of hunted species (Roe et al., 2026), but this is not always the case, especially where hunting is well managed. The impact on target species also varies. For example, for American black bear (*Ursus americanus*) and brown bear (*U. arctos*), trophy hunting may contribute only to short-term population fluctuations, rather than ongoing declines (Garshelis et al., 2016; McLellan et al., 2017). For Siberian ibex (*Capra sibirica*), trophy hunting can negatively affect population demographic composition if poorly managed (Reading et al., 2020a). Other recorded impacts include behavioural, where hunted species change their movement patterns in response to hunting, for example, lion (Loveridge et al., 2016), and morphological, such as a decrease in horn size over time, for example, bighorn sheep (*Ovis canadensis*) (Pigeon et al., 2016).

Positive conservation impacts from trophy hunting, however, are not considered (or are contested) by proponents of import bans despite clear evidence of benefits. These include the removal of old, male black rhinoceros (*Diceros bicornis*), which can help achieve demographic and genetic conservation goals for the species by reducing population densities and inbreeding (’t Sas-Rolfes et al., 2022). The establishment of trophy hunting programs in Pakistan has resulted in the cessation of illegal killing and increasing populations of several caprinid species (IUCN, 2019; Khan & Baig, 2020). Indeed, many species, both hunted and non-hunted, are impacted positively by trophy hunting through the incentives it creates for conserving both the species and their habitats and the practice is frequently used to manage and conserve – beneficially – an array of species (IUCN, 2019; NACSO, 2022; Naidoo et al., 2016). Moreover, recent research highlights the favorable conservation status of mammals that are trophy hunted versus those that are not, partly because of the incentives hunting creates for wildlife management (Hill et al., 2026). The ecological, economic and social impacts of trophy hunting are therefore more nuanced than proponents of bans suggest (see Roe et al., 2026 for a review), which has important implications for public policymaking.

### Benefits don’t reach indigenous peoples and local communities or are meagre

Organizations advocating for import bans claim that benefits from trophy hunting do not reach indigenous peoples and local communities or, if they do, are meagre (Born Free Foundation et al., 2022; Table1). This includes continual use of the claim that only 3% of trophy hunting revenue reaches local communities (Table 1). However, the 3% figure has been discredited – it refers to a hypothetical hunting-operator model in Tanzania and does not reflect the range of contemporary trophy hunting programs across different parts of the world (see Booth, 2010 and Roe et al., 2026). There is also substantial compelling evidence that benefits from trophy hunting programs do flow to indigenous peoples and local communities in many instances, and frequently have meaningful impacts (IUCN, 2019; Lindsey et al., 2007; NACSO, 2022; Naidoo et al., 2016; Parker et al., 2023). The proportion of revenue that flows to these groups or stays in the local area does vary but frequently is substantial. For example, in Tajikistan, 35% of the revenue from Markhor (*Capra falconeri*) hunting licenses goes to the villages in and around the conservancies where the hunting takes place (Parker et al., 2023). In Zambia 50% of hunting revenue is given back to local communities (CITES, 2025), 80% goes to local communities in Pakistan, and it can be as much as 100% in some countries (IUCN, 2019; Parker et al., 2023).

While benefits are context-specific and may not be substantial by some Global North standards, they can be significant to marginalized rural people where hunting occurs. For example, in Namibia, per capita benefits of USD11 annually from trophy hunting are important to rural people living on a few dollars a day (NACSO, 2022). Benefits are not limited to cash income and may also include meat, housing, employment, and the provision of other services, including education through schools. In Pakistan, where trophy hunting has generated millions of dollars, revenue from hunting (e.g. of markhor *Capra spp.*) flows directly into community bank accounts and these funds have been used to build medical centers and schools, provide communities with better access to clean water, and improve rural infrastructure (Frisina & Tareen, 2009; IUCN, 2019; Roe et al., 2026). Thus, as with other forms of wildlife use, including photo-tourism, monetary and non-monetary benefits do reach indigenous peoples and local communities and can be significant for recipients but are context-dependent.

### Trophy hunting doesn’t contribute to conservation and should be replaced with photo-tourism

Groups calling for import bans frequently suggest that trophy hunting does not contribute to wildlife conservation and should be replaced by photo tourism, which they deem more ethically acceptable (Nowak et al., 2019; Table 1). As with previous arguments, however, the evidence is clear that trophy hunting can, and does, support conservation; furthermore, it cannot necessarily be replaced by photo-tourism in many specific contexts. Arguably the most significant contribution that trophy hunting makes to conservation is maintaining wildlife habitat as a viable land use (Roe et al., 2026). The most recent available estimate shows that the area of land in Africa used for trophy hunting exceeds the national park network, thus helping to conserve vast areas of land for biodiversity (Lindsey et al., 2007; Taylor et al., 2021). Trophy hunting can also support the management and protection of wild areas and species, including funding anti-poaching efforts. For instance, research in Namibia highlights that income streams from both trophy hunting and tourism are key to the success of the community conservancy model and that without the trophy hunting contribution, many conservancies would not be financially viable, threatening the existence of large areas managed for wildlife (Goergen et al., 2024; Lindsey et al., 2007; Naidoo et al., 2016). Similarly, in a recent survey of lion range countries, the most frequently stated primary aim of lion trophy hunting is revenue generation for conservation, which was assessed as very or moderately important for managing both hunting and non-hunting areas in countries including Botswana, Namibia, South Africa, Tanzania, Zambia, and Zimbabwe (CITES, 2025). In such countries, this revenue is critical to bolstering state funding for site management, including anti-poaching efforts, infrastructure maintenance, equipment and remuneration for personnel (CITES, 2025).

The scale at which trophy hunting takes place and the role it therefore plays in supporting wildlife habitat conservation in Africa, Asia and other regions (Roe et al., 2026) means it is implausible that it could be replaced entirely by photo-tourism (Lindsey et al., 2018). This would require a substantial and sustained increase in photo-tourism and associated revenue, recognizing that trophy hunting generates substantially more income per client than photo-tourism (Lindsey et al., 2007; Roe et al., 2026). It would also require a vast expansion of tourist infrastructure, which is highly unlikely because much trophy hunting takes place in remote, inhospitable and insecure landscapes, sometimes with heightened risk of disease transmission (e.g. from tsetse flies), often lacking easily viewable wildlife, and therefore of limited appeal to more mainstream photo-tourists (Lindsey et al., 2007). Consequently, replacing trophy hunting with photo-tourism is unlikely to be realistic at a meaningful scale globally. Photo-tourism can also have negative impacts on wildlife populations (e.g. through disturbance and habitat degradation), which is increasingly recognized (Dickman et al., 2018). Replacing trophy hunting with photo-tourism may also not be desirable to key stakeholders locally (Mbaiwa & Hambira, 2021; see next section) and may render some existing conservation areas financially unviable (Goergen et al., 2024; Naidoo et al., 2016).

### There is overwhelming public support for import bans

Proponents of import bans argue that there is overwhelming public support for these policies, based on ethical concerns (Ares & Sturge, 2023; Table 1). However, the public opinion polls from which these conclusions are drawn frequently fail to distinguish among types of hunting (Hare et al., 2023). There is also substantial evidence that opposition is not overwhelming. Other polls have highlighted greater acceptability of trophy hunting if it is legal and regulated, and support for import bans has been found to decline if they would negatively affect rural people or wildlife conservation (Vinogradova, 2021). Recent research indicates that the acceptability of trophy hunting depends on the attributes of a hunt, including the animal to be hunted and how the meat and revenue would be used; acceptability is generally higher for hunts that would tangibly benefit local people and support conservation or economic development (Hare et al., 2024; Ostermann-Mayashita et al., 2026). Thus, support for import bans is not overwhelming and depends on hunt attributes and the wider context.

## Sweeping bans are disproportionate policies

Proposed and enacted import bans on hunting trophies, especially indiscriminate (i.e., non-targeted or sweeping) measures, are disproportionate in conservation terms because, based on the best available data, trophy hunting is not a major threat to any CITES-listed species traded internationally as a hunting trophy in the last ∼20 years (Fig. 1a-c., Supp. Mat. 3). Additionally, trophy hunting programs can, and do, provide substantial benefits to people and wildlife (Challender et al. 2024; IUCN, 2019). Certain import bans are targeted (e.g. Australia’s ban on lion trophy trade), but most are not, may apply to hundreds or thousands of species, and constitute an overreach in public policy terms while providing no obvious conservation benefit. Key arguments used by proponents to justify these bans frequently mischaracterize trophy hunting and associated trade. They do so by failing to recognize that: i) although it may have some localized population-level impacts, trophy hunting is not a major threat to species (and there are other genuine major threats), ii) it can provide meaningful benefits to indigenous peoples and local communities where revenue allocation is supported by effective governance, iii) it can and does contribute to conservation in many cases, including by providing additional economic value to land and wildlife, iv) proposed alternatives are currently unrealistic at meaningful scales, and v) there is not necessarily overwhelming public support for bans and the acceptability of trophy hunting depends on the attributes of the hunt (Table 1). This is concerning because these arguments risk misleading policymakers and the adoption of sub-optimal public policies for both wildlife and people (Challender et al., 2022; Yeomans et al., 2022).

Indeed, there is evidence to suggest that the public and politicians in several countries may have been misled or intentionally misinformed about trophy hunting, which raises additional concerns about integrity and the use of evidence in public policymaking, as well as about accountability in animal protection organizations (Jepson, 2005; see Challender & MacMillan, 2019). Organizations advocating for import bans have sought to create moral outrage from the killing of ‘Cecil’ and to exploit it by stirring up public and political interest in trophy hunting in the hope of fast-tracking policy action (see Ban Trophy Hunting, n.d.). This has involved media campaigns (e.g. Farhoud, 2026), including social media, celebrity engagement, and parliamentary events (e.g. World Animal News, 2024). In the UK, there was a pronounced increase in coverage of trophy hunting on social media after the death of ‘Cecil’, including the use of misleading narratives, corresponding with an increase in public disapproval and proposed legislative restrictions (Yeomans et al., 2022). In the second reading of the Hunting Trophies (Import Prohibition) Bill in the UK Parliament in 2023, ∼75% of checkable statements by Members of Parliament (MPs) were factually incorrect (Dickman et al., 2024). More than 80% of these statements, where likely sources could be determined, can be linked directly to campaign group materials, indicating that these organizations have informed, if not influenced, MP positions (Dickman et al., 2024). Meanwhile, countries considering import bans, including France and the UK, have proposed legislation but failed to consult exporting countries on such policies (stricter domestic measures in CITES terms), despite established protocols to do so under CITES (e.g. in Res. Conf. 6.7).

Import bans are also poorly calibrated considering their potential impact. Most international trade in hunting trophies from CITES-listed species involves the US and Europe as the largest markets for hunting trips (Fig. 1d). More research is needed on the impacts of import bans, which risk significant adverse effects. The 12 countries that have enacted or have, or are, considering these bans account for 64% of direct global trade in hunting trophies from CITES-listed species in the wild (2015-2024) with the US the largest (Fig. 1d). Under a scenario where all these bans are enacted, this would prohibit 19% of this trade based on data for 2015-2024 (Supp. Mat. 3) and include species that command some of the highest trophy fees such as African elephant, lion, and leopard, likely with a disproportionate impact on revenue streams. A plausible outcome is that trophy hunting would decline substantially, with significant implications for conservation and rural economies. These include a reduction in monetary and non-monetary benefits to indigenous people and local communities (IUCN, 2019); reduced revenue for governments (e.g. in Southern Africa) to manage and protect both hunting and non-hunting conservation areas (CITES, 2025); reduced revenue to rural economies (e.g. lodges and taxidermists); reduced employment; weaker incentives for conservation, including on private land (Abensperg-Traun, 2009; Parker et al., 2020); an increase in other threats (e.g. habitat loss and poaching); an increase in human-wildlife conflict as tolerance for species such as African elephant and lion declines, and, ultimately, biodiversity being less competitive economically as a land use (Dickman et al., 2019; Lindsey et al., 2007; Naidoo et al., 2016). This assumes that a key motivation for hunters is to take their trophies home, which may or may not be the case (Hare et al., 2023). An alternative outcome is that trophy hunting would continue at a slightly reduced rate, as some hunters stop while the majority continue, obtaining trophies via other means (e.g. 3D printing). Under this scenario, import bans would fall short of addressing apparent concerns raised by animal protection organizations and would not reduce trophy hunting by much even if they would reduce international trade in trophies. Other outcomes are also plausible, and further research is needed on the impact of import bans and similar restrictions.

## Policy solutions

Sustainable use of biodiversity is enshrined in prominent multilateral environmental agreements including CITES and the Convention on Biological Diversity (CBD) and in regional and national policies (Hutton & Leader-Williams, 2003). All countries use and trade wildlife and manage species; for example, hunting and culling of deer (Cervidae spp.) occurs in Europe. Legal hunting for trophies and associated international trade are important parts of the wildlife economy of several countries in Africa and other regions (Naidoo et al., 2016; Roe et al., 2026), but this is an increasingly contentious issue in key importing countries. This raises the question: what should a country’s wildlife laws reflect? Should they support real and pragmatic conservation solutions that ignore public sentiment about ‘Cecil’ and trophy hunting more broadly, but which risk extending moral outrage? Or should they reflect public opinion and perceptions of what constitutes conservation (e.g. bans) and the abandonment of practical conservation because this is politically expedient, even if, as in this case, the public may have been misled? Import bans are, at least in part, based on moral outrage (Ares & Sturge, 2023; Batavia et al., 2019), which also raises the question: why are some governments pursuing import bans on trophies from the Global South but not seeking to ban the extensive trophy hunting that takes place domestically? Inevitably, policy decisions on this issue will involve trade-offs between biodiversity conservation outcomes; local livelihoods, values and benefit flows where trophy hunting occurs; animal welfare; and public sentiment in importing countries (Challender & Dickman, 2025; Ostermann-Mayashita et al., 2026).

Scientific evidence may not necessarily guide future policy decisions on trophy hunting and associated international trade (Clark et al., 2023; Salzman, 1995), but to avoid regulatory failure, public policies should be appropriately designed. Policy formulation should be informed by robust scientific evidence (e.g. on threats to species), assess cause-and-effect relationships regarding bans and other trade restrictions, and explicitly consider potential adverse consequences and likely impacts in realistic terms. Evaluation of conservation models and how wildlife trade systems function could be used to understand outcomes and impacts (e.g. ’t Sas-Rolfes et al., 2022), recognizing broader institutional arrangements (Challender et al., 2025). This includes understanding how hunters, hunting operators, and landowners may respond (Di Minin et al., 2021; Parker et al., 2020). Policy design that involves consultation with exporting countries (in compliance with international obligations under CITES) and other stakeholders on the ground, including but not limited to indigenous peoples and local communities, will better reflect the preferences of multiple stakeholders affected by these policies (Mbaiwa & Hambira, 2021). Existing impact assessments of proposed import restrictions indicate that such actors will bear most of the costs, but they were not consulted on proposed policies (DEFRA, 2021; see Challender et al., 2024). Doing so would ensure principles of social justice and legitimacy are considered in policy design (Milner-Gulland, 2024; Yeomans et al., 2022). Public opinion in importing countries, including underlying ethical frameworks (e.g. deontology and virtue ethics), may also be considered (Ghasemi, 2020), recognizing inherent policy trade-offs.

The need to consider how wildlife trade systems function, the different actors involved, and their values along international supply chains for hunting trophies reflects the socio-ecological interdependencies between distant regions, or “telecoupling” (Ostermann-Mayashita et al., 2026). Telecoupling refers to the flow of goods and services through connected but distant regions, and shared values between such regions (Liu et al., 2013). Failing to adequately consider these interdependencies increases the risk of regulatory failure and sub-optimal policies (Challender et al., 2024). Nation states can, of course, legislate without considering international impacts—an argument sometimes used in support of import bans (House of Commons, 2024)—but this does not negate the social and economic impacts in exporting countries, including on the marginalized rural poor in many cases. It would, however, undermine importing countries’ commitments to multilateralism and risk delegitimizing the CITES convention (Clark et al., 2023).

The above approach to policymaking would enable evaluation of whether further trade restrictions on hunting trophies may be warranted. For example, do they need to be applied to particular species, countries, regions, sites or a combination thereof. If warranted, appropriate regulations and controls could subsequently be devised. If not, and concerns about trophy hunting programs persist, guidance exists to improve management (IUCN SSC, 2012; Roe et al., 2025a, 2025b). Importantly, this could all be achieved within CITES.

If countries are intent on taking unilateral action, “smart bans” would be most appropriate (Webster et al., 2022). They would arguably be more legitimate because they could be used to restrict trade where there are genuine conservation concerns about hunted species but would not penalize trophy hunting where it is well-regulated and benefits species and indigenous peoples and local communities (Challender et al., 2024; Webster et al., 2022). Several conditions have been proposed to accompany smart bans. These include that trophy hunting would only be exempt from these policies if benefits from hunting tangibly contribute to species and habitat conservation, there is an equitable sharing of hunting revenues with indigenous peoples and local communities, an adaptive management and monitoring system is in place, and the hunting area has good governance (Fleming, 2024; IUCN SSC, 2012). These conditions could help incentivize good practice and raise standards in the trophy hunting industry, by requiring permit applications to demonstrate conservation and other benefits (Fleming, 2024). Smart bans would still need to be underpinned by robust evidence and involve consultations with exporting countries and other stakeholders to ensure that policies are socially legitimate and not devised by distant stakeholders who unilaterally risk harm to wildlife and people (Dickman et al., 2019).

We highlight that trophy hunting may not take place at current levels in perpetuity. Scalable alternatives that provide sustainable financing for protected and conserved areas and benefit indigenous peoples and local communities have been—and are being—considered and may prove viable in the future. Examples include novel financial mechanisms such as biodiversity bonds and credits (Medina & Scales, 2023; Thompson, 2022); however, relatively speaking, these remain in their infancy and may prove difficult to scale, particularly if they rely on voluntary markets. In an unpredictable world, more resilient conservation models are urgently needed (Lindsey et al., 2020). However, until such models and instruments can be feasibly implemented at scale, trophy hunting, if well-managed, remains a valuable tool for conservation.

Finally, it is not uncommon for animal protection organizations and researchers to make conservation-based arguments in pursuit of policy goals, but which ostensibly reflect a moral dislike of consumptive use or commercial trade in wildlife based on underlying philosophical beliefs (’t Sas-Rolfes et al., 2024, 2025; Challender & MacMillan, 2019). This can be the case even if use and trade may not threaten particular species or groups (Colston et al., 2026; Natusch et al. 2021). These perspectives are valid, but real conservation gains will not be achieved with policy prescribed by remote stakeholders. Effective conservation requires socially legitimate and just policies that address key threats to species, are informed by scientific evidence, co-developed with local actors and stakeholders, are context-specific, and appropriately coordinated at local to international levels (IPBES, 2022). Wildlife trade solutions may appear simple, but evidence-based, just policies are needed to conserve wildlife and avoid harming people.

## Data availability statement

No primary data were collected. Data from the IUCN Red List and CITES Trade Database are available online and summarised in this article and the Supplementary Material.

## Ethics approval statement

Ethics approval was not sought for this study because no primary data were collected.

## Acknowledgments

We thank A. Chidakel for interpretation of NACSO data and M.P. Louis and B. Child for advice regarding Namibia.

