## Supplementary Material for "Disproportionate regulation of trade in hunting trophies"

**Supplementary Material - Disproportionate regulation of trade in hunting trophies**


**Supplementary Material 1 – Background** **2**

Table S1. Details of proposed or enacted policies to restrict hunting trophy imports by selected countries in the Global North 2

**Supplementary Material 2 – Materials and Methods** **4**

CITES trade data 4

Table S2. Conversion factors used to estimate WOEs for hunting trophies 6

Trophy hunting as a threat to species 6

Limitations to using IUCN Red List data 9

Population status of species traded as hunting trophies 10

The impact of bans 12

Relative Exploitation Index (REI) 13

Table S3. Data used to calculate REIs 14

**Supplementary Material 3 – Results** **16**

Tables S4. CITES-listed species traded internationally as hunting trophies (2000-2024) and information on populations and threats 16

Table S5. Impact of trophy hunting on CITES-listed species for which trophy hunting is Likely/Possibly a localized threat 37

Table S6. CITES-listed species traded internationally as hunting trophies (2000-2024) and estimated trophies and WOEs 38

Table S7. CITES-listed species traded internationally as hunting trophies (2015-2024) and estimated trophies and WOEs 50

Table S8. Summary of hunting trophies and WOEs from CITES-listed species imported by countries in the Global North (2015-2024) that have enacted or are considering import bans 59

Tables S9-20. CITES-listed species imported as hunting trophies (2015-2024) to countries in the Global North and the impact of bans 60

S9. Australia 60

S9. Belgium 62

S11. Canada 65

S12. Finland 68

S13. France 70

S14. Germany 73

S15. Italy 76

S16. Netherlands 78

S17. Poland 80

S18. Spain 83

S19. United Kingdom 86

S20. Unite States 88

**Supplementary Data 1** 93

**References** 94

### Supplementary Material 1 - Background

#### Table S1. Details of proposed and enacted policies to restrict hunting trophy imports by selected countries in the Global North.

| Country | Measures proposed/enacted | Source |
|---|---|---|
| Australia | In response to Australian public concerns about "canned hunting" of African lions, Australia has, since 13 March 2015, treated African lions as though they are listed on CITES Appendix I and limits trade in African lion items, including preventing imports and exports of African lion hunting trophies. | Department of Climate Change, Energy, the Environment and Water (2021) |
| Belgium | In January 2024, Belgium agreed to ban the import of hunting trophies from ~700 species included under the EU Wildlife Trade Regulations. The exact number of species is difficult to determine but the Belgian CITES Management Authority confirms that this includes at least all Annex A species and an additional 12 species listed in Annex B and included under Annex XIII of EU Regulation 865/2006. These species are: *Ceratotherium simum simum, Hippopotamus amphibius, Loxodonta africana, Ovis ammon, O. collium, O. darwini, O. jubata, O. karelini, O. polii, O. severtzovi, Panthera leo, Ursus maritimus*. | Chini (2024); Pers. comms from Belgian CITES Management Authority to lead author (2026). |
| Canada | In November 2023, Canada amended its wildlife regulations to "*prohibit the import and export of raw elephant ivory and raw rhinoceros horn unless the specimens are destined for a museum or zoo; use in scientific research; or use in support of law enforcement activities*." As such, "*the import and export of most raw elephant ivory and raw rhinoceros horn is prohibited, including elephant ivory and rhinoceros horn hunting trophies.*" | Canada Gazette (2023) |
| European Parliament | The European Parliament's strategic objectives for CITES CoP19 included urging the Commission and the Member States "*to take immediate effective action in the framework of its commitments outlined in the EU biodiversity strategy to ban the import of hunting trophies derived from CITES-listed species.*" | European Parliament (2022) |
| Finland | In 2022, Finland approved a new nature conservation law, which bans the import of hunting trophies from all animal species listed in Appendix A and 6 species in Appendix B of the Wildlife Trade Regulations (African elephant, argali, white rhinoceros, polar bear, lion and hippopotamus). | Suomen Elainsuojelu (2023) |
| France | In 2025, two bills were submitted to the French Parliament to ban the import, export, and re-export of hunting trophies from species in Annex A and B of the EU Wildlife Trade Regulations. | Resource Africa (2025) |
| Germany | In 2022, then German Environment Minister Steffi Lemke announced restrictions on imports of hunting trophies from protected animal species. For our analysis, we assume this would apply to species in Annex A of the EU WTRs and selected species in Annex B: (southern) white rhino, African elephant, argali, lion, hippo and polar bear. | Humane World for Animals (2022) |
| Italy | In 2022, Bill no. 3430 was introduced to the Italian Parliament, which would ban the import, export and re-export of hunting trophies of species included in Appendices I and II of the CITES Convention. | Italian Parliament (2021) |

| | | |
|---|---|---|
| Netherlands | In 2016, the Netherlands banned the import of hunting trophies from over 200 animal species including those included in Annex A of the EU WTRs and several species in Annex B, including the southern white rhino, African elephant (from Botswana, Namibia, Zambia and Zimbabwe) argali, lion, hippo and polar bear. | Party for the Animals (2016) |
| Poland | In July 2024, the Polish Parliamentary Group for the Protection of Animal Rights met and discussed the need for stricter regulations on the import of hunting trophies to the country. For our analysis we have assumed this would apply to all species in Annex A of the EU WTRs and selected species in Annex B: (southern) white rhino, African elephant, argali, lion, hippo and polar bear. | World Animal News (2024) |
| Spain | In 2022, Parliamentary Association for the Defense of Animal Rights' MP presented a Parliamentary initiative calling for a ban on hunting trophy imports for species in Annex A and seven species from Annex B of the EU Wildlife Trade Regulations. Annex B species: polar bear, African elephant, lion, argali, hippopotamus, white rhino and giraffe. | Humane World for Animals (2023) |
| United Kingdom | Between 2022 and 2024 the UK debated banning the import of 6233 species on Annexes A and B of the UK and EU Wildlife Trade Regulations. | UK Parliament (2024) |
| United States | From 1 May 2024, the African elephant rule under section 4(d) of the Endangered Species Act is amended. Under this rule, African elephant range countries that export sport hunted African elephant trophies to the United States will be required to provide the U.S. Fish and Wildlife Service with an annual certification on the current management and status of their elephants and the hunting programs in their country. This will demonstrate what African range countries are doing to conserve the species and help inform decision-making on permit applications. The import of African elephant sport hunted trophies (and live African elephants and African elephant parts and products – expect ivory [regulated separately]) into the U.S. will be limited to the countries that have enacted national legislation to effectively implement the basic requirements of CITES and are designed to ensure species conservation.<br><br>In March 2024, a bill was submitted that would ban the import of hunting trophies. The Prohibiting Threatened and Endangered Creature Trophies Act of 2024 (or ProTECT Act of 2024) would modify the Endangered Species Act of 1973 to prohibit any person from taking threatened species within the United States or U.S. territorial seas as trophies or importing such trophies. A trophy is a dead animal, or recognizable part of an animal, that was obtained under an authorization issued by a state, foreign government, or private landowner. | U.S Fish & Wildlife Service (2024); U.S. Congress (2024) |

**Supplementary Material 2 – Materials and Methods**

To analyze data from the CITES trade database and IUCN Red List of Threatened Species (hereafter, "Red List") we followed Challender et al. (2024) but deviate slightly for CITES data. We detail the methods used below. We focus on CITES-listed species because international trade in these species is regulated to avoid overexploitation. We then provide details of how we determined the potential impact of import bans on trade and how we calculated a relative exploitation index (REI) for selected CITES-listed species trophy hunted and traded internationally between 2015 and 2024.

**CITES trade data**

To estimate the overall number of hunting trophies involving CITES-listed species traded internationally between 2000 and 2024, the associated number of individual animals, and the number of, and which, species and subspecies were involved (hereafter "species"), we used CITES trade data. We also used these data to estimate imports of hunting trophies from CITES-listed species, the associated number of individual animals involved, and the number of, and which species, for selected countries in the Global North. These countries have enacted or are considering import bans on hunting trophies at some level: Australia, Belgium, Canada, Finland, France, Germany, Italy, the Netherlands, Poland, Spain, the United Kingdom (UK) and the United States (US). We downloaded a comparative tabulation report on 11 May 2026 for the period 2000-2024 (CITES Trade Database, 2026; trade.cites.org/), including the latest year for which data were available (CITES Secretariat & UNEP-WCMC, 2022). We downloaded this report for all exporting and importing countries, including all sources, for the purpose of "hunting trophy" (purpose code H), and all trade terms and taxa.

Using these data but excluding trade records for which the taxon was recorded at the genus, family or order level only, and using records of direct trade only because it comprised the majority of trade by volume, we identified each unique species traded as a hunting trophy, estimated the number of hunting trophies traded internationally both overall and by species, and the associated number of individual animals in trade in the period 2000-2024. For the above countries, we identified the species imported and estimated the number of hunting trophies and associated number of individual animals imported for the period 2015-2024. This enabled us to make inferences on the sustainability of trade based on information in Red List assessments (see below).

As the focus of our analyses is hunting trophies taken from the wild, in estimating trade volumes we used source codes W (specimens taken from the wild), R (ranched specimens) and source=blank, assuming trade with the latter source code refers to specimens taken from the wild. We used all terms except “live” and “wood products” and the units hornback skins, bellyskins, number of specimens, and units=blank to estimate the number of trophies. We excluded records where trade was recorded in other units (e.g., kg). We used exporter-reported quantities, which are often more complete than importer reported quantities (CITES Secretariat & UNEP-WCMC, 2022), recognizing that this may include records of permits issued rather than actual trade volumes. However, if trade in a species was reported by importing countries only, we used importer-reported quantities. Our results for the UK differ slightly from Challender et al. (2024) because that study used exporter-reported data and selected data reported by importers; this study uses importer-reported data where not reported by exporters.

**Table S2. Conversion factors used to estimate WOEs for hunting trophies. Adapted from Harfoot et al. (2018) and following Challender et al. (2024).**

| Class | Term | Description | Factor to equate to WOE |
|---|---|---|---|
| Mammalia | BOD | Bodies | 1 |
| | FEE | Feet | 0.25 |
| | HOR | Horns | 1 |
| | SKE | Skeletons | 1 |
| | SKI | Skins | 1 |
| | SKU | Skulls | 1 |
| | TAI | Tails | 1 |
| | TEE | Teeth – *Hippopotamus amphibius* only | 0.083 |
| | TRO | Trophies | 1 |
| | TRU | Trunk – Elephantidae spp. only* | 1 |
| | TUS | Tusks – Elephantidae spp. only | 0.532 |
| Reptilia | BOD | Bodies | 1 |
| | CAR | Carapace | 1 |
| | FEE | Feet | 0.25 |
| | SKE | Skeletons | 1 |
| | SKI | Skins | 1 |
| | SKU | Skulls | 1 |
| | TRO | Trophies | 1 |
| Aves | BOD | Bodies | 1 |
| | SKE | Skeletons | 1 |
| | SKI | Skins | 1 |
| | SKU | Skulls | 1 |
| | TAI | Tails | 1 |
| | TRO | Trophies | 1 |

*This includes records for *Elephas maximus*, *Loxodonta africana, and L. cyclotis*.

We estimated the number of whole organism equivalents (WOEs), that is, the number of individual animals, associated with trade volumes, adapting the approach by Harfoot et al. (2018) and following Challender et al. (2024). One animal may produce one or more trophies meaning that the number of trophies does not necessarily equate to the number of animals killed. Converting trade volumes to WOEs enables estimates of the number of animals involved in trade (Harfoot et al. 2018). We converted products in trade into WOEs (e.g., 5 bodies equate to 5 WOEs), using the conversion factors in Table S2 for trade records with the units hornback skins, bellyskins, number of specimens, and unit=blank. We estimated the number of WOEs associated with trade in species, countries, species-country combinations and overall, by the summarizing data. We used pivot tables in MS Excel and RStudio version 1.4.1617 to summarize the data and calculate means and standard deviations for trade in trophies and WOEs over time.

**Trophy hunting as a threat to species**

To classify species based on the level of threat, if any, from trophy hunting, we expanded the database reported in Challender et al. (2024) to include all unique CITES-listed species traded as hunting trophies between 2000 and 2024. We omitted 4 exclusively marine species (e.g., Pacific White-sided dolphin *Lagenorhynchus obliquidens*) because although these species were traded as hunting trophies, they are unlikely to have been hunted for trophies in line with our definition (see below). This resulted in 340 species in our database. We updated our database in May 2026 to ensure the data were up to date using the latest version of the Red List (version 2025-2; IUCN Red List, 2026).

The database contains the following fields: class, common name, scientific name, date of last assessment, Red List Threat Category, population size, population trend, and whether intentional hunting of the species is a current threat (yes/no) based on the following threat codes: 5.1.1. (Hunting & collecting terrestrial animals → Intentional use (species being assessed is the target)), 5.4.1. (Fishing & harvesting aquatic resources → Intentional use: subsistence/small scale (species being assessed is the target)), and/or 5.4.2. (Fishing & harvesting aquatic resources → Intentional use: large scale (species being assessed is the target)). The database also includes a field for which of these threat codes had been applied to species (e.g., 5.4.1.) and the timing of those threats (e.g., ongoing or past, unlikely to return). More information on how the Red List generally defines major threats is available (IUCN, 2023). The database also includes a field containing the narrative text from the threats and justification fields respectively from assessments and relevant information from the Use and Trade section of each assessment where this information was available. The Use and Trade Classification Scheme includes a specific category (“Sport hunting/specimen collecting”), which provided a helpful check when classifying species (see below), although this field is occasionally incorrectly coded when the end use is for food or other purposes. There is no direct relation between the use and trade scheme and the threat scheme on the Red List; rather, the Use and Trade classification scheme is intended to document use at any level whether a threat or not. We also included information on which CITES Appendix species are in, or if they are not listed under CITES, using information for species in Species+ (UNEP, 2026).

Following Challender et al. (2024), we classified species based on whether legal hunting for trophies is i) likely a major threat contributing to species being of elevated conservation concern (i.e., to a species meeting, or approximating, the thresholds for listing in any of categories NT [Near Threatened], VU [Vulnerable], EN [Endangered], CR [Critically Endangered] or EW [Extinct in the Wild], as defined by IUCN, ii) likely or possibly causing localized declines, or iii) not a threat to species.

To classify species in the above categories we read the narrative text in the threats and justification fields of each assessment and interpreted this information with available information on coded threats (e.g., 5.1.1. [Hunting & collecting terrestrial animals → Intentional use [species being assessed is the target]]). We considered the timing (e.g., past), scope (e.g., minority of the population), and severity (e.g., causing rapid declines) of threats (where these data were available), to assist in distinguishing between major and minor threats (IUCN, 2016). We read the Use and Trade field of assessments for additional context and used information on use and/or trade to inform decision-making. Following Challender et al. (2024), we categorized legal hunting for trophies as a major threat contributing to species being of elevated conservation concern where available evidence indicated that a species has intentional use as a major threat (based on the threats narrative and/or coded threats, including timing, scope, and severity), and the threats narrative indicates that legal hunting for trophies as defined (see below) is currently a primary factor driving this threat. Where this was not the case, species were not considered to have legal hunting for trophies as a major threat. We then categorized these species based on whether there is evidence indicating that legal hunting for trophies is likely (i.e., probable) or possibly (i.e., stated but qualified as uncertain, e.g., "potentially" or similar) causing localized declines but is not a major threat, i.e., not contributing to a species being of elevated conservation concern. Species that did not meet either of these criteria were considered to not be threatened by legal hunting for trophies at any level. We used the definition of trophy hunting in Challender et al. (2024) which draws on (IUCN SSC, 2012): "*legal, low-offtake hunting, where hunters pay a high fee to hunt individual animals with particular characteristics (e.g., horn length) and retain all or part of the animal.*"

**Limitations to using IUCN Red List data**

Our assessment of whether trophy hunting is a threat to species, or not, is based on a direct interpretation of information contained in the Red List, which may not be current or complete. As such we cannot, and do not, purport to definitively state that trophy hunting is not a major threat to any species; instead, we highlight that, based on the best available data in the Red List, this does not currently appear to be the case. Yet, as 97% of the species in our dataset (*n* = 330) have been assessed on the Red List or have some information available on threats in the assessment, and 86% of species (*n* = 293) have assessments that are less than 10 years old (i.e., do not need updating), we are confident our approach is robust. However, we note that focusing on CITES-listed species means that we do not include other species for which trophy hunting may be causing localized or potentially species-level declines (e.g., the Mountain Nyala) (IUCN SSC Antelope Specialist Group, 2017a).

**Population status of species traded as hunting trophies**

For the 198 CITES-listed species traded as hunting trophies (2015–2024) we determined the population status (e.g., increasing, decreasing, stable or unknown) of each species for each exporting country. We reviewed available information for each exporting country in the global Red List assessment for each species (using Red List version 2023-1, then checked against 2025-2) or regional Red List assessments where they were available. For a small number of species more recent population estimates were available in sources such as CITES documents (e.g., listing proposals) or other publications (e.g., journal articles); these sources were used where this was the case. We calculated the proportion of trade overall sourced from countries where populations of the hunted species are increasing, decreasing, stable or similar.

A note on taxonomy. The CITES nomenclature sometimes deviates from the Red List and can result in anomalies. For example, several species of baboon (*Papio* spp.) are indicated as exported from South Africa (ZA), even though only a single species (*P. ursinus*) occurs there. We highlight such cases (see SD1) and present these population trends as “Indeterminate”. These trends were not used in our calculations of species status. In additions, for several species (see Tables S4 and S6) the number of traded trophies relate wholly to extralimital populations and therefore have no impact on wild populations. These were also excluded from our calculations of species status.

**The impact of bans**

To quantify the impact of proposed and enacted import bans on future international trade in hunting trophies from CITES-listed species, we identified which species are or would be the subject of such bans for each country. For EU countries we determined which annex of the Wildlife Trade Regulations (WTRs) imported species are on and whether these species would be in-scope or out of scope of bans. To do this, we used species listings in the WTRs downloaded from Species+ (UNEP, 2026) on 26 May 2026. For the United States, we checked whether species are on the Endangered Species Act or not by using an up-to-date species list downloaded on 27 May 2026. For Germany and Poland exact details on proposed legislation were not available (see Table S1) and so we assumed import bans for these countries to be similar in scope to those enacted by Belgium, the Netherlands and Finland, involving all species listed in Annex A and selected species in Annex B of the WTRs. Our assessment focuses on CITES-listed species only whereas the proposed PROTECT Act in the US refers to many species not listed in the CITES Appendices and which are therefore out of scope of this study.

**Relative Exploitation Index (REI)**

For 29/48 species that compromised 98% of international trade in hunting trophies involving CITES-listed species in the period 2015-2024, we calculated a relative exploitation index (REI) using available information in Red List assessments. To do so, we used the number of animals traded as trophies annually (i.e. WOEs) as the numerator and population estimates or number of mature individuals (based on which was available in assessments) as the denominator. We calculated an annual REI for 2015-2024 and report both the mean REI and the range for this period in percentages in the main text. We calculated an REI for 29 species because the assessments for these taxa contained population estimates or number of mature individuals, or both. Where both were present we used population estimates (see also below). To determine the most appropriate population estimate to use (e.g. global, regional, or national), we examined where hunting trophies were exported from and used the most appropriate estimate based on these exports. For some species this was global, for others it was regional (Table S3). Where population estimates were presented as intervals we used the mid-point. We excluded extra-limital and introduced populations.

We acknowledge that this REI is not a direct estimate of annual exploitation rates for species but does provide *a* measure of relative exploitation pressure from trophy hunting on populations at a coarse scale, informed by the best available information on the Red List and informed by CITES trade data. However, it does not consider population growth rates, demographic effects, or previous or more up to date population estimates.

**Table S3. Exporting countries and population estimates used to calculate REI for 29 CITES-listed species traded as hunting trophies (2015-2024). Source: IUCN Red List (2026) and CITES Trade Database (2026).** AM = Armenia, AZ = Azerbaijan. BG = Bulgaria, BJ = Benin, BW = Botswana, CA = Canada, CF = Central African Republic, CG = Congo, CM = Cameroon, DE = Germany, EE = Estonia, ES = Spain, ET = Ethiopia, FI = Finland, FR = France, HR = Croatia, HU = Hungary, KG = Kyrgyzstan, KZ = Kazakhstan, LT = Lithuania, LV = Latvia, MA = Morocco, MK = North Macedonia, MN = Mongolia, MX = Mexico, MZ = Mozambique, NA = Namibia, PK = Pakistan, RO = Romania, RS = Serbia, RU = Russian Federation, SE = Sweden, SI = Slovenia, TD = Chad, TJ = Tajikistan, TR = Türkiye, TZ = United Republic of Tanzania, UG = Uganda, US = United States of America, XX = Unknown, ZA = South Africa, ZM = Zambia, ZW = Zimbabwe.

| Scientific name | Common name | Exporting countries (2015-2024) | Population estimate | Population notes |
|---|---|---|---|---|
| Ursus americanus | *American Black Bear* | CA, US | 850,000-950,000 | Global estimate |
| Crocodylus niloticus | *Nile Crocodile* | BW, MZ, NA, TZ, UG, ZM, ZW | 50,000-70,000 | No. of mature individuals |
| Loxodonta africana | *African Savanna Elephant* | BW, CM, MZ, NA, TZ, XX, ZA, ZM, ZW | 415,428 | Continental estimate |
| Equus zebra hartmannae | *Hartmann's Mountain Zebra* | NA, ZA | 33,265 | No. of mature individuals |
| Giraffa camelopardalis | *Giraffe* | BW, NA, ZA, ZM, ZW | 52,569 | Regional estimate* |
| Hippopotamus amphibius | *Common Hippopotamus* | BJ, BW, CM, MZ, NA, TZ, UG, ZA, ZM, ZW | 115,000-130,000 | Global estimate |
| Canis lupus | *Grey Wolf* | BG, CA, EE, ES, KG, KZ, LT, LV, MK, MN, RO, RU, SE, TJ, TR, US | 130,000 | Global estimate |
| Panthera pardus | *Leopard* | BW, CF, ET, MZ, NA, RU, TZ, US, XX, ZA, ZA, ZM, ZW | 231,636 | Global estimate |
| Ursus arctos | *Brown Bear* | AM, CA, DE, EE, FI, FR, HR, HU, KZ, RO, RU, SE, SI, US | 200,000 | Global estimate |
| Meleagris ocellata | *Ocellated Turkey* | MX | 20,000-49,999 | No. of mature individuals |
| Capra Sibirica | *Siberian Ibex* | KG, MN, PK, RU, TJ | 170,000-250,000 | Global estimate |
| Kobus leche | *Lechwe* | NA, ZA, ZM | 212,000 | Global estimate |
| Ovis canadensis | *Bighorn Sheep* | MX | 70,000 | Global estimate |
| Panthera leo | *Lion* | BJ, BW, CM, ET, MZ, NA, TZ, XX, ZA, ZM, ZW | 22,000-25,000 | Global estimate |
| Ceratotherium simum simum | *Southern White Rhinoceros* | NA, ZA | 18,064 | Global estimate |
| Capra hircus aegagrus | *Wild Goat* | AM, TR | 35,000 | Armenia and Türkiye only |

| | | | | |
|---|---|---|---|---|
| Philantomba monticola | *Blue Duiker* | CF, CG, CM, MZ, TZ, ZA, ZM | 7,000,000 | Global estimate |
| Alligator mississippiensis | *American Alligator* | US | 3,000,000-4,000,000 | Global estimate |
| Acinonyx jubatus | *Cheetah* | NA, ZA, ZW | 7,100 | Global estimate |
| Crax rubra | *Great Curassow* | MX | 50,000-499,999 | Global estimate |
| Ovis ammon polii | *Marco Polo Argali* | KG | 4,104 | Kyrgyzstan only |
| Ovis ammon | *Wild Sheep* | KG, MN, TJ | 42,235 | Kyrgyzstan, Mongolia, and Tajikistan only |
| Capra caucasica | *Western Tur* | AZ, RU | 4,000-5,000 | Global estimate |
| Lynx rufus | *Bobcat* | CA, MX, US | 2,352,276-3,571,681 | Global estimate |
| Ursus maritimus | *Polar Bear* | CA | 26,000 | Global estimate |
| Penelope purpurascens | *Crested Guan* | MX | 50,000-499,999 | No. of mature individuals |
| Ammotragus lervia | *Aoudad* | MA | 5,000-10,000 | Global estimate |
| Equus zebra | *Mountain Zebra* | NA, ZA | 34,979 | No. of mature individuals |
| Felis silvestris | *European Wildcat* | ET, NA, RO, RS, TD, TZ, ZA, ZW | 140,000 | Global estimate |

*Estimate based on the total population of all subspecies that occur in exporting countries.

### Supplementary Material 3 – Results

**Table S4. CITES-listed species and subspecies traded internationally as hunting trophies (2000-2024) (*n* = 340), CITES Appendix, IUCN Red List Category, population size, population trend, information on intentional hunting/harvesting threats, whether trophy hunting is a major threat to species or not, and whether trophy hunting is likely or possibly a localized threat to species but does not contribute to the species being of elevated conservation concern.** Seventeen of these species/subspecies have not been evaluated on the Red List (NE in the fourth column) but for eight of these species some information was available on threats indicated by a double asterix (**) in the ninth and tenth columns accordingly (see also footnotes). Synonyms on the IUCN Red List are presented in parentheses in the second column.

| Common name | Scientific name | CITES Appendix | IUCN Red List Threat Category | Population size | Population trend | Intentional hunting or harvesting threat? (Yes/No) | Threat code(s) (timing) | Is trophy hunting a major contributor to elevated conservation concern (Yes/No) | Is trophy hunting likely or possibly causing localized declines (Likely/ Possibly /No) |
|---|---|---|---|---|---|---|---|---|---|
| Bongo | *Tragelaphus eurycerus* | NC | NT | 28,000 | Decreasing | Y | 5.1.1. (ongoing) | N | L |
| Brown Bear | *Ursus arctos* | I/II | LC | >200,000 | Stable | Y | 5.1.1. (ongoing) | N | L |
| Gobi Argali | *Ovis darwini* (*Ovis ammon darwini*) | II | NE | 12,100-14,800 | Unknown | NE | | N** | L** |
| Siberian Ibex | *Capra sibirica* | III | NT | 170,000-250,000 | Decreasing | Y | 5.1.1. (ongoing) | N | L |
| American Black Bear | *Ursus americanus* | II | LC | 850,000-950,000 | Increasing | Y | 5.1.1. | N | P |
| Puma | *Puma concolor* | I/II | LC | Unknown† | Decreasing | Y | 5.1.1. (ongoing) | N | P |
| Lion | *Panthera leo* | I/II | VU | 22,000-25,000 | Decreasing | Y | 5.1.1. (ongoing) | N | P |
| Leopard | *Panthera pardus* | I | VU | 231,636 | Decreasing | Y | 5.1.1. (ongoing) | N | P |
| Northern Goshawk | *Accipiter gentilis* | II | LC | 900,000-1,460,000 | Unknown | Y | 5.1.1. (ongoing) | N | N |
| Black-mantled Goshawk | *Accipiter melanochlamys* | II | LC | Unknown | Stable | N | | N | N |
| Black Sparrowhawk | *Accipiter melanoleucus* | II | LC | Unknown | Decreasing | N | | N | N |

| Common name | Scientific name | CITES Appendix | IUCN Red List Threat Category | Population size | Population trend | Intentional hunting or harvesting threat? (Yes/No) | Threat code(s) (timing) | Is trophy hunting a major contributor to elevated conservation concern (Yes/No) | Is trophy hunting likely or possibly causing localized declines (Likely/ Possibly /No) |
|---|---|---|---|---|---|---|---|---|---|
| Sharp-shinned Hawk | *Accipiter striatus* | II | LC | 1,000,000‡ | Decreasing | N | | N | N |
| Cheetah | *Acinonyx jubatus* | I | VU | 7100 | Decreasing | Y | 5.1.1. (ongoing) | N | N |
| Addax* | *Addax nasomaculatus* | I | CR | <100 | Decreasing | Y | 5.1.1. (ongoing) | N | N |
| Cinereous Vulture | *Aegypius monachus* | II | NT | 25,200-34,200 | Decreasing | Y | 5.1.1. (ongoing) | N | N |
| Southern Black Bustard | *Afrotis afra* | II | VU | Unknown | Decreasing | N | | N | N |
| Northern Black Bustard | *Afrotis afraoides* | II | LC | Unknown (but generally common) | Stable | N | | N | N |
| American Alligator | *Alligator mississippiensis* | II | LC | 3,000,000-4,000,000 | Increasing | N | | N | N |
| Egyptian Goose | *Alopochen aegyptiaca* | NC | LC | Unknown | Decreasing/stable | Y | | N | N |
| Aoudad* | *Ammotragus lervia* | II | VU | 5000-10,000‡ | Decreasing | Y | 5.1.1. (ongoing) | N | N |
| Northern Pintail | *Anas acuta* | NC | LC | 7,100,000-7,200,000 | Decreasing | Y | | N | N |
| Cape Teal | *Anas capensis* | NC | LC | 25,751-82,500 | Decreasing | N | 5.1.1. (ongoing) | N | N |
| Northern Shoveler | *Anas clypeata (Spatula clypeata)* | NC | LC | 6,500,000-7,000,000 | Decreasing | Y | 5.1.1. (ongoing) | N | N |
| Eurasian Teal | *Anas crecca* | NC | LC | 2,800,000‡ | Unknown | N | | N | N |
| Baikal Teal | *Anas formosa (Sibirionetta formosa)* | II | LC | 500,000-700,000 | Stable | Y | 5.1.1. (ongoing) | N | N |
| Campbell Teal | *Anas nesiotis* | I | VU | Unknown† | Stable | N | | N | N |
| Eurasian Wigeon | *Anas penelope* | NC | LC | 2,650,000-3,590,000 | Increasing | Y | | N | N |
| Garganey | *Anas querquedula (Spatula querquedula)* | NC | LC | 1,550,000-2,550,000 | Decreasing | Y | 5.1.1. (ongoing) | N | N |

| Common name | Scientific name | CITES Appendix | IUCN Red List Threat Category | Population size | Population trend | Intentional hunting or harvesting threat? (Yes/No) | Threat code(s) (timing) | Is trophy hunting a major contributor to elevated conservation concern (Yes/No) | Is trophy hunting likely or possibly causing localized declines (Likely/ Possibly /No) |
|---|---|---|---|---|---|---|---|---|---|
| Demoiselle Crane | *Anthropoides virgo* | II | LC | 230,000-261,000 | Increasing | Y | 5.1.1. (ongoing) | N | N |
| Sandhill Crane | *Antigone canadensis (Grus canadensis)* | II | LC | 670,000-830,000 | Increasing | N | | N | N |
| Pronghorn | *Antilocapra americana* | I (Mexico) | LC | ~1,000,000 | Stable | Y | 5.1.1. (ongoing) | N | N |
| Blackbuck* | *Antilope cervicapra* | III | LC | 35,200 (78,800 with introduced animals) | Unknown | Y | 5.1.1. (ongoing) | N | N |
| African Clawless Otter | *Aonyx capensis* | II | NT | Unknown† | Decreasing | Y | 5.1.1. (ongoing) | N | N |
| Smooth Softshell Turtle | *Apalone mutica* | II | LC | Unknown | Decreasing | Y | 5.4.1. (ongoing) | N | N |
| Golden Eagle | *Aquila chrysaetos* | II | LC | 85,000-160,000‡ | Stable | N | | N | N |
| Steppe Eagle | *Aquila nipalensis* | II | EN | 78,042-110,193 | Decreasing | Y | 5.1.1. (ongoing) | N | N |
| Tawny Eagle | *Aquila rapax* | II | VU | 100,000-999,999 | Decreasing | Y | 5.1.1. (ongoing) | N | N |
| Verreaux's Eagle | *Aquila verreauxii* | II | LC | Unknown but in the tens of thousands | Stable | N | | N | N |
| Red-and-green Macaw | *Ara chloropterus* | II | LC | 50,000-499,999‡ | Decreasing | N | | N | N |
| Afro-Australian Fur Seal | *Arctocephalus pusillus* | II | LC | 2,120,000 | Increasing | Y | 5.4.2. (ongoing) | N | N |
| Arabian Bustard | *Ardeotis arabs* | II | NT | Unknown | Decreasing | Y | 5.1.1. (ongoing) | N | N |
| Kori Bustard | *Ardeotis kori* | II | NT | Unknown | Decreasing | Y | 5.1.1. (ongoing) | N | N |
| Marsh Owl | *Asio capensis* | II | LC | Unknown (but locally common) | Stable | N | | N | N |
| Northern Long-eared Owl | *Asio otus* | II | LC | 2,230,000-3,680,000‡ | Decreasing | N | | N | N |
| Burrowing Owl | *Athene cunicularia* | II | LC | 1,000,000-9,999,999 | Stable | N | | N | N |

| Common name | Scientific name | CITES Appendix | IUCN Red List Threat Category | Population size | Population trend | Intentional hunting or harvesting threat? (Yes/No) | Threat code(s) (timing) | Is trophy hunting a major contributor to elevated conservation concern (Yes/No) | Is trophy hunting likely or possibly causing localized declines (Likely/ Possibly /No) |
|---|---|---|---|---|---|---|---|---|---|
| Little Owl | *Athene noctua* | II | LC | 4,960,000-9,360,000‡ | Stable | N | | N | N |
| Hog Deer | *Axis porcinus* | III | EN | Unknown | Decreasing | Y | 5.1.1. (ongoing) | N | N |
| Grey Crowned Crane | *Balearica regulorum* | II | EN | 30,200-36,900 | Decreasing | Y | 5.1.1. (ongoing) | N | N |
| American Bison | *Bison bison* | NC | NT | 18,748 | Stable | Y | 5.1.1. (past, unlikely to return) | N | N |
| Wood Bison | *Bison bison athabascae* | NC | NE | 11,000 | Stable | NE | | N** | N** |
| Wild Yak* | *Bos mutus* | I | VU | 10,000‡ | Decreasing | Y | 5.1.1. (ongoing) | N | N |
| Nilgai | *Boselaphus tragocamelus* | III | LC | 100,000 | Stable | Y | | N | N |
| Hadada Ibis | *Bostrychia hagedash* | NC | LC | 70,000-400,000 | Unknown | Y | 5.1.1. (ongoing) | N | N |
| Aleutian Canada Goose | *Branta canadensis leucopareia* | II | NE | 162,000§ | Unknown | NE | | NE | N |
| Wild Water Buffalo | *Bubalus arnee* | III | EN | <4000 | Decreasing | Y | 5.1.1. (ongoing) | N | N |
| Eurasian Eagle-owl | *Bubo bubo* | II | LC | 180,000-300,000 | Decreasing | N | | N | N |
| Cape Eagle-owl | *Bubo capensis* | II | LC | Unknown | Stable | N | | N | N |
| Great Horned Owl | *Bubo virginianus* | II | LC | Unknown | Stable | N | | N | N |
| Takin | *Budorcas taxicolor* | II | VU | ~10,000 | Decreasing | Y | 5.1.1. (ongoing) | N | N |
| Grasshopper Buzzard | *Butastur rufipennis* | II | LC | 20,000-49,999‡ | Decreasing | Y | 5.1.1. (ongoing) | N | N |
| Augur Buzzard | *Buteo augur* | II | LC | Unknown | Decreasing | N | | N | N |
| Eurasian Buzzard | *Buteo buteo* | II | LC | 2,038,000-3,463,000‡ | Increasing | Y | 5.1.1. (ongoing) | N | N |
| Red-tailed Hawk | *Buteo jamaicensis* | II | LC | 3,100,000‡ | Increasing | N | | N | N |

| Common name | Scientific name | CITES Appendix | IUCN Red List Threat Category | Population size | Population trend | Intentional hunting or harvesting threat? (Yes/No) | Threat code(s) (timing) | Is trophy hunting a major contributor to elevated conservation concern (Yes/No) | Is trophy hunting likely or possibly causing localized declines (Likely/ Possibly /No) |
|---|---|---|---|---|---|---|---|---|---|
| Long-legged Buzzard | *Buteo rufinus* | II | LC | 162,000-269,000‡ | Stable | Y | 5.1.1. (ongoing) | N | N |
| Jackal Buzzard | *Buteo rufofuscus* | II | LC | Unknown but in the tens of thousands | Stable | N | | N | N |
| Sulphur-crested Cockatoo | *Cacatua galerita* | II | LC | Unknown | Decreasing | N | | N | N |
| Spectacled Caiman | *Caiman crocodilus* | II | LC | 1,000,000‡ | Stable | Y | 5.4.2. (ongoing) | N | N |
| South American Spectacled Caiman | *Caiman crocodilus crocodilus* | II | NE | Unknown | Unknown | NE | | NE | N |
| Brown Spectacled Caiman | *Caiman crocodilus fuscus* | II | NE | Unknown | Unknown | NE | | NE | N |
| Muscovy Duck | *Cairina moschata* | NC | LC | 50,000-499,999 | Decreasing | N | | N | N |
| Common Jackal | *Canis aureus* | III | LC | 150,000† | Increasing | N | | N | N |
| Grey Wolf | *Canis lupus* | I/II | LC | ~130,000¶ | Stable | N | | N | N |
| Northern Rocky Mountain Wolf | *Canis lupus irremotus* | II | NE | Unknown | Unknown | NE | | NE | N |
| Texas Grey Wolf | *Canis lupus monstrabilis* | II | NE | Unknown | Unknown | NE | | NE | N |
| Western Tur | *Capra caucasica* | II | EN | 4000-5000 | Decreasing | Y | 5.1.1. (ongoing) | N | N |
| Markhor | *Capra falconeri heptneri* | I | NE | 1680 | Unknown | NE | | N** | N** |
| Straight-horned Markhor | *Capra falconeri megaceros* | I | NE | ~3000 | Unknown | NE | | N** | N** |
| Markhor | *Capra falconeri***** | I | NT | 9700 | Increasing | Y | 5.1.1. (ongoing) | N | N |
| Goat | *Capra hircus* | III (subsp. *Capra hircus aegagrus*) | NE | Unknown | Unknown | NE | | NE | N |
| Wild Goat | *Capra hircus aegagrus (Capra aegagrus)* | III | NT | 224,400 | Stable | Y | 5.1.1. (ongoing) | N | N |

| Common name | Scientific name | CITES Appendix | IUCN Red List Threat Category | Population size | Population trend | Intentional hunting or harvesting threat? (Yes/No) | Threat code(s) (timing) | Is trophy hunting a major contributor to elevated conservation concern (Yes/No) | Is trophy hunting likely or possibly causing localized declines (Likely/ Possibly /No) |
|---|---|---|---|---|---|---|---|---|---|
| Mainland Serow | *Capricornis sumatraensis* | I | VU | Unknown | Decreasing | Y | 5.1.1. (ongoing) | N | N |
| African Golden Cat | *Caracal aurata* | II | VU | Unknown (but rare) | Decreasing | N | | N | N |
| Caracal | *Caracal caracal* | I/II | LC | Unknown | Unknown | N | | N | N |
| Keel-scaled Boa | *Casarea dussumieri* | I | VU | 1818‡ | Increasing | N | | N | N |
| African Spurred Tortoise | *Centrochelys sulcata* | II | EN | Unknown | Decreasing | Y | 5.1.1. (ongoing) | N | N |
| Bay Duiker | *Cephalophus dorsalis* | II | NT | 725,000 | Decreasing | Y | 5.1.1. (ongoing) | N | N |
| Ogilby's Duiker | *Cephalophus ogilbyi* | II | LC | 35,000 | Decreasing | Y | 5.1.1. (ongoing) | N | N |
| Yellow-backed Duiker | *Cephalophus silvicultor* | II | NT | 160,000 | Decreasing | Y | 5.1.1. (ongoing) | N | N |
| Zebra Duiker | *Cephalophus zebra* | II | VU | 15,000 | Decreasing | Y | 5.1.1. (ongoing) | N | N |
| White Rhinoceros | *Ceratotherium simum* | I/II | NT | 18,064 | Decreasing | Y | 5.1.1. (ongoing) | N | N |
| Southern White Rhinoceros | *Ceratotherium simum simum* | I/II | NT | 18,064 | Decreasing | Y | 5.1.1. (ongoing) | N | N |
| South Asian Bockadam | *Cerberus rynchops* | III | LC | 374-1,396 | Unknown | Y | 5.4.1. (ongoing), 5.4.2. (ongoing) | N | N |
| Agile Mangabey | *Cercocebus agilis* | II | LC | Unknown | Decreasing | Y | 5.1.1. (ongoing) | N | N |
| Tana River Mangabey | *Cercocebus galeritus* | I | CR | 1000 | Decreasing | N | | N | N |
| Sanje River Mangabey | *Cercocebus sanjei* | II | EN | 1300-3500 | Decreasing | Y | 5.1.1. (ongoing) | N | N |
| Red-tailed Monkey | *Cercopithecus ascanius* | II | LC | Unknown | Decreasing | Y | 5.1.1. (ongoing) | N | N |
| Moustached Monkey | *Cercopithecus cephus* | II | LC | Unknown | Unknown | Y | 5.1.1. (ongoing) | N | N |
| Blue Monkey | *Cercopithecus mitis* | II | LC | Unknown | Decreasing | Y | 5.1.1. (ongoing) | N | N |

| Common name | Scientific name | CITES Appendix | IUCN Red List Threat Category | Population size | Population trend | Intentional hunting or harvesting threat? (Yes/No) | Threat code(s) (timing) | Is trophy hunting a major contributor to elevated conservation concern (Yes/No) | Is trophy hunting likely or possibly causing localized declines (Likely/ Possibly /No) |
|---|---|---|---|---|---|---|---|---|---|
| Mona Monkey | *Cercopithecus mona* | II | NT | Unknown | Decreasing | Y | 5.1.1. (ongoing) | N | N |
| De Brazza's Monkey | *Cercopithecus neglectus* | II | LC | Unknown | Unknown | Y | 5.1.1. (ongoing) | N | N |
| Putty-nosed Monkey | *Cercopithecus nictitans* | II | NT | Unknown† | Decreasing | Y | 5.1.1. (ongoing) | N | N |
| Spot-nosed Monkey | *Cercopithecus petaurista* | II | NT | Unknown† | Decreasing | Y | 5.1.1. (ongoing) | N | N |
| Crowned Monkey | *Cercopithecus pogonias* | II | NT | Unknown | Decreasing | Y | 5.1.1. (ongoing) | N | N |
| Red Deer | *Cervus elaphus* | NC (subsp. on I/II/III) | LC | ≥2,400,000 | Increasing | Y | 5.1.1. (past, unlikely to return) | N | N |
| Bactrian Red Deer | *Cervus elaphus bactrianus (C. hanglu bactrianus)* | II | LC | 2000-2500 | Increasing | Y | 5.1.1. (ongoing) | N | N |
| Green Turtle | *Chelonia mydas* | I | LC | Unknown | Increasing | Y | 5.1.1. (ongoing) 5.4.1. (ongoing) | N | N |
| Snapping Turtle | *Chelydra serpentina* | II | LC | Unknown | Decreasing | Y | 5.1.1. (ongoing) 5.4.1. (ongoing) | N | N |
| Grivet Monkey | *Chlorocebus aethiops* | II | LC | Unknown | Decreasing | N | | N | N |
| Vervet Monkey | *Chlorocebus pygerythrus* | II | LC | Unknown (but locally abundant) | Decreasing | Y | 5.1.1. (ongoing) | N | N |
| Green Monkey | *Chlorocebus sabaeus* | II | LC | At least 40,000 | Decreasing | Y | 5.1.1. (ongoing) | N | N |
| Tantalus Monkey | *Chlorocebus tantalus* | II | LC | Unknown (but common) | Stable | N | | N | N |
| Black Stork | *Ciconia nigra* | II | LC | 20,200-32,400 | Unknown | N | | N | N |
| Brown Snake-eagle | *Circaetus cinereus* | II | LC | 6700-67,000‡ | Decreasing | Y | 5.1.1. (ongoing) | N | N |
| Southern Banded Snake-eagle | *Circaetus fasciolatus* | II | NT | 1000-3000 | Decreasing | N | | N | N |

| Common name | Scientific name | CITES Appendix | IUCN Red List Threat Category | Population size | Population trend | Intentional hunting or harvesting threat? (Yes/No) | Threat code(s) (timing) | Is trophy hunting a major contributor to elevated conservation concern (Yes/No) | Is trophy hunting likely or possibly causing localized declines (Likely/ Possibly /No) |
|---|---|---|---|---|---|---|---|---|---|
| Black-chested Snake-eagle | *Circaetus pectoralis* | II | LC | Unknown | Stable | N | | N | N |
| Western Marsh-harrier | *Circus aeruginosus* | II | LC | 631,000-1,010,000‡ | Stable | N | | N | N |
| African Civet | *Civettictis civetta* | III | LC | Unknown (but generally common) | Unknown | Y | 5.1.1. (ongoing) | N | N |
| Brown Violetear | *Colibri delphinae* | II | LC | 500,000-4,999,999 | Decreasing | N | | N | N |
| Guereza | *Colobus guereza* | II | LC | Unknown (widespread and locally abundant) | Decreasing | Y | 5.1.1. (ongoing) | N | N |
| King Colobus | *Colobus polykomos* | II | EN | Unknown | Decreasing | Y | 5.1.1. (ongoing) | N | N |
| Black Colobus | *Colobus satanas* | II | VU | Unknown† | Decreasing | Y | 5.1.1. (ongoing) | N | N |
| Speckled Pigeon | *Columba guinea* | NC | LC | Unknown (but common) | Stable | N | | N | N |
| Rock Dove | *Columba livia* | NC | LC | 140,000,000 | Unknown | N | | N | N |
| Great Curassow | *Crax rubra* | III | VU | 50,000-499,999 | Decreasing | Y | 5.1.1. (ongoing) | N | N |
| Slender-snouted Crocodile | *Crocodylus cataphractus* (*Mecistops cataphractus*) | I | CR | 1000-20,000‡ | Decreasing | Y | 5.1.1. (ongoing) | N | N |
| Nile Crocodile | *Crocodylus niloticus* | I/II | LC | 50,000-70,000‡ | Stable | Y | 5.4.1., 5.4.2. | N | N |
| Saltwater Crocodile | *Crocodylus porosus* | I/II | LC | 500,000 | Stable | N | | N | N |
| Cascabel Rattlesnake | *Crotalus durissus* | III | LC | Unknown (but common) | Unknown | N | | N | N |
| Paca | *Cuniculus paca* | III | LC | Unknown† | Stable | Y | 5.1.1. (ongoing) | N | N |
| Topi | *Damaliscus lunatus* | III | LC | Unknown† | Decreasing | Y | 5.1.1. (ongoing) | N | N |

| **Common name** | **Scientific name** | **CITES Appendix** | **IUCN Red List Threat Category** | **Population size** | **Population trend** | **Intentional hunting or harvesting threat? (Yes/No)** | **Threat code(s) (timing)** | **Is trophy hunting a major contributor to elevated conservatio n concern (Yes/No)** | **Is trophy hunting likely or possibly causing localized declines (Likely/ Possibly /No)** |
|---|---|---|---|---|---|---|---|---|---|
| Blesbok | *Damaliscus pygargus* | II (subsp. *pygargus*) | LC | 77,751 | Increasing | N | | N | N |
| Bontebok* | *Damaliscus pygargus pygargus* | II | VU | 515-1618‡ | Stable | Y | 5.1.1. (ongoing) | N | N |
| Central American Agouti | *Dasyprocta punctata* | III | LC | Unknown (but common) | Stable | Y | 5.1.1. (ongoing) | N | N |
| West Indian Whistling-duck | *Dendrocygna arborea* | II | NT | 6000-15000‡ | Decreasing | Y | 5.1.1. (ongoing) | N | N |
| Black-bellied Whistling-duck | Dendrocygna autumnalis | III | LC | 200,000-2,000,000‡ | Increasing | N | | N | N |
| Fulvous Whistling-duck | *Dendrocygna bicolor* | III | LC | 1,230,000-1,470,001 | Decreasing | Y | 5.1.1. (ongoing) | N | N |
| White-faced Whistling duck | *Dendrocygna viduata* | NC | LC | 2,120,000-2,500,001 | Increasing | Y | | N | N |
| Black Rhinoceros | *Diceros bicornis* | I/II | CR | 5630 | Increasing | Y | 5.1.1. (ongoing) | N | N |
| Little Egret | *Egretta garzetta* | NC | LC | 645,000-3,145,061 | Stable | N | | N | N |
| Black-winged Kite | *Elanus caeruleus* | II | LC | Unknown (but common) | Stable | Y | 5.1.1. (ongoing) | N | N |
| White-tailed Kite | *Elanus leucurus* | II | LC | 260,000 | Decreasing | Y | 5.1.1. (ongoing) | N | N |
| Copper-head Trinket Snake | *Elaphe radiata (Coelognathus radiatus)* | NC | LC | Unknown (but common) | Unknown | Y | 5.1.1. (ongoing) | N | N |
| Sea Otter | *Enhydra lutris* | II | EN | 128,902 | Decreasing | Y | 5.1.1. (ongoing) | N | N |
| Saddlebill | *Ephippiorhynchus senegalensis* | NC | NT | 1000-25,000 | Decreasing | Y | 5.1.1. (ongoing) | N | N |
| Grevy's Zebra | *Equus grevyi* | II | EN | 2680 | Stable | Y | 5.1.1. (ongoing) | N | N |
| Mountain Zebra | *Equus zebra* | NC (subsp. in II) | VU | 34,979‡ | Increasing | N | | N | N |

| Common name | Scientific name | CITES Appendix | IUCN Red List Threat Category | Population size | Population trend | Intentional hunting or harvesting threat? (Yes/No) | Threat code(s) (timing) | Is trophy hunting a major contributor to elevated conservatio n concern (Yes/No) | Is trophy hunting likely or possibly causing localized declines (Likely/ Possibly /No) |
|---|---|---|---|---|---|---|---|---|---|
| Hartmann's Mountain Zebra | *Equus zebra hartmannae* | II | VU | 33,265 | Increasing | Y | 5.1.1. (ongoing) | N | N |
| Cape Mountain Zebra | *Equus zebra zebra* | II | LC | 3247 | Increasing | Y | 5.1.1. (past) | N | N |
| Patas Monkey | *Erythrocebus patas* | II | NT | Unknown† | Decreasing | N | | N | N |
| Purple-throated Carib | *Eulampis jugularis* | II | LC | Unknown (but common) | Decreasing | N | | N | N |
| Green Anaconda | *Eunectes murinus* | II | LC | Unknown | Unknown | Y | 5.1.1. (ongoing) | N | N |
| White-bellied Bustard | *Eupodotis senegalensis* | II | LC | Unknown | Decreasing | N | | N | N |
| Karoo Bustard | *Eupodotis vigorsii (Heterotetrax vigorsii)* | II | LC | Unknown (but generally common) | Increasing | N | | N | N |
| Saker Falcon | *Falco cherrug* | II | EN | 12,200-29,800‡ | Decreasing | Y | 5.1.1. (ongoing) | N | N |
| Red-headed Falcon | *Falco chicquera* | II | NT | 25,000-99,000 | Decreasing | Y | 5.1.1. (ongoing) | N | N |
| Lesser Kestrel | *Falco naumanni* | II | LC | 120,000-200,000 | Stable | Y | 5.1.1. (ongoing) | N | N |
| Peregrine Falcon | *Falco peregrinus* | I | LC | 248,000-478,000‡ | Increasing | Y | 5.1.1. (ongoing) | N | N |
| American Kestrel | *Falco sparverius* | II | LC | 9,200,000‡ | Decreasing | N | | N | N |
| Eurasian Hobby | *Falco Subbuteo* | II | LC | 933,000-1,457,000‡ | Decreasing | Y | 5.1.1. (ongoing) | N | N |
| Common Kestel | *Falco tinnunculus* | II | LC | 4,330,000-6,680,000‡ | Decreasing | Y | 5.1.1. (ongoing) | N | N |
| Jungle Cat | *Felis chaus* | II | LC | Unknown | Decreasing | Y | 5.1.1. (ongoing) | N | N |
| Afro-Asiatic Wildcat | *Felis lybica* | II | LC | Unknown | Unknown | Y | 5.1.1. (ongoing) | N | N |
| Black-footed Cat | *Felis nigripes* | I | VU | 13,867‡ | Decreasing | Y | 5.1.1. (ongoing) | N | N |
| European Wildcat | *Felis silvestris* | II | LC | 140,000 | Unknown | Y | 5.1.1. (ongoing) | N | N |

| Common name | Scientific name | CITES Appendix | IUCN Red List Threat Category | Population size | Population trend | Intentional hunting or harvesting threat? (Yes/No) | Threat code(s) (timing) | Is trophy hunting a major contributor to elevated conservatio n concern (Yes/No) | Is trophy hunting likely or possibly causing localized declines (Likely/ Possibly /No) |
|---|---|---|---|---|---|---|---|---|---|
| Southern Lesser Galago | *Galago moholi* | II | LC | Unknown (widespread and common) | Stable | N | | N | N |
| Northern Lesser Galago | *Galago senegalensis* | II | LC | Unknown† | Decreasing | N | | N | N |
| Grey Jungefowl | *Gallus sonneratii* | II | LC | Unknown (but locally common) | Decreasing | N | | N | N |
| Chinkara | *Gazella bennettii* | III | LC | 50,000-70,000‡ | Decreasing | Y | 5.1.1. (ongoing) | N | N |
| Dorcas Gazelle | *Gazella dorcas* | III | VU | 35,000-40,000 | Decreasing | Y | 5.1.1. (ongoing) | N | N |
| Giraffe | *Giraffa camelopardalis* | II | VU | 95,762 | Decreasing | Y | 5.1.1. (ongoing) | N | N |
| African Barred Owlet | *Glaucidium capense* | II | LC | Unknown | Decreasing | N | | N | N |
| Pearl-spotted Owlet | *Glaucidium perlatum* | II | LC | Unknown | Decreasing | N | | N | N |
| Bearded Vulture | *Gypaetus barbatus* | II | NT | 2500-10,000 | Decreasing | Y | 5.1.1. (ongoing) | N | N |
| Palm-nut Vulture | *Gypohierax angolensis* | II | LC | Unknown | Decreasing | Y | 5.1.1. (ongoing) | N | N |
| White-backed Vulture | *Gyps africanus* | II | CR | 27,000-270,000 | Decreasing | Y | 5.1.1. (ongoing) | N | N |
| Rüppell's Vulture | *Gyps rueppellii* | II | CR | <30,000 | Decreasing | Y | 5.1.1. (ongoing) | N | N |
| Bald Eagle | *Haliaeetus leucocephalus* | II | LC | 200,000‡ | Increasing | N | | N | N |
| Pallas's Fish-eagle | *Haliaeetus leucoryphus* | II | EN | <2,500 | Decreasing | Y | 5.1.1. (ongoing) | N | N |
| African Fish-eagle | *Haliaeetus vocifer* | II | LC | Unknown | Stable | N | | N | N |
| Jaguarundi | *Herpailurus yagouaroundi* | I/II | LC | Unknown (but uncommon) | Decreasing | N | | N | N |
| Common Hippopotamus | *Hippopotamus amphibius* | II | VU | 115,000-130,000 | Stable | Y | 5.1.1. (ongoing) | N | N |
| Roan Antelope | *Hippotragus equinus* | NC | LC | 50,000-60,000‡ | Decreasing | Y | 5.1.1. (ongoing) | N | N |

| Common name | Scientific name | CITES Appendix | IUCN Red List Threat Category | Population size | Population trend | Intentional hunting or harvesting threat? (Yes/No) | Threat code(s) (timing) | Is trophy hunting a major contributor to elevated conservation concern (Yes/No) | Is trophy hunting likely or possibly causing localized declines (Likely/ Possibly /No) |
|---|---|---|---|---|---|---|---|---|---|
| Sable Antelope* | *Hippotragus niger* | NC (subsp. in I) | LC | 50,000-60,000‡ | Stable | Y | 5.1.1. (ongoing) | N | N |
| Giant Sable* | *Hippotragus niger variani* | I | CR | 70-100‡ | Decreasing | Y | | N | N |
| Brown Hyaena | *Hyaena brunnea* | NC | NT | 5000-8000 | Stable | N | | N | N |
| Striped Hyena | *Hyaena hyaena* | III | NT | 5000-14,000 | Decreasing | Y | 5.1.1. (ongoing) | N | N |
| Spotted-necked Otter | *Hydrictis maculicollis* | II | NT | Unknown | Decreasing | Y | 5.1.1. (ongoing) | N | N |
| Water Chevrotain | *Hyemoschus aquaticus* | NC | LC | 278,000 | Decreasing | Y | 5.1.1. (ongoing) | N | N |
| Crested Porcupine | *Hystrix cristata* | NC | LC | Unknown | Unknown | N | | N | N |
| Blood Pheasant | *Ithaginis cruentus* | II | LC | Unknown | Decreasing | N | | N | N |
| Lizard Buzzard | *Kaupifalco monogrammicus* | II | LC | Unknown | Stable | N | | N | N |
| KwaZulu-Natal Hinged-back Tortoise | *Kinixys natalensis* | II | VU | Unknown | Decreasing | Y | | N | N |
| Lechwe | *Kobus leche* | II | NT | 212,000 | Decreasing | Y | 5.1.1. (ongoing) | N | N |
| Serval | *Leptailurus serval* | II | LC | Unknown | Stable | Y | 5.1.1. (ongoing) | N | N |
| Black-bellied Bustard | *Lissotis melanogaster* | II | LC | Unknown (but common-frequent) | Decreasing | N | | N | N |
| North American River Otter | *Lontra canadensis* | II | LC | Unknown | Stable | Y | 5.1.1. (ongoing) | N | N |
| Long-crested Eagle | *Lophaetus occipitalis* | II | LC | Unknown | Stable | N | 5.1.1. (ongoing) | N | N |
| Grey-cheeked Mangabey | *Lophocebus albigena* | II | VU | Unknown | Decreasing | Y | 5.1.1. (ongoing) | N | N |
| African Savanna Elephant | *Loxodonta africana* | I/II | EN | 415,428# | Decreasing | Y | 5.1.1. (ongoing) | N | N |
| African Forest Elephant | *Loxodonta cyclotis* | I/II | CR | 415,428# | Decreasing | Y | 5.1.1. (ongoing) | N | N |

| Common name | Scientific name | CITES Appendix | IUCN Red List Threat Category | Population size | Population trend | Intentional hunting or harvesting threat? (Yes/No) | Threat code(s) (timing) | Is trophy hunting a major contributor to elevated conservatio n concern (Yes/No) | Is trophy hunting likely or possibly causing localized declines (Likely/ Possibly /No) |
|---|---|---|---|---|---|---|---|---|---|
| Eurasian Otter | *Lutra lutra* | I | NT | 57,880-361,140‡ | Decreasing | Y | 5.1.1. (ongoing) | N | N |
| Culpeo | *Lycalopex culpaeus* | II | LC | Unknown† | Stable | Y | 5.1.1. (ongoing) | N | N |
| Chilla | *Lycalopex griseus* | II | LC | Unknown | Stable | Y | 5.1.1. (ongoing) | N | N |
| Pampas Fox | *Lycalopex gymnocercus* | II | LC | Unknown | Stable | Y | 5.1.1. (past, unlikely to return) | N | N |
| Canada Lynx | *Lynx canadensis* | II | LC | Unknown | Unknown | Y | 5.1.1. (ongoing) | N | N |
| Eurasian lynx | *Lynx lynx* | II | LC | Unknown† | Stable | Y | 5.1.1. (ongoing) | N | N |
| Bobcat | *Lynx rufus* | II | LC | 2,352,276-3,571,681 | Stable | Y | 5.1.1. (ongoing) | N | N |
| Long-tailed Macaque | *Macaca fascicularis* | II | EN | Unknown† | Decreasing | Y | 5.1.1. (ongoing) | N | N |
| Alligator Snapping Turtle | *Macrochelys temminckii* | II | EN | 68,154-1,436,825 adults | Decreasing | N | 5.4.1. (ongoing), 5.4.2. (past) | N | N |
| Reticulated Python | *Malayopython reticulatus* | II | LC | Unknown | Unknown | Y | 5.1.1. (ongoing) | N | N |
| Giant Pangolin | *Manis gigantea* | I | EN | Unknown | Decreasing | Y | 5.1.1. (ongoing) | N | N |
| White-bellied Pangolin | *Manis tricuspis* | I | EN | Unknown† | Decreasing | Y | 5.1.1. (ongoing) | N | N |
| Central American Red Brocket | *Mazama temama* | III | DD | Unknown | Decreasing | Y | 5.1.1. (ongoing) | N | N |
| Guatemalan Red Brocket | *Mazama temama cerasina* | III | DD | Unknown | Decreasing | Y | 5.1.1. (ongoing) | N | N |
| Ocellated Turkey | *Meleagris ocellata* | III | NT | 20,000-49,999 | Decreasing | Y | 5.1.1. (ongoing) | N | N |
| Pale-Chanting goshawk | *Melierax canorus* | II | LC | Unknown | Stable | N | | N | N |

| Common name | Scientific name | CITES Appendix | IUCN Red List Threat Category | Population size | Population trend | Intentional hunting or harvesting threat? (Yes/No) | Threat code(s) (timing) | Is trophy hunting a major contributor to elevated conservation concern (Yes/No) | Is trophy hunting likely or possibly causing localized declines (Likely/ Possibly /No) |
|---|---|---|---|---|---|---|---|---|---|
| Eastern Chanting-goshawk | *Melierax poliopterus* | II | LC | Unknown | Stable | N | | N | N |
| Honey Badger | *Mellivora capensis* | III | LC | Unknown | Decreasing | Y | 5.1.1. (ongoing) | N | N |
| Gabar Goshawk | *Micronisus gabar* | II | LC | Unknown | Stable | N | | N | N |
| Black Kite | *Milvus migrans* | II | LC | 4,100,000-5,600,000‡ | Stable | Y | 5.1.1. (ongoing) | N | N |
| Southern Talapoin Monkey | *Miopithecus talapoin* | II | VU | Unknown | Decreasing | Y | 5.1.1. (ongoing) | N | N |
| Siberian Musk Deer | *Moschus moschiferus* | II | VU | Unknown† | Decreasing | Y | 5.1.1. (ongoing) | N | N |
| Stoat | *Mustela erminea* | III | LC | Unknown | Stable | N | | N | N |
| Siberian Weasel | *Mustela sibirica* | III | LC | Unknown | Stable | Y | | N | N |
| Monk Parakeet | *Myiopsitta monachus* | II | LC | Unknown (but species is common) | Increasing | N | | N | N |
| Giant Anteater | *Myrmecophaga tridactyla* | II | VU | Unknown | Decreasing | Y | 5.1.1. (ongoing) | N | N |
| Himalayan Goral | *Naemorhedus goral* | I | NT | Unknown | Decreasing | Y | 5.1.1. (ongoing) | N | N |
| Dama Gazelle* | *Nanger dama* | I | CR | <100-250 | Decreasing | Y | 5.1.1. (ongoing) | N | N |
| White-nosed Coati | *Nasua narica* | III | LC | Unknown (rare to common) | Decreasing | Y | 5.1.1. (ongoing) | N | N |
| Hooded Vulture | *Necrosyrtes monachus* | II | CR | 197,000 | Decreasing | Y | 5.1.1. (ongoing) | N | N |
| Denham's Bustard | *Neotis denhami* | II | NT | Unknown | Decreasing | Y | 5.1.1. (ongoing) | N | N |
| African Pygmy-goose | *Nettapus auritus* | NC | LC | 57,000-320,000 | Stable | Y | | N | N |
| Snowy Owl | *Nyctea scandiaca (Bubo scandiacus)* | II | VU | 14,000-28,000‡ | Decreasing | Y | 5.1.1. (ongoing) | N | N |

| Common name | Scientific name | CITES Appendix | IUCN Red List Threat Category | Population size | Population trend | Intentional hunting or harvesting threat? (Yes/No) | Threat code(s) (timing) | Is trophy hunting a major contributor to elevated conservation concern (Yes/No) | Is trophy hunting likely or possibly causing localized declines (Likely/ Possibly /No) |
|---|---|---|---|---|---|---|---|---|---|
| Walrus | *Odobenus rosmarus* | III | VU | >245,000 | Unknown | Y | 5.4.1. (ongoing), 5.4.2. (past) | N | N |
| White-tailed Deer | *Odocoileus virginianus* | NC (subsp. on III) | LC | 11,500,000 | Stable | Y | 5.1.1. (ongoing) | N | N |
| Guatemalan White-tailed Deer | *Odocoileus virginianus mayensis* | III | NE | Unknown | Unknown | NE | | NE | N |
| Namaqua Dove | *Oena capensis* | NC | LC | Unknown (common to abundant) | Increasing | Y | | N | N |
| Plain Chachalaca | *Ortalis vetula* | III | LC | 2,000,000‡ | Stable | N | | N | N |
| Aardvark | *Orycteropus afer* | NC | LC | Unknown | Unknown | Y | 5.1.1. (ongoing) | N | N |
| Scimitar-horned Oryx* | *Oryx dammah* | I | EN | 427-1652 | Increasing | Y | 5.1.1. (past, likely to return) | N | N |
| Arabian Oryx* | *Oryx leucoryx* | I | VU | 1220 reintroduced (plus 6000 ex situ) | Stable | Y | 5.1.1. (ongoing) | N | N |
| Great Bustard | *Otis tarda* | II | EN | 29,600-33,000‡ | Decreasing | Y | 5.1.1. (ongoing) | N | N |
| Manul | *Otocolobus manul* | II | LC | 49,000–98,000‡ | Decreasing | Y | 5.1.1. (ongoing) | N | N |
| Thick-tailed Greater Galago | *Otolemur crassicaudatus* | II | LC | Unknown (relatively common) | Stable | N | | N | N |
| Wild Sheep | *Ovis ammon* | II | NT | <107,000 | Decreasing | Y | 5.1.1. (ongoing) | N | N |
| Urial | *Ovis aries* | NC | NE | Unknown | Unknown | NE | | NE | N |
| Bukhara Urial | *Ovis bochariensis (Ovis v. bochariensis)* | II | EN | <800 | Decreasing | Y | | N | N |
| Bighorn Sheep | *Ovis canadensis* | II | LC | 70,000 | Stable | Y | 5.1.1. (ongoing) | N | N |
| Afghan Urial | *Ovis cycloceros (O. v. cycloceros)* | II | VU | 20,000-30,000 | Decreasing | Y | | N | N |

| Common name | Scientific name | CITES Appendix | IUCN Red List Threat Category | Population size | Population trend | Intentional hunting or harvesting threat? (Yes/No) | Threat code(s) (timing) | Is trophy hunting a major contributor to elevated conservation concern (Yes/No) | Is trophy hunting likely or possibly causing localized declines (Likely/ Possibly /No) |
|---|---|---|---|---|---|---|---|---|---|
| Afghan Urial (subsp. arkal) | *Ovis cycloceros arkal (O.v. arkal)* | II | CR | 1500-1850 | Decreasing | Y | | N | N |
| Afghan Urial (subsp. cycloceros) | *Ovis cycloceros cycloceros (O.v. cycloceros)* | II | NE | Unknown | Unknown | NE | | N** | N** |
| Mouflon | *Ovis gmelini* | I (Cyprus population) | NT | 26,500 | Unknown | Y | 5.1.1. (ongoing) | N | N |
| Tibetan Argali | *Ovis hodgsoni (O. a. hodgonsi)* | I | NE | 29,000-36,000 | Unknown | NE | | N** | N** |
| Tianshan Argali | *Ovis karelini (O.a. karelini)* | II | NE | 22,371-25,371 | Unknown | NE | | N** | N** |
| Karatau Argali | *Ovis nigrimontana (O.a. nigrimontana)* | I | EN | <250 | Decreasing | Y | | N | N |
| Marco Polo Argali | *Ovis polii* | II | NE | 4000-7000 (see *Ovis ammon*) | Unknown | NE | | N** | N** |
| Punjab Urial | *Ovis punjabiensis (Ovis v. punjabensis)* | II | VU | <800 | Decreasing | Y | | N | N |
| Severtzov's Argali | *Ovis severtzovi (O.a. severtzovi)* | II | VU | <2000 | Decreasing | Y | | N | N |
| Urial | *Ovis vignei* | I | VU | 18,000‡ | Decreasing | Y | 5.1.1. (ongoing) | N | N |
| Ruddy Duck | *Oxyura jamaicensis* | NC | LC | Unknown | Decreasing | N | | N | N |
| White-headed Duck | *Oxyura leucocephala* | II | EN | 5300-8700‡ | Decreasing | Y | 5.1.1. (ongoing) | N | N |
| Chimpanzee | *Pan troglodytes* | I | EN | 172,700–299,700 | Decreasing | Y | 5.1.1. (ongoing) | N | N |
| Chiru* | *Pantholops hodgsonii* | I | NT | 100,000-150,000 | Increasing | Y | 5.1.1. (ongoing) | N | N |
| Olive Baboon | *Papio anubis* | II | LC | Unknown (widespread and locally common) | Stable | Y | 5.1.1. (ongoing) | N | N |

| Common name | Scientific name | CITES Appendix | IUCN Red List Threat Category | Population size | Population trend | Intentional hunting or harvesting threat? (Yes/No) | Threat code(s) (timing) | Is trophy hunting a major contributor to elevated conservation concern (Yes/No) | Is trophy hunting likely or possibly causing localized declines (Likely/ Possibly /No) |
|---|---|---|---|---|---|---|---|---|---|
| Yellow Baboon | *Papio cynocephalus* | II | LC | Unknown (widespread and locally common) | Stable | N | | N | N |
| Hamadryas Baboon | *Papio hamadryas* | II | LC | 2000 | Increasing | Y | 5.1.1. (ongoing) | N | N |
| Guinea Baboon | *Papio papio* | II | NT | Unknown (but common in places) | Decreasing | Y | 5.1.1. (ongoing) | N | N |
| Chacma Baboon | *Papio ursinus* | II | LC | Unknown (common and widespread) | Decreasing | N | | N | N |
| Harris's Hawk | *Parabuteo unicinctus* | II | LC | 920,000‡ | Unknown | N | | N | N |
| Indian Peafowl | *Pavo cristatus* | III | LC | Unknown (common) | Increasing | N | | N | N |
| Collared Pecary | *Pecari tajacu* | II | LC | Unknown | Stable | Y | 5.1.1. (ongoing) | N | N |
| Crested Guan | *Penelope purpurascens* | III | NT | 50,000-499999‡ | Decreasing | Y | 5.1.1. (ongoing) | N | N |
| West African Potto | *Perodicticus potto* | II | NT | Unknown | Decreasing | Y | 5.1.1. (ongoing) | N | N |
| Maxwell's Duiker | *Philantomba maxwellii* | II | LC | 2,137,000 | Decreasing | Y | 5.1.1. (ongoing) | N | N |
| Blue Duiker | *Philantomba monticola* | II | LC | 7,000,000 | Decreasing | Y | 5.1.1. (ongoing) | N | N |
| Lesser Flamingo | *Phoeniconaias minor* | II | NT | 2,220,000-3,240,000 | Decreasing | N | | N | N |
| Greater Flamingo | *Phoenicopterus roseus* | II | LC | 550,000-680,000 | Increasing | Y | | N | N |
| American Flamingo | *Phoenicopterus ruber* | II | LC | 219,490-307,490 | Increasing | N | | N | N |
| Eastern Rosella | *Platycercus eximius* | II | LC | Unknown (extremely common) | Increasing | N | | N | N |
| Spur-winged Goose | *Plectropterus gambensis* | NC | LC | 300,000-500,000 | Stable | Y | | N | N |

| Common name | Scientific name | CITES Appendix | IUCN Red List Threat Category | Population size | Population trend | Intentional hunting or harvesting threat? (Yes/No) | Threat code(s) (timing) | Is trophy hunting a major contributor to elevated conservation concern (Yes/No) | Is trophy hunting likely or possibly causing localized declines (Likely/ Possibly /No) |
|---|---|---|---|---|---|---|---|---|---|
| Brown-headed Parrot | *Poicephalus cryptoxanthus* | II | LC | Unknown | Decreasing | Y | 5.1.1. (ongoing) | N | N |
| Brown-necked Parrot | *Poicephalus fuscicollis* | II | LC | Unknown | Decreasing | N | | N | N |
| Red-fronted Parrot | *Poicephalus gulielmi* | II | LC | Unknown | Decreasing | Y | | N | N |
| Martial Eagle | *Polemaetus bellicosus* | II | EN | Unknown† | Decreasing | Y | 5.1.1. (ongoing) | N | N |
| African Pygmy-falcon | *Polihierax semitorquatus* | II | LC | Unknown | Stable | N | | N | N |
| African Harrier-hawk | *Polyboroides typus* | II | LC | Unknown | Decreasing | N | | N | N |
| Aardwolf | *Proteles cristata* | III | LC | Unknown | Stable | Y | | N | N |
| Blue Sheep | *Pseudois nayaur* | III | LC | 47,000-414,000 | Unknown | Y | 5.1.1. (ongoing) | N | N |
| Grey Parrot | *Psittacus erithacus* | I | EN | 560,000-12,700,000 | Decreasing | Y | 5.1.1. (ongoing) | N | N |
| Hartlaub's Duck | *Pteronetta hartlaubii* | NC | LC | 25,001-110,000 | Decreasing | Y | 5.1.1. (ongoing) | N | N |
| Florida Puma | *Puma concolor couguar* | I/II | NE | Unknown† | Decreasing | NE | | NE | N |
| Brongersma's Short-tailed Python | *Python brongersmai* | II | LC | Unknown | Increasing | Y | 5.1.1. (ongoing) | N | N |
| Southern African Rock Python | *Python natalensis* | II | LC | Unknown | Decreasing | Y | 5.1.1. (ongoing) | N | N |
| Ball Python | *Python regius* | II | NT | Unknown | Decreasing | Y | 5.1.1. (ongoing) | N | N |
| Central African Rock Python | *Python sebae* | II | NT | Unknown | Decreasing | Y | 5.1.1. (ongoing) | N | N |
| Swamp Deer | *Rucervus duvaucelii* | I | VU | Unknown† | Decreasing | Y | 5.1.1. (ongoing) | N | N |
| Apennine Chamois | *Rupicapra pyrenaica ornata* | II | VU | 2,500 | Increasing | N | | N | N |
| Secretarybird | *Sagittarius serpentarius* | II | EN | 6700-67,000‡ | Decreasing | Y | 5.1.1. (ongoing) | N | N |

| Common name | Scientific name | CITES Appendix | IUCN Red List Threat Category | Population size | Population trend | Intentional hunting or harvesting threat? (Yes/No) | Threat code(s) (timing) | Is trophy hunting a major contributor to elevated conservation concern (Yes/No) | Is trophy hunting likely or possibly causing localized declines (Likely/ Possibly /No) |
|---|---|---|---|---|---|---|---|---|---|
| Golden-handed Tamarin | *Saguinus midas* | II | LC | Unknown† | Stable | Y | | N | N |
| Saiga* | *Saiga tatarica* | II | NT | 1,344,275 | Increasing | Y | 5.1.1. (ongoing) | N | N |
| Guianan Squirrel Monkey | *Saimiri sciureus* | II | LC | Unknown | Decreasing | N | | N | N |
| Red Tegu | *Salvator rufescens* | II | LC | Unknown | Unknown | Y | 5.1.1. (ongoing) | N | N |
| Red-headed Vulture | *Sarcogyps calvus* | II | CR | 3500-15,000 | Decreasing | Y | 5.1.1. (ongoing) | N | N |
| Knob-billed Duck | *Sarkidiornis melanotos* | II | LC | 90,000-340,000 | Decreasing | Y | 5.1.1. (ongoing) | N | N |
| Yellow-fronted Canary | *Serinus mozambicus* | NC | LC | Unknown (common to locally abundant) | Decreasing | Y | | N | N |
| Crowned Eagle | *Stephanoaetus coronatus* | II | NT | 5000-50,000‡ | Decreasing | Y | 5.1.1. (ongoing) | N | N |
| Leopard Tortoise | *Stigmochelys pardalis* | II | LC | Unknown | Unknown | Y | 5.1.1. (ongoing) | N | N |
| Mourning Collared-dove | *Streptopelia decipiens* | NC | LC | Unknown (widespread and common) | Stable | N | | N | N |
| Red-eyed Dove | *Streptopelia semitorquata* | NC | LC | Unknown (but species is common) | Decreasing | N | | N | N |
| Laughing Dove | *Streptopelia senegalensis* | NC | LC | 2,400,000-8,200,000‡ | Stable | N | | N | N |
| Western Turtle-Dove | *Streptopelia turtur* | NC | VU | 19,300,000-71,400,00 | Decreasing | Y | 5.1.1. (ongoing) | N | N |
| Ural Owl | *Strix uralensis* | II | LC | 640,000-1,052,000‡ | Stable | N | | N | N |
| African Wood-owl | *Strix woodfordii* | II | LC | Unknown | Decreasing | N | | N | N |

| Common name | Scientific name | CITES Appendix | IUCN Red List Threat Category | Population size | Population trend | Intentional hunting or harvesting threat? (Yes/No) | Threat code(s) (timing) | Is trophy hunting a major contributor to elevated conservation concern (Yes/No) | Is trophy hunting likely or possibly causing localized declines (Likely/ Possibly /No) |
|---|---|---|---|---|---|---|---|---|---|
| Common Ostrich | *Struthio camelus* | I (populations of some African countries. | LC | Unknown | Decreasing | N | | N | N |
| Reeves's Pheast | *Syrmaticus reevesii* | II | VU | 3,500-15,000 | Decreasing | Y | 5.1.1. (ongoing) | N | N |
| Knysna Turaco | *Tauraco corythaix* | II | LC | Unknown (but common) | Decreasing | N | | N | N |
| Livingstone's Turaco | *Tauraco livingstonii* | II | LC | Unknown | Decreasing | N | | N | N |
| Yellow-billed Turaco | *Tauraco macrorhynchus* | II | LC | Unknown | Decreasing | N | | N | N |
| Green Turaco | *Tauraco persa* | II | LC | Unknown (but common) | Decreasing | N | | N | N |
| Purple-crested Turaco | *Tauraco porphyreolophus* | II | LC | Unknown | Decreasing | N | | N | N |
| White-lipped Peccary | *Tayassu pecari* | II | VU | Unknown | Decreasing | Y | 5.1.1. (ongoing) | N | N |
| Bateleur | *Terathopius ecaudatus* | II | EN | Unknown | Decreasing | Y | | N | N |
| Ornate Box Turtle | *Terrapene ornata* | II | NT | Unknown | Decreasing | Y | 5.4.1. (past, unlikely to return) | N | N |
| Gelada | *Theropithecus gelada* | II | LC | ~4300 | Decreasing | Y | 5.1.1. (past) | N | N |
| African Sacred Ibis | *Threskiornis aethiopicus* | NC | LC | 200,000-450,000 | Stable | Y | 5.1.1. (ongoing) | N | N |
| Lappet-faced Vulture | *Torgos tracheliotus* | II | EN | 9200 | Decreasing | Y | 5.1.1. (ongoing) | N | N |
| Sitatunga | *Tragelaphus spekii* | NC | LC | 170,000 | Decreasing | Y | 5.1.1. (ongoing) | N | N |
| African Green-Pigeon | *Treron calvus* | NC | LC | Unknown (but locally abundant) | Decreasing | N | | N | N |
| White-headed Vulture | *Trigonoceps occipitalis* | II | CR | 3,685† | Decreasing | Y | 5.1.1. (ongoing) | N | N |

| Common name | Scientific name | CITES Appendix | IUCN Red List Threat Category | Population size | Population trend | Intentional hunting or harvesting threat? (Yes/No) | Threat code(s) (timing) | Is trophy hunting a major contributor to elevated conservation concern (Yes/No) | Is trophy hunting likely or possibly causing localized declines (Likely/ Possibly /No) |
|---|---|---|---|---|---|---|---|---|---|
| Emmon's Black Bear | *Ursus americanus emmonsii* | II | NE | Unknown | Unknown | NE | | NE | NE |
| Himalayan Brown Bear | *Ursus arctos isabellinus* | I | EN | Unknown | Unknown | Y | 5.1.1. (ongoing) | N | N |
| Polar Bear | *Ursus maritimus* | II | VU | 26,000 | Unknown | Y | 5.1.1. (ongoing) | N | N |
| Asiatic Black Bear | *Ursus thibetanus* | I | VU | Unknown† | Decreasing | Y | 5.1.1. (ongoing) | N | N |
| White-throated Monitor | *Varanus albigularis* | II | LC | Unknown | Stable | N | | N | N |
| Savanna Monitor | *Varanus exanthematicus* | II | LC | Unknown† | Unknown | Y | 5.1.1. (ongoing) | N | N |
| Nile Monitor | *Varanus niloticus* | II | LC | Unknown (but naturally common) | Stable | Y | 5.1.1. (ongoing) | N | N |
| Common Water Monitor | *Varanus salvator* | II | LC | Unknown | Unknown | Y | 5.1.1. (ongoing) | N | N |
| Malabar Civet | *Viverra civettina* | III | CR | 249‡ | Decreasing | Y | 5.1.1. (ongoing) | N | N |
| Red Fox | *Vulpes vulpes* | NC (various subsp. in III) | LC | Unknown | Stable | Y | | N | N |
| Fennec Fox | *Vulpes zerda* | II | LC | Unknown (but common) | Stable | Y | 5.1.1. (ongoing) | N | N |

*Trophy hunting of wild populations does not take place formally and/or exports are exclusively, or close to exclusively, from extra-limital/non-wild populations.
†Regional estimates are in the Red List assessment.
‡Number of mature individuals.
§Based on CITES (2022) rather than the Red List.
¶Based on Sillero-Zubiri et al. (2004) as detailed in the Red List assessment for these species.
#Estimate for continental Africa rather than individual species.
**Species/subspecies that have not been formally evaluated on the Red List but have some information on threats in the associated assessment.

**Table S5. Impact of trophy hunting on CITES-listed species for which it is a localized threat (or was in the past) but does not contribute to the species being of elevated conservation concern. L=Likely, P=Possibly. Source: IUCN Red List (2026).**

| Species | Impact of trophy hunting | Source |
|---|---|---|
| Siberian Ibex [L]<br>*Capra sibirica* | If trophy hunting of this species is poorly regulated and managed it can have negative impacts, including on the age and sex composition of populations. These impacts are mitigated if only a minor proportion of males in a population are hunted. | Reading et al. (2020a) |
| Gobi argali [L]<br>*Ovis darwini (O.a. darwini*) | Impacts are not documented but trophy hunting is recognized as a minor local threat in Mongolia. | Reading et al. (2020b) |
| Leopard [L]<br>*Panthera pardus* | Intentional use of the species is a major threat, but the impact is not reported. Poorly managed trophy hunting can add pressure to local leopard populations and has been a key driver of population declines in parts of South Africa. | Stein et al. (2025) |
| Bongo [L]<br>*Tragelaphus eurycerus* | In a small portion of the species' range (Cameroon and Central African Republic) the species is threatened by demand for hunting trophies where this hunting is poorly regulated. Trophy hunting also has the potential to provide economic justification for the preservation of larger areas of Bongo habitat than national parks, especially in remote regions of Central Africa where possibilities for commercially successful tourism are very limited. | IUCN SSC Antelope Specialist Group (2017b) |
| Brown Bear [L]<br>*Ursus arctos* | Hunting rates may sometimes be unsustainable in the short term where populations exist in large, contiguous populations but cause population fluctuations rather than ongoing declines. | McLellan et al. (2017) |
| Lion [P]<br>*Panthera leo* | Intentional use of the species is a major threat, affecting a minority of the population with slow, significant declines. Trophy hunting is not a primary driver of this threat but is a potential threat where it is not well regulated. It may have, at times, contributed to population declines in Botswana, Namibia, Tanzania, Zimbabwe, Cameroon and Zambia. Trophy hunting can be an important tool for conserving lions if it is regulated and managed. | Nicholson et al. (2025) |
| Puma [P]<br>*Puma concolor* | The species is legally hunted in the US, which potentially comprises a minor, local threat. | Neilsen et al. (2016) |
| American Black Bear [P]<br>*Ursus americanus* | Trophy hunting of this species, although well controlled in North America, may contribute to population fluctuations. | Garshelis et al. (2016) |

**Table S6. CITES-listed species traded as hunting trophies (2000-2024; *n* = 340), estimated number of trophies, mean (± SD) number of trophies traded/year, WOEs traded in this period, and mean (± SD) WOEs traded in this period/year, ranked by number of trophies**. Synonyms on the IUCN Red List are presented in parentheses with scientific names. For some species trophy hunting does not take place formally and/or exports are exclusively, or close to exclusively, from extra-limital/non-wild populations, indicated by an asterix (*) on common names.

| Rank | Scientific name | Common name | Trophies | Mean + SD trophies/year | WOEs | Mean + SD WOEs/year |
|---|---|---|---|---|---|---|
| 1 | *Ursus americanus* | American Black Bear | 126974 | 5078.96 ± 1663.74 | 93280.25 | 3731.21 ± 1467.45 |
| 2 | *Crocodylus niloticus* | Nile Crocodile | 77802 | 3112.08 ± 4121.96 | 55205.25 | 2208.21 ± 2722.11 |
| 3 | *Loxodonta africana* | African Savanna Elephant | 52139 | 2085.58 ± 2172.33 | 17897.01 | 715.88 ± 567.46 |
| 4 | *Hippopotamus amphibius* | Common Hippopotamus | 41868 | 1674.72 ± 1396.94 | 13595.75 | 543.83 ± 283.41 |
| 5 | *Streptopelia turtur* | Western Turtle-Dove | 33020 | 1320.8 ± 2564.35 | 33020 | 1320.8 ± 2564.35 |
| 6 | *Equus zebra hartmannae* | Hartmann's Mountain Zebra | 31941 | 1277.64 ± 477.53 | 31129.25 | 1245.17 ± 473.01 |
| 7 | *Papio ursinus* | Chacma Baboon | 24940 | 997.6 ± 415.2 | 24545.25 | 981.81 ± 409.34 |
| 8 | *Ursus arctos* | Brown Bear | 19260 | 770.4 ± 857.63 | 15284 | 611.36 ± 289.49 |
| 9 | *Canis lupus* | Grey Wolf | 15536 | 621.44 ± 153.04 | 14657.25 | 586.29 ± 142.62 |
| 10 | *Panthera pardus* | Leopard | 15530 | 621.2 ± 231.17 | 15019.5 | 600.78 ± 213.77 |
| 11 | *Giraffa camelopardalis* | Giraffe | 12745 | 509.8 ± 1518.09 | 5562 | 222.48 ± 464.95 |
| 12 | *Caracal caracal* | Caracal | 11161 | 446.44 ± 159.59 | 11110 | 444.4 ± 159.53 |
| 13 | *Panthera leo* | Lion | 7777 | 311.08 ± 183.44 | 7095 | 283.8 ± 159.93 |
| 14 | *Kobus leche* | Lechwe | 7518 | 300.72 ± 150.8 | 7447 | 297.88 ± 148.35 |
| 15 | *Antilope cervicapra* | Blackbuck | 6973 | 278.92 ± 239.59 | 6972 | 278.88 ± 239.56 |
| 16 | *Meleagris ocellata* | Ocellated Turkey | 6342 | 253.68 ± 187.25 | 6201 | 248.04 ± 181.66 |
| 17 | *Puma concolor* | Puma | 5728 | 229.12 ± 86.95 | 5597 | 223.88 ± 82.3 |
| 18 | *Chlorocebus pygerythrus* | Vervet Monkey | 5695 | 227.8 ± 211.44 | 5681 | 227.24 ± 211.09 |
| 19 | *Ceratotherium simum simum* | Southern White Rhinoceros | 5369 | 214.76 ± 132.9 | 4561.25 | 182.45 ± 94.11 |
| 20 | *Salvator rufescens* | Red Tegu | 5000 | 200 ± 1000 | 5000 | 200 ± 1000 |
| 21 | *Lynx canadensis* | Canada Lynx | 4517 | 180.68 ± 144.72 | 3931.75 | 157.27 ± 114.34 |
| 22 | *Damaliscus pygargus pygargus* | Bontebok | 4155 | 166.2 ± 78.41 | 4147 | 165.88 ± 78.2 |
| 23 | *Civettictis civetta* | African Civet | 3436 | 137.44 ± 54.9 | 3426 | 137.04 ± 55.04 |
| 24 | *Ursus maritimus* | Polar Bear | 3183 | 127.32 ± 126.52 | 2314.5 | 92.58 ± 85.72 |

| Rank | Scientific name | Common name | Trophies | Mean + SD trophies/year | WOEs | Mean + SD WOEs/year |
|---|---|---|---|---|---|---|
| 25 | *Capra sibirica* | Siberian Ibex | 3171 | 126.84 ± 184.19 | 3171 | 126.84 ± 184.19 |
| 26 | *Ovis canadensis* | Bighorn Sheep | 2960 | 118.4 ± 41.71 | 2960 | 118.4 ± 41.71 |
| 27 | *Papio cynocephalus* | Yellow Baboon | 2744 | 109.76 ± 64.48 | 2728 | 109.12 ± 64.16 |
| 28 | *Chlorocebus aethiops* | Grivet Monkey | 2603 | 104.12 ± 131.56 | 2597 | 103.88 ± 131.34 |
| 29 | *Papio hamadryas* | Hamadryas Baboon | 2338 | 93.52 ± 130.83 | 2320.5 | 92.82 ± 129.68 |
| 30 | *Felis silvestris* | European Wildcat | 2251 | 90.04 ± 62.82 | 2241 | 89.64 ± 62.76 |
| 31 | *Damaliscus lunatus* | Topi | 2242 | 89.68 ± 123.17 | 2232 | 89.28 ± 122.48 |
| 32 | *Leptailurus serval* | Serval | 2158 | 86.32 ± 42.96 | 2084.75 | 83.39 ± 39.87 |
| 33 | *Acinonyx jubatus* | Cheetah | 2090 | 83.6 ± 41.55 | 1980 | 79.2 ± 34.44 |
| 34 | *Papio anubis* | Olive Baboon | 2078 | 83.12 ± 53.24 | 2071.5 | 82.86 ± 53.21 |
| 35 | *Philantomba monticola* | Blue Duiker | 2019 | 80.76 ± 70.01 | 1998 | 79.92 ± 68.84 |
| 36 | *Lynx rufus* | Bobcat | 1562 | 62.48 ± 83.53 | 1542 | 61.68 ± 83.76 |
| 37 | *Mellivora capensis* | Honey Badger | 1535 | 61.4 ± 36.35 | 1532 | 61.28 ± 36.24 |
| 38 | *Crax rubra* | Great Curassow | 1481 | 59.24 ± 53.71 | 1474 | 58.96 ± 53.63 |
| 39 | *Philantomba maxwellii* | Maxwell's Duiker | 1471 | 58.84 ± 76.42 | 1469 | 58.76 ± 76.46 |
| 40 | *Ovis ammon* | Wild Sheep | 1405 | 56.2 ± 33.64 | 1405 | 56.2 ± 33.64 |
| 41 | *Capra hircus aegagrus* | Wild Goat | 1192 | 47.68 ± 65.72 | 1191 | 47.64 ± 65.71 |
| 42 | *Arctocephalus pusillus* | Afro-Australian Fur Seal | 1043 | 41.72 ± 199.67 | 1038 | 41.52 ± 199.71 |
| 43 | *Alligator mississippiensis* | American Alligator | 972 | 38.88 ± 113.44 | 878 | 35.12 ± 110.49 |
| 44 | *Cerberus rynchops* | South Asian Bockadam | 936 | 37.44 ± 187.2 | 0 | 0 ± 0 |
| 45 | *Anas crecca* | Eurasian Teal | 854 | 34.16 ± 71.97 | 854 | 34.16 ± 71.97 |
| 46 | *Lontra canadensis* | North American River Otter | 848 | 33.92 ± 106.5 | 780 | 31.2 ± 106.41 |
| 47 | *Ovis darwini (Ovis ammon darwini)* | Gobi Argali | 822 | 32.88 ± 21.17 | 822 | 32.88 ± 21.17 |
| 48 | *Penelope purpurascens* | Crested Guan | 820 | 32.8 ± 30.52 | 819 | 32.76 ± 30.49 |
| 49 | *Ammotragus lervia* | Aoudad | 799 | 31.96 ± 19.19 | 799 | 31.96 ± 19.19 |
| 50 | *Ovis polii* | Marco Polo Argali | 661 | 26.44 ± 49.22 | 661 | 26.44 ± 49.22 |
| 51 | *Anas acuta* | Northern Pintail | 646 | 25.84 ± 56.55 | 646 | 25.84 ± 56.55 |
| 52 | *Anas clypeata (Spatula clypeata)* | Northern Shoveler | 632 | 25.28 ± 56.99 | 632 | 25.28 ± 56.99 |
| 53 | *Anas penelope* | Eurasian Wigeon | 630 | 25.2 ± 59.59 | 630 | 25.2 ± 59.59 |
| 54 | *Proteles cristata* | Aardwolf | 595 | 23.8 ± 17.79 | 594 | 23.76 ± 17.74 |

| Rank | Scientific name | Common name | Trophies | Mean + SD trophies/year | WOEs | Mean + SD WOEs/year |
|---|---|---|---|---|---|---|
| 55 | *Tragelaphus eurycerus* | Bongo | 555 | 22.2 ± 32.64 | 555 | 22.2 ± 32.64 |
| 56 | *Nasua narica* | White-nosed Coati | 531 | 21.24 ± 21.25 | 531 | 21.24 ± 21.25 |
| 57 | *Capra caucasica* | Western Tur | 512 | 20.48 ± 36.93 | 512 | 20.48 ± 36.93 |
| 58 | *Felis lybica* | Afro-Asiatic Wildcat | 493 | 19.72 ± 25.3 | 492 | 19.68 ± 25.27 |
| 59 | *Alopochen aegyptiaca* | Egyptian Goose | 473 | 18.92 ± 33.05 | 449 | 17.96 ± 31.68 |
| 60 | *Puma concolor couguar* | Florida Puma | 401 | 16.04 ± 52.1 | 329 | 13.16 ± 39.79 |
| 61 | *Python brongersmai* | Brongersma's Short-tailed Python | 400 | 16 ± 80 | 400 | 16 ± 80 |
| 62 | *Odobenus rosmarus* | Walrus | 392 | 15.68 ± 20.31 | 132 | 5.28 ± 4.77 |
| 63 | *Ovis aries* | Urial | 372 | 14.88 ± 12.86 | 372 | 14.88 ± 12.86 |
| 64 | *Equus zebra* | Mountain Zebra | 358 | 14.32 ± 70.98 | 354 | 14.16 ± 70.18 |
| 65 | *Cephalophus silvicultor* | Yellow-backed Duiker | 338 | 13.52 ± 13.18 | 336 | 13.44 ± 12.96 |
| 66 | *Stigmochelys pardalis* | Leopard Tortoise | 314 | 12.56 ± 59.9 | 14 | 0.56 ± 1.42 |
| 67 | *Lynx lynx* | Eurasian lynx | 277 | 11.08 ± 8.77 | 276 | 11.04 ± 8.75 |
| 68 | *Cephalophus dorsalis* | Bay Duiker | 269 | 10.76 ± 8.29 | 269 | 10.76 ± 8.29 |
| 69 | *Tragelaphus spekii* | Sitatunga | 269 | 10.76 ± 18.9 | 269 | 10.76 ± 18.9 |
| 70 | *Plectropterus gambensis* | Spur-winged Goose | 254 | 10.16 ± 16.68 | 230 | 9.2 ± 15.54 |
| 71 | *Colobus guereza* | Guereza | 229 | 9.16 ± 6.61 | 229 | 9.16 ± 6.61 |
| 72 | *Anas querquedula (Spatula querquedula)* | Garganey | 219 | 8.76 ± 19.89 | 219 | 8.76 ± 19.89 |
| 73 | *Canis aureus* | Common Jackal | 213 | 8.52 ± 7.14 | 213 | 8.52 ± 7.14 |
| 74 | *Elaphe radiata (Coelognathus radiatus)* | Copper-head Trinket Snake | 206 | 8.24 ± 41.2 | 206 | 8.24 ± 41.2 |
| 75 | *Capra falconeri* | Markhor | 192 | 7.68 ± 4.26 | 192 | 7.68 ± 4.26 |
| 76 | *Damaliscus pygargus* | Blesbok | 192 | 7.68 ± 14.47 | 192 | 7.68 ± 14.47 |
| 77 | *Ortalis vetula* | Plain Chachalaca | 185 | 7.4 ± 9.62 | 185 | 7.4 ± 9.62 |
| 78 | *Ceratotherium simum* | White Rhinoceros | 182 | 7.28 ± 8.12 | 151.5 | 6.06 ± 5.8 |
| 79 | *Oryx dammah* | Scimitar-horned Oryx | 175 | 7 ± 8.67 | 174 | 6.96 ± 8.51 |
| 80 | *Dendrocygna autumnalis* | Black-bellied Whistling-duck | 152 | 6.08 ± 17.93 | 152 | 6.08 ± 17.93 |
| 81 | *Pecari tajacu* | Collared Pecary | 150 | 6 ± 12.1 | 150 | 6 ± 12.1 |
| 82 | *Dendrocygna bicolor* | Fulvous Whistling-duck | 137 | 5.48 ± 16.76 | 137 | 5.48 ± 16.76 |
| 83 | *Python sebae* | Central African Rock Python | 128 | 5.12 ± 5.16 | 128 | 5.12 ± 5.16 |
| 84 | *Pseudois nayaur* | Blue Sheep | 115 | 4.6 ± 7.3 | 115 | 4.6 ± 7.3 |

| Rank | Scientific name | Common name | Trophies | Mean + SD trophies/year | WOEs | Mean + SD WOEs/year |
|---|---|---|---|---|---|---|
| 85 | *Ovis bochariensis (Ovis v. bochariensis)* | Bukhara Urial | 107 | 4.28 ± 7.71 | 107 | 4.28 ± 7.71 |
| 86 | *Bison bison athabascae* | Wood Bison | 101 | 4.04 ± 4.93 | 94 | 3.76 ± 4.71 |
| 87 | *Dendrocygna viduata* | White-faced Whistling duck | 100 | 4 ± 7.58 | 100 | 4 ± 7.58 |
| 88 | *Malayopython reticulatus* | Reticulated Python | 100 | 4 ± 20 | 100 | 4 ± 20 |
| 89 | *Equus zebra zebra* | Cape Mountain Zebra | 92 | 3.68 ± 7.87 | 89 | 3.56 ± 7.86 |
| 90 | *Dasyprocta punctata* | Central American Agouti | 89 | 3.56 ± 5.43 | 89 | 3.56 ± 5.43 |
| 91 | *Diceros bicornis* | Black Rhinoceros | 88 | 3.52 ± 3.39 | 88 | 3.52 ± 3.39 |
| 92 | *Ovis cycloceros (O. v. cycloceros)* | Afghan Urial | 86 | 3.44 ± 12.25 | 86 | 3.44 ± 12.25 |
| 93 | *Pan troglodytes* | Chimpanzee | 82 | 3.28 ± 15.79 | 0 | 0 ± 0 |
| 94 | *Capra falconeri heptneri* | Markhor | 79 | 3.16 ± 5.52 | 79 | 3.16 ± 5.52 |
| 95 | *Axis porcinus* | Hog Deer | 78 | 3.12 ± 5.89 | 78 | 3.12 ± 5.89 |
| 96 | *Sarkidiornis melanotos* | Knob-billed Duck | 70 | 2.8 ± 3.23 | 70 | 2.8 ± 3.23 |
| 97 | *Columba guinea* | Speckled Pigeon | 69 | 2.76 ± 8.66 | 57 | 2.28 ± 8.4 |
| 98 | *Caiman crocodilus fuscus* | Brown Spectacled Caiman | 66 | 2.64 ± 13.2 | 0 | 0 ± 0 |
| 99 | *Capra hircus* | Goat | 64 | 2.56 ± 11.59 | 64 | 2.56 ± 11.59 |
| 100 | *Ovis cycloceros cycloceros (O.v. cycloceros)* | Afghan Urial (subsp. cycloceros) | 63 | 2.52 ± 7.65 | 63 | 2.52 ± 7.65 |
| 101 | *Gazella dorcas* | Dorcas Gazelle | 58 | 2.32 ± 6.23 | 58 | 2.32 ± 6.23 |
| 102 | *Ovis cycloceros arkal (O.v. arkal)* | Afghan Urial (subsp. arkal) | 57 | 2.28 ± 5.08 | 57 | 2.28 ± 5.08 |
| 103 | *Streptopelia semitorquata* | Red-eyed Dove | 57 | 2.28 ± 8.47 | 57 | 2.28 ± 8.47 |
| 104 | *Buteo jamaicensis* | Red-tailed Hawk | 52 | 2.08 ± 9.99 | 0 | 0 ± 0 |
| 105 | *Ovis punjabiensis (Ovis v. punjabensis)* | Punjab Urial | 51 | 2.04 ± 4.92 | 51 | 2.04 ± 4.92 |
| 106 | *Streptopelia senegalensis* | Laughing Dove | 51 | 2.04 ± 6.89 | 51 | 2.04 ± 6.89 |
| 107 | *Theropithecus gelada* | Gelada | 51 | 2.04 ± 1.7 | 51 | 2.04 ± 1.7 |
| 108 | *Erythrocebus patas* | Patas Monkey | 50 | 2 ± 2.35 | 50 | 2 ± 2.35 |
| 109 | *Budorcas taxicolor* | Takin | 45 | 1.8 ± 4.55 | 44 | 1.76 ± 4.54 |
| 110 | *Anas capensis* | Cape Teal | 41 | 1.64 ± 4.63 | 23 | 0.92 ± 2.89 |
| 111 | *Bubalus arnee* | Wild Water Buffalo | 35 | 1.4 ± 5.67 | 9 | 0.36 ± 1.25 |
| 112 | *Loxodonta cyclotis* | African Forest Elephant | 34 | 1.36 ± 3.53 | 23 | 0.92 ± 2.4 |
| 113 | *Tayassu pecari* | White-lipped Peccary | 34 | 1.36 ± 2.63 | 34 | 1.36 ± 2.63 |
| 114 | *Hyaena hyaena* | Striped Hyena | 33 | 1.32 ± 2.63 | 33 | 1.32 ± 2.63 |

| Rank | Scientific name | Common name | Trophies | Mean + SD trophies/year | WOEs | Mean + SD WOEs/year |
|---|---|---|---|---|---|---|
| 115 | *Hystrix cristata* | Crested Porcupine | 33 | 1.32 ± 1.89 | 33 | 1.32 ± 1.89 |
| 116 | *Canis lupus monstrabilis* | Texas Grey Wolf | 29 | 1.16 ± 3.17 | 29 | 1.16 ± 3.17 |
| 117 | *Antigone canadensis (Grus canadensis)* | Sandhill Crane | 26 | 1.04 ± 1.4 | 26 | 1.04 ± 1.4 |
| 118 | *Cercopithecus mitis* | Blue Monkey | 25 | 1 ± 1.08 | 25 | 1 ± 1.08 |
| 119 | *Crocodylus porosus* | Saltwater Crocodile | 24 | 0.96 ± 1.31 | 24 | 0.96 ± 1.31 |
| 120 | *Anas nesiotis* | Campbell Teal | 20 | 0.8 ± 4 | 20 | 0.8 ± 4 |
| 121 | *Cephalophus zebra* | Zebra Duiker | 20 | 0.8 ± 2.81 | 20 | 0.8 ± 2.81 |
| 122 | *Cuniculus paca* | Paca | 20 | 0.8 ± 1.29 | 20 | 0.8 ± 1.29 |
| 123 | *Varanus niloticus* | Nile Monitor | 20 | 0.8 ± 1.29 | 20 | 0.8 ± 1.29 |
| 124 | *Ovis karelini (O.a. karelini)* | Tianshan Argali | 19 | 0.76 ± 1.74 | 19 | 0.76 ± 1.74 |
| 125 | *Columba livia* | Rock Dove | 18 | 0.72 ± 2.46 | 18 | 0.72 ± 2.46 |
| 126 | *Myiopsitta monachus* | Monk Parakeet | 17 | 0.68 ± 3.2 | 17 | 0.68 ± 3.2 |
| 127 | *Necrosyrtes monachus* | Hooded Vulture | 17 | 0.68 ± 1.77 | 17 | 0.68 ± 1.77 |
| 128 | *Papio papio* | Guinea Baboon | 17 | 0.68 ± 1.89 | 17 | 0.68 ± 1.89 |
| 129 | *Varanus albigularis* | White-throated Monitor | 17 | 0.68 ± 1.38 | 17 | 0.68 ± 1.38 |
| 130 | *Mazama temama* | Central American Red Brocket | 16 | 0.64 ± 3 | 16 | 0.64 ± 3 |
| 131 | *Otolemur crassicaudatus* | Thick-tailed Greater Galago | 16 | 0.64 ± 1.32 | 16 | 0.64 ± 1.32 |
| 132 | *Ovis severtzovi (O.a. severtzovi)* | Severtzov's Argali | 15 | 0.6 ± 1 | 15 | 0.6 ± 1 |
| 133 | *Ursus thibetanus* | Asiatic Black Bear | 15 | 0.6 ± 1.68 | 15 | 0.6 ± 1.68 |
| 134 | *Moschus moschiferus* | Siberian Musk Deer | 13 | 0.52 ± 1 | 13 | 0.52 ± 1 |
| 135 | *Neotis denhami* | Denham's Bustard | 13 | 0.52 ± 1.12 | 13 | 0.52 ± 1.12 |
| 136 | *Lycalopex griseus* | Chilla | 12 | 0.48 ± 1.08 | 12 | 0.48 ± 1.08 |
| 137 | *Aonyx capensis* | African Clawless Otter | 10 | 0.4 ± 0.71 | 10 | 0.4 ± 0.71 |
| 138 | *Cervus elaphus* | Red Deer | 10 | 0.4 ± 1.41 | 10 | 0.4 ± 1.41 |
| 139 | *Ardeotis arabs* | Arabian Bustard | 9 | 0.36 ± 1.8 | 9 | 0.36 ± 1.8 |
| 140 | *Cercopithecus nictitans* | Putty-nosed Monkey | 9 | 0.36 ± 0.99 | 9 | 0.36 ± 0.99 |
| 141 | *Haliaeetus vocifer* | African Fish-eagle | 9 | 0.36 ± 1.41 | 9 | 0.36 ± 1.41 |
| 142 | *Oryx leucoryx* | Arabian Oryx | 8 | 0.32 ± 0.69 | 8 | 0.32 ± 0.69 |
| 143 | *Pavo cristatus* | Indian Peafowl | 8 | 0.32 ± 0.75 | 8 | 0.32 ± 0.75 |
| 144 | *Streptopelia decipiens* | Mourning Collared-dove | 8 | 0.32 ± 1.41 | 8 | 0.32 ± 1.41 |

| Rank | Scientific name | Common name | Trophies | Mean + SD trophies/year | WOEs | Mean + SD WOEs/year |
|---|---|---|---|---|---|---|
| 145 | *Tauraco livingstonii* | Livingstone's Turaco | 8 | 0.32 ± 1.6 | 8 | 0.32 ± 1.6 |
| 146 | *Cephalophus ogilbyi* | Ogilby's Duiker | 7 | 0.28 ± 0.79 | 7 | 0.28 ± 0.79 |
| 147 | *Mazama temama cerasina* | Guatemalan Red Brocket | 7 | 0.28 ± 1.21 | 7 | 0.28 ± 1.21 |
| 148 | *Ovis gmelini* | Mouflon | 7 | 0.28 ± 0.61 | 7 | 0.28 ± 0.61 |
| 149 | *Ovis vignei* | Urial | 7 | 0.28 ± 0.98 | 7 | 0.28 ± 0.98 |
| 150 | *Sagittarius serpentarius* | Secretarybird | 7 | 0.28 ± 0.61 | 7 | 0.28 ± 0.61 |
| 151 | *Tauraco macrorhynchus* | Yellow-billed Turaco | 7 | 0.28 ± 1.21 | 7 | 0.28 ± 1.21 |
| 152 | *Aegypius monachus* | Cinereous Vulture | 6 | 0.24 ± 0.83 | 6 | 0.24 ± 0.83 |
| 153 | *Ardeotis kori* | Kori Bustard | 6 | 0.24 ± 0.52 | 6 | 0.24 ± 0.52 |
| 154 | *Athene cunicularia* | Burrowing Owl | 6 | 0.24 ± 1.2 | 6 | 0.24 ± 1.2 |
| 155 | *Bison bison* | American Bison | 6 | 0.24 ± 1.2 | 5 | 0.2 ± 1 |
| 156 | *Cairina moschata* | Muscovy Duck | 6 | 0.24 ± 0.66 | 0 | 0 ± 0 |
| 157 | *Circaetus fasciolatus* | Southern Banded Snake-eagle | 6 | 0.24 ± 1.2 | 6 | 0.24 ± 1.2 |
| 158 | *Galago moholi* | Southern Lesser Galago | 6 | 0.24 ± 0.83 | 6 | 0.24 ± 0.83 |
| 159 | *Galago senegalensis* | Northern Lesser Galago | 6 | 0.24 ± 0.6 | 6 | 0.24 ± 0.6 |
| 160 | *Gyps africanus* | White-backed Vulture | 6 | 0.24 ± 0.72 | 6 | 0.24 ± 0.72 |
| 161 | *Hippotragus equinus* | Roan Antelope | 6 | 0.24 ± 0.83 | 6 | 0.24 ± 0.83 |
| 162 | *Hyaena brunnea* | Brown Hyaena | 6 | 0.24 ± 0.88 | 6 | 0.24 ± 0.88 |
| 163 | *Hyemoschus aquaticus* | Water Chevrotain | 6 | 0.24 ± 1.2 | 6 | 0.24 ± 1.2 |
| 164 | *Strix woodfordii* | African Wood-owl | 6 | 0.24 ± 1.2 | 6 | 0.24 ± 1.2 |
| 165 | *Trigonoceps occipitalis* | White-headed Vulture | 6 | 0.24 ± 0.88 | 6 | 0.24 ± 0.88 |
| 166 | *Varanus exanthematicus* | Savanna Monitor | 6 | 0.24 ± 0.6 | 6 | 0.24 ± 0.6 |
| 167 | *Capra falconeri megaceros* | Straight-horned Markhor | 5 | 0.2 ± 0.65 | 5 | 0.2 ± 0.65 |
| 168 | *Cercopithecus cephus* | Moustached Monkey | 5 | 0.2 ± 0.82 | 5 | 0.2 ± 0.82 |
| 169 | *Chlorocebus tantalus* | Tantalus Monkey | 5 | 0.2 ± 1 | 5 | 0.2 ± 1 |
| 170 | *Glaucidium capense* | African Barred Owlet | 5 | 0.2 ± 1 | 5 | 0.2 ± 1 |
| 171 | *Gyps rueppellii* | Rüppell's Vulture | 5 | 0.2 ± 0.65 | 5 | 0.2 ± 0.65 |
| 172 | *Lycalopex gymnocercus* | Pampas Fox | 5 | 0.2 ± 0.65 | 5 | 0.2 ± 0.65 |
| 173 | *Oena capensis* | Namaqua Dove | 5 | 0.2 ± 0.82 | 5 | 0.2 ± 0.82 |
| 174 | *Poicephalus cryptoxanthus* | Brown-headed Parrot | 5 | 0.2 ± 1 | 5 | 0.2 ± 1 |

| Rank | Scientific name | Common name | Trophies | Mean + SD trophies/year | WOEs | Mean + SD WOEs/year |
|---|---|---|---|---|---|---|
| 175 | *Tauraco porphyreolophus* | Purple-crested Turaco | 5 | 0.2 ± 0.58 | 5 | 0.2 ± 0.58 |
| 176 | *Accipiter gentilis* | Northern Goshawk | 4 | 0.16 ± 0.8 | 4 | 0.16 ± 0.8 |
| 177 | *Addax nasomaculatus* | Addax | 4 | 0.16 ± 0.47 | 4 | 0.16 ± 0.47 |
| 178 | *Buteo buteo* | Eurasian Buzzard | 4 | 0.16 ± 0.62 | 4 | 0.16 ± 0.62 |
| 179 | *Circaetus pectoralis* | Black-chested Snake-eagle | 4 | 0.16 ± 0.47 | 3 | 0.12 ± 0.44 |
| 180 | *Eulampis jugularis* | Purple-throated Carib | 4 | 0.16 ± 0.8 | 4 | 0.16 ± 0.8 |
| 181 | *Falco naumanni* | Lesser Kestrel | 4 | 0.16 ± 0.62 | 4 | 0.16 ± 0.62 |
| 182 | *Ithaginis cruentus* | Blood Pheasant | 4 | 0.16 ± 0.8 | 4 | 0.16 ± 0.8 |
| 183 | *Kaupifalco monogrammicus* | Lizard Buzzard | 4 | 0.16 ± 0.62 | 4 | 0.16 ± 0.62 |
| 184 | *Lophocebus albigena* | Grey-cheeked Mangabey | 4 | 0.16 ± 0.62 | 4 | 0.16 ± 0.62 |
| 185 | *Lycalopex culpaeus* | Culpeo | 4 | 0.16 ± 0.8 | 4 | 0.16 ± 0.8 |
| 186 | *Polemaetus bellicosus* | Martial Eagle | 4 | 0.16 ± 0.47 | 4 | 0.16 ± 0.47 |
| 187 | *Aquila chrysaetos* | Golden Eagle | 3 | 0.12 ± 0.6 | 3 | 0.12 ± 0.6 |
| 188 | *Caracal aurata* | African Golden Cat | 3 | 0.12 ± 0.44 | 3 | 0.12 ± 0.44 |
| 189 | *Elanus caeruleus* | Black-winged Kite | 3 | 0.12 ± 0.6 | 3 | 0.12 ± 0.6 |
| 190 | *Eupodotis senegalensis* | White-bellied Bustard | 3 | 0.12 ± 0.44 | 3 | 0.12 ± 0.44 |
| 191 | *Falco tinnunculus* | Common Kestel | 3 | 0.12 ± 0.6 | 3 | 0.12 ± 0.6 |
| 192 | *Gallus sonneratii* | Grey Jungefowl | 3 | 0.12 ± 0.6 | 3 | 0.12 ± 0.6 |
| 193 | *Glaucidium perlatum* | Pearl-spotted Owlet | 3 | 0.12 ± 0.6 | 3 | 0.12 ± 0.6 |
| 194 | *Haliaeetus leucocephalus* | Bald Eagle | 3 | 0.12 ± 0.44 | 0 | 0 ± 0 |
| 195 | *Haliaeetus leucoryphus* | Pallas's Fish-eagle | 3 | 0.12 ± 0.44 | 2 | 0.08 ± 0.4 |
| 196 | *Hydrictis maculicollis* | Spotted-necked Otter | 3 | 0.12 ± 0.44 | 3 | 0.12 ± 0.44 |
| 197 | *Lissotis melanogaster* | Black-bellied Bustard | 3 | 0.12 ± 0.44 | 3 | 0.12 ± 0.44 |
| 198 | *Macrochelys temminckii* | Alligator Snapping Turtle | 3 | 0.12 ± 0.6 | 3 | 0.12 ± 0.6 |
| 199 | *Manis tricuspis* | White-bellied Pangolin | 3 | 0.12 ± 0.33 | 3 | 0.12 ± 0.33 |
| 200 | *Odocoileus virginianus mayensis* | Guatemalan White-tailed Deer | 3 | 0.12 ± 0.44 | 3 | 0.12 ± 0.44 |
| 201 | *Ovis nigrimontana (O.a. nigrimontana)* | Karatau Argali | 3 | 0.12 ± 0.6 | 3 | 0.12 ± 0.6 |
| 202 | *Oxyura jamaicensis* | Ruddy Duck | 3 | 0.12 ± 0.6 | 3 | 0.12 ± 0.6 |
| 203 | *Phoeniconaias minor* | Lesser Flamingo | 3 | 0.12 ± 0.6 | 3 | 0.12 ± 0.6 |
| 204 | *Poicephalus fuscicollis* | Brown-necked Parrot | 3 | 0.12 ± 0.6 | 3 | 0.12 ± 0.6 |

| Rank | Scientific name | Common name | Trophies | Mean + SD trophies/year | WOEs | Mean + SD WOEs/year |
|---|---|---|---|---|---|---|
| 205 | *Sarcogyps calvus* | Red-headed Vulture | 3 | 0.12 ± 0.6 | 3 | 0.12 ± 0.6 |
| 206 | *Terathopius ecaudatus* | Bateleur | 3 | 0.12 ± 0.44 | 3 | 0.12 ± 0.44 |
| 207 | *Treron calvus* | African Green-Pigeon | 3 | 0.12 ± 0.6 | 3 | 0.12 ± 0.6 |
| 208 | *Ursus arctos isabellinus* | Himalayan Brown Bear | 3 | 0.12 ± 0.44 | 3 | 0.12 ± 0.44 |
| 209 | *Afrotis afra* | Southern Black Bustard | 2 | 0.08 ± 0.4 | 2 | 0.08 ± 0.4 |
| 210 | *Afrotis afraoides* | Northern Black Bustard | 2 | 0.08 ± 0.4 | 2 | 0.08 ± 0.4 |
| 211 | *Antilocapra americana* | Pronghorn | 2 | 0.08 ± 0.28 | 2 | 0.08 ± 0.28 |
| 212 | *Apalone mutica* | Smooth Softshell Turtle | 2 | 0.08 ± 0.4 | 2 | 0.08 ± 0.4 |
| 213 | *Aquila nipalensis* | Steppe Eagle | 2 | 0.08 ± 0.28 | 2 | 0.08 ± 0.28 |
| 214 | *Aquila rapax* | Tawny Eagle | 2 | 0.08 ± 0.28 | 2 | 0.08 ± 0.28 |
| 215 | *Athene noctua* | Little Owl | 2 | 0.08 ± 0.4 | 2 | 0.08 ± 0.4 |
| 216 | *Boselaphus tragocamelus* | Nilgai | 2 | 0.08 ± 0.28 | 2 | 0.08 ± 0.28 |
| 217 | *Bubo virginianus* | Great Horned Owl | 2 | 0.08 ± 0.28 | 2 | 0.08 ± 0.28 |
| 218 | *Buteo augur* | Augur Buzzard | 2 | 0.08 ± 0.4 | 2 | 0.08 ± 0.4 |
| 219 | *Buteo rufinus* | Long-legged Buzzard | 2 | 0.08 ± 0.4 | 2 | 0.08 ± 0.4 |
| 220 | *Caiman crocodilus* | Spectacled Caiman | 2 | 0.08 ± 0.4 | 2 | 0.08 ± 0.4 |
| 221 | *Caiman crocodilus crocodilus* | South American Spectacled Caiman | 2 | 0.08 ± 0.28 | 2 | 0.08 ± 0.28 |
| 222 | *Cercopithecus mona* | Mona Monkey | 2 | 0.08 ± 0.4 | 2 | 0.08 ± 0.4 |
| 223 | *Cercopithecus petaurista* | Spot-nosed Monkey | 2 | 0.08 ± 0.4 | 2 | 0.08 ± 0.4 |
| 224 | *Cercopithecus pogonias* | Crowned Monkey | 2 | 0.08 ± 0.4 | 0 | 0 ± 0 |
| 225 | *Chlorocebus sabaeus* | Green Monkey | 2 | 0.08 ± 0.4 | 2 | 0.08 ± 0.4 |
| 226 | *Ciconia nigra* | Black Stork | 2 | 0.08 ± 0.4 | 2 | 0.08 ± 0.4 |
| 227 | *Circaetus cinereus* | Brown Snake-eagle | 2 | 0.08 ± 0.28 | 2 | 0.08 ± 0.28 |
| 228 | *Colibri delphinae* | Brown Violetear | 2 | 0.08 ± 0.4 | 2 | 0.08 ± 0.4 |
| 229 | *Colobus polykomos* | King Colobus | 2 | 0.08 ± 0.4 | 2 | 0.08 ± 0.4 |
| 230 | *Crotalus durissus* | Cascabel Rattlesnake | 2 | 0.08 ± 0.28 | 2 | 0.08 ± 0.28 |
| 231 | *Egretta garzetta* | Little Egret | 2 | 0.08 ± 0.28 | 2 | 0.08 ± 0.28 |
| 232 | *Equus grevyi* | Grevy's Zebra | 2 | 0.08 ± 0.4 | 2 | 0.08 ± 0.4 |
| 233 | *Falco cherrug* | Saker Falcon | 2 | 0.08 ± 0.28 | 2 | 0.08 ± 0.28 |
| 234 | *Gypaetus barbatus* | Bearded Vulture | 2 | 0.08 ± 0.28 | 2 | 0.08 ± 0.28 |

| Rank | Scientific name | Common name | Trophies | Mean + SD trophies/year | WOEs | Mean + SD WOEs/year |
|---|---|---|---|---|---|---|
| 235 | *Gypohierax angolensis* | Palm-nut Vulture | 2 | 0.08 ± 0.4 | 2 | 0.08 ± 0.4 |
| 236 | *Hippotragus niger* | Sable Antelope | 2 | 0.08 ± 0.4 | 1 | 0.04 ± 0.2 |
| 237 | *Hippotragus niger variani* | Giant Sable | 2 | 0.08 ± 0.4 | 2 | 0.08 ± 0.4 |
| 238 | *Lophaetus occipitalis* | Long-crested Eagle | 2 | 0.08 ± 0.28 | 2 | 0.08 ± 0.28 |
| 239 | *Lutra lutra* | Eurasian Otter | 2 | 0.08 ± 0.4 | 2 | 0.08 ± 0.4 |
| 240 | *Macaca fascicularis* | Long-tailed Macaque | 2 | 0.08 ± 0.28 | 2 | 0.08 ± 0.28 |
| 241 | *Miopithecus talapoin* | Southern Talapoin Monkey | 2 | 0.08 ± 0.4 | 2 | 0.08 ± 0.4 |
| 242 | *Nanger dama* | Dama Gazelle | 2 | 0.08 ± 0.4 | 2 | 0.08 ± 0.4 |
| 243 | *Orycteropus afer* | Aardvark | 2 | 0.08 ± 0.4 | 2 | 0.08 ± 0.4 |
| 244 | *Oxyura leucocephala* | White-headed Duck | 2 | 0.08 ± 0.4 | 2 | 0.08 ± 0.4 |
| 245 | *Perodicticus potto* | West African Potto | 2 | 0.08 ± 0.4 | 2 | 0.08 ± 0.4 |
| 246 | *Platycercus eximius* | Eastern Rosella | 2 | 0.08 ± 0.4 | 0 | 0 ± 0 |
| 247 | *Poicephalus gulielmi* | Red-fronted Parrot | 2 | 0.08 ± 0.28 | 2 | 0.08 ± 0.28 |
| 248 | *Polyboroides typus* | African Harrier-hawk | 2 | 0.08 ± 0.28 | 2 | 0.08 ± 0.28 |
| 249 | *Python regius* | Ball Python | 2 | 0.08 ± 0.4 | 2 | 0.08 ± 0.4 |
| 250 | *Struthio camelus* | Common Ostrich | 2 | 0.08 ± 0.4 | 2 | 0.08 ± 0.4 |
| 251 | *Syrmaticus reevesii* | Reeves's Pheasant | 2 | 0.08 ± 0.4 | 2 | 0.08 ± 0.4 |
| 252 | *Tauraco persa* | Green Turaco | 2 | 0.08 ± 0.4 | 2 | 0.08 ± 0.4 |
| 253 | *Torgos tracheliotus* | Lappet-faced Vulture | 2 | 0.08 ± 0.28 | 2 | 0.08 ± 0.28 |
| 254 | *Vulpes vulpes* | Red Fox | 2 | 0.08 ± 0.28 | 2 | 0.08 ± 0.28 |
| 255 | *Vulpes zerda* | Fennec Fox | 2 | 0.08 ± 0.4 | 2 | 0.08 ± 0.4 |
| 256 | *Accipiter melanochlamys* | Black-mantled Goshawk | 1 | 0.04 ± 0.2 | 1 | 0.04 ± 0.2 |
| 257 | *Accipiter melanoleucus* | Black Sparrowhawk | 1 | 0.04 ± 0.2 | 1 | 0.04 ± 0.2 |
| 258 | *Accipiter striatus* | Sharp-shinned Hawk | 1 | 0.04 ± 0.2 | 1 | 0.04 ± 0.2 |
| 259 | *Anas formosa (Sibirionetta formosa)* | Baikal Teal | 1 | 0.04 ± 0.2 | 1 | 0.04 ± 0.2 |
| 260 | *Anthropoides virgo* | Demoiselle Crane | 1 | 0.04 ± 0.2 | 1 | 0.04 ± 0.2 |
| 261 | *Aquila verreauxii* | Verreaux's Eagle | 1 | 0.04 ± 0.2 | 0 | 0 ± 0 |
| 262 | *Ara chloropterus* | Red-and-green Macaw | 1 | 0.04 ± 0.2 | 1 | 0.04 ± 0.2 |
| 263 | *Asio capensis* | Marsh Owl | 1 | 0.04 ± 0.2 | 1 | 0.04 ± 0.2 |
| 264 | *Asio otus* | Northern Long-eared Owl | 1 | 0.04 ± 0.2 | 1 | 0.04 ± 0.2 |

| Rank | Scientific name | Common name | Trophies | Mean + SD trophies/year | WOEs | Mean + SD WOEs/year |
|---|---|---|---|---|---|---|
| 265 | *Bos mutus* | Wild Yak | 1 | 0.04 ± 0.2 | 1 | 0.04 ± 0.2 |
| 266 | *Bostrychia hagedash* | Hadada Ibis | 1 | 0.04 ± 0.2 | 1 | 0.04 ± 0.2 |
| 267 | *Branta canadensis leucopareia* | Aleutian Canada Goose | 1 | 0.04 ± 0.2 | 1 | 0.04 ± 0.2 |
| 268 | *Bubo bubo* | Eurasian Eagle-owl | 1 | 0.04 ± 0.2 | 1 | 0.04 ± 0.2 |
| 269 | *Bubo capensis* | Cape Eagle-owl | 1 | 0.04 ± 0.2 | 1 | 0.04 ± 0.2 |
| 270 | *Butastur rufipennis* | Grasshopper Buzzard | 1 | 0.04 ± 0.2 | 1 | 0.04 ± 0.2 |
| 271 | *Buteo rufofuscus* | Jackal Buzzard | 1 | 0.04 ± 0.2 | 1 | 0.04 ± 0.2 |
| 272 | *Cacatua galerita* | Sulphur-crested Cockatoo | 1 | 0.04 ± 0.2 | 1 | 0.04 ± 0.2 |
| 273 | *Canis lupus irremotus* | Northern Rocky Mountain Wolf | 1 | 0.04 ± 0.2 | 1 | 0.04 ± 0.2 |
| 274 | *Capricornis sumatraensis* | Mainland Serow | 1 | 0.04 ± 0.2 | 1 | 0.04 ± 0.2 |
| 275 | *Casarea dussumieri* | Keel-scaled Boa | 1 | 0.04 ± 0.2 | 1 | 0.04 ± 0.2 |
| 276 | *Centrochelys sulcata* | African Spurred Tortoise | 1 | 0.04 ± 0.2 | 1 | 0.04 ± 0.2 |
| 277 | *Cercocebus agilis* | Agile Mangabey | 1 | 0.04 ± 0.2 | 1 | 0.04 ± 0.2 |
| 278 | *Cercocebus galeritus* | Tana River Mangabey | 1 | 0.04 ± 0.2 | 1 | 0.04 ± 0.2 |
| 279 | *Cercocebus sanjei* | Sanje River Mangabey | 1 | 0.04 ± 0.2 | 1 | 0.04 ± 0.2 |
| 280 | *Cercopithecus ascanius* | Red-tailed Monkey | 1 | 0.04 ± 0.2 | 1 | 0.04 ± 0.2 |
| 281 | *Cercopithecus neglectus* | De Brazza's Monkey | 1 | 0.04 ± 0.2 | 1 | 0.04 ± 0.2 |
| 282 | *Cervus elaphus bactrianus (C. hanglu bactrianus)* | Bactrian Red Deer | 1 | 0.04 ± 0.2 | 1 | 0.04 ± 0.2 |
| 283 | *Chelonia mydas* | Green Turtle | 1 | 0.04 ± 0.2 | 1 | 0.04 ± 0.2 |
| 284 | *Chelydra serpentina* | Snapping Turtle | 1 | 0.04 ± 0.2 | 1 | 0.04 ± 0.2 |
| 285 | *Circus aeruginosus* | Western Marsh-harrier | 1 | 0.04 ± 0.2 | 1 | 0.04 ± 0.2 |
| 286 | *Colobus satanas* | Black Colobus | 1 | 0.04 ± 0.2 | 1 | 0.04 ± 0.2 |
| 287 | *Crocodylus cataphractus (Mecistops cataphractus)* | Slender-snouted Crocodile | 1 | 0.04 ± 0.2 | 1 | 0.04 ± 0.2 |
| 288 | *Dendrocygna arborea* | West Indian Whistling-duck | 1 | 0.04 ± 0.2 | 1 | 0.04 ± 0.2 |
| 289 | *Elanus leucurus* | White-tailed Kite | 1 | 0.04 ± 0.2 | 0 | 0 ± 0 |
| 290 | *Enhydra lutris* | Sea Otter | 1 | 0.04 ± 0.2 | 1 | 0.04 ± 0.2 |
| 291 | *Ephippiorhynchus senegalensis* | Saddlebill | 1 | 0.04 ± 0.2 | 1 | 0.04 ± 0.2 |
| 292 | *Eunectes murinus* | Green Anaconda | 1 | 0.04 ± 0.2 | 1 | 0.04 ± 0.2 |

| Rank | Scientific name | Common name | Trophies | Mean + SD trophies/year | WOEs | Mean + SD WOEs/year |
|---|---|---|---|---|---|---|
| 293 | *Eupodotis vigorsii (Heterotetrax vigorsii)* | Karoo Bustard | 1 | 0.04 ± 0.2 | 1 | 0.04 ± 0.2 |
| 294 | *Falco chicquera* | Red-headed Falcon | 1 | 0.04 ± 0.2 | 1 | 0.04 ± 0.2 |
| 295 | *Falco peregrinus* | Peregrine Falcon | 1 | 0.04 ± 0.2 | 0 | 0 ± 0 |
| 296 | *Falco sparverius* | American Kestrel | 1 | 0.04 ± 0.2 | 1 | 0.04 ± 0.2 |
| 297 | *Falco subbuteo* | Eurasian Hobby | 1 | 0.04 ± 0.2 | 1 | 0.04 ± 0.2 |
| 298 | *Felis chaus* | Jungle Cat | 1 | 0.04 ± 0.2 | 1 | 0.04 ± 0.2 |
| 299 | *Felis nigripes* | Black-footed Cat | 1 | 0.04 ± 0.2 | 1 | 0.04 ± 0.2 |
| 300 | *Gazella bennettii* | Chinkara | 1 | 0.04 ± 0.2 | 1 | 0.04 ± 0.2 |
| 301 | *Herpailurus yagouaroundi* | Jaguarundi | 1 | 0.04 ± 0.2 | 1 | 0.04 ± 0.2 |
| 302 | *Kinixys natalensis* | KwaZulu-Natal Hinged-back Tortoise | 1 | 0.04 ± 0.2 | 1 | 0.04 ± 0.2 |
| 303 | *Manis gigantea* | Giant Pangolin | 1 | 0.04 ± 0.2 | 1 | 0.04 ± 0.2 |
| 304 | *Melierax canorus* | Pale-Chanting Goshawk | 1 | 0.04 ± 0.2 | 1 | 0.04 ± 0.2 |
| 305 | *Melierax poliopterus* | Eastern Chanting-goshawk | 1 | 0.04 ± 0.2 | 1 | 0.04 ± 0.2 |
| 306 | *Micronisus gabar* | Gabar Goshawk | 1 | 0.04 ± 0.2 | 1 | 0.04 ± 0.2 |
| 307 | *Milvus migrans* | Black Kite | 1 | 0.04 ± 0.2 | 1 | 0.04 ± 0.2 |
| 308 | *Mustela erminea* | Stoat | 1 | 0.04 ± 0.2 | 1 | 0.04 ± 0.2 |
| 309 | *Mustela sibirica* | Siberian Weasel | 1 | 0.04 ± 0.2 | 1 | 0.04 ± 0.2 |
| 310 | *Myrmecophaga tridactyla* | Giant Anteater | 1 | 0.04 ± 0.2 | 1 | 0.04 ± 0.2 |
| 311 | *Naemorhedus goral* | Himalayan Goral | 1 | 0.04 ± 0.2 | 1 | 0.04 ± 0.2 |
| 312 | *Nettapus auritus* | African Pygmy-goose | 1 | 0.04 ± 0.2 | 1 | 0.04 ± 0.2 |
| 313 | *Nyctea scandiaca (Bubo scandiacus)* | Snowy Owl | 1 | 0.04 ± 0.2 | 1 | 0.04 ± 0.2 |
| 314 | *Odocoileus virginianus* | White-tailed Deer | 1 | 0.04 ± 0.2 | 1 | 0.04 ± 0.2 |
| 315 | *Otis tarda* | Great Bustard | 1 | 0.04 ± 0.2 | 1 | 0.04 ± 0.2 |
| 316 | *Otocolobus manul* | Manul | 1 | 0.04 ± 0.2 | 1 | 0.04 ± 0.2 |
| 317 | *Ovis hodgsoni (O. a. hodgonsi)* | Tibetan Argali | 1 | 0.04 ± 0.2 | 1 | 0.04 ± 0.2 |
| 318 | *Pantholops hodgsonii* | Chiru | 1 | 0.04 ± 0.2 | 1 | 0.04 ± 0.2 |
| 319 | *Parabuteo unicinctus* | Harris's Hawk | 1 | 0.04 ± 0.2 | 1 | 0.04 ± 0.2 |
| 320 | *Phoenicopterus roseus* | Greater Flamingo | 1 | 0.04 ± 0.2 | 1 | 0.04 ± 0.2 |
| 321 | *Phoenicopterus ruber* | American Flamingo | 1 | 0.04 ± 0.2 | 1 | 0.04 ± 0.2 |

| Rank | Scientific name | Common name | Trophies | Mean + SD trophies/year | WOEs | Mean + SD WOEs/year |
|---|---|---|---|---|---|---|
| 322 | *Polihierax semitorquatus* | African Pygmy-falcon | 1 | 0.04 ± 0.2 | 1 | 0.04 ± 0.2 |
| 323 | *Psittacus erithacus* | Grey Parrot | 1 | 0.04 ± 0.2 | 1 | 0.04 ± 0.2 |
| 324 | *Pteronetta hartlaubii* | Hartlaub's Duck | 1 | 0.04 ± 0.2 | 1 | 0.04 ± 0.2 |
| 325 | *Python natalensis* | Southern African Rock Python | 1 | 0.04 ± 0.2 | 1 | 0.04 ± 0.2 |
| 326 | *Rucervus duvaucelii* | Swamp Deer | 1 | 0.04 ± 0.2 | 1 | 0.04 ± 0.2 |
| 327 | *Rupicapra pyrenaica ornata* | Apennine Chamois | 1 | 0.04 ± 0.2 | 1 | 0.04 ± 0.2 |
| 328 | *Saguinus midas* | Golden-handed Tamarin | 1 | 0.04 ± 0.2 | 1 | 0.04 ± 0.2 |
| 329 | *Saiga tatarica* | Saiga | 1 | 0.04 ± 0.2 | 1 | 0.04 ± 0.2 |
| 330 | *Saimiri sciureus* | Guianan Squirrel Monkey | 1 | 0.04 ± 0.2 | 1 | 0.04 ± 0.2 |
| 331 | *Serinus mozambicus* | Yellow-fronted Canary | 1 | 0.04 ± 0.2 | 1 | 0.04 ± 0.2 |
| 332 | *Stephanoaetus coronatus* | Crowned Eagle | 1 | 0.04 ± 0.2 | 0 | 0 ± 0 |
| 333 | *Strix uralensis* | Ural Owl | 1 | 0.04 ± 0.2 | 1 | 0.04 ± 0.2 |
| 334 | *Tauraco corythaix* | Knysna Turaco | 1 | 0.04 ± 0.2 | 1 | 0.04 ± 0.2 |
| 335 | *Terrapene ornata* | Ornate Box Turtle | 1 | 0.04 ± 0.2 | 1 | 0.04 ± 0.2 |
| 336 | *Threskiornis aethiopicus* | African Sacred Ibis | 1 | 0.04 ± 0.2 | 1 | 0.04 ± 0.2 |
| 337 | *Ursus americanus emmonsii* | Emmon's Black Bear | 1 | 0.04 ± 0.2 | 1 | 0.04 ± 0.2 |
| 338 | *Varanus salvator* | Common Water Monitor | 1 | 0.04 ± 0.2 | 1 | 0.04 ± 0.2 |
| 339 | *Viverra civettina* | Malabar Civet | 1 | 0.04 ± 0.2 | 1 | 0.04 ± 0.2 |
| 340 | *Balearica regulorum* | Grey Crowned Crane | 1 | 0.04 ± 0.2 | 1 | 0.04 ± 0.2 |
| | | | **585,801** | | **447,443** | |

**Table S7. CITES-listed species traded as hunting trophies (2015-2024; *n* = 198), estimated number of trophies, mean (± SD) number of trophies traded/year, WOEs traded in this period, and mean (± SD) WOEs traded in this period/year, ranked by number of trophies**. Synonyms on the Red List are presented in parentheses with scientific names. For some species trophy hunting does not take place formally and/or exports are close to or exclusively from extra-limital/non-wild populations, indicated by an asterix (*).

| Rank | Scientific name | Common name | Total trophies | Mean + SD trophies/year | WOEs | Mean + SD WOEs/year |
|---|---|---|---|---|---|---|
| 1 | *Ursus americanus* | American Black Bear | 42030 | 4203 ± 1838.03 | 23681 | 2368.1 ± 1838.03 |
| 2 | *Crocodylus niloticus* | Nile Crocodile | 36100 | 3610 ± 4664.91 | 26022 | 2602.2 ± 4664.91 |
| 3 | *Loxodonta africana* | African Savanna Elephant | 14816 | 1481.6 ± 1803.74 | 3812.872 | 381.29 ± 1803.74 |
| 4 | *Equus zebra hartmannae* | Hartmann's Mountain Zebra | 14397 | 1439.7 ± 380.44 | 14221.25 | 1422.13 ± 380.44 |
| 5 | *Giraffa camelopardalis* | Giraffe | 12745 | 1274.5 ± 2250.04 | 5562 | 556.2 ± 2250.04 |
| 6 | *Hippopotamus amphibius* | Common Hippopotamus | 11882 | 1188.2 ± 298.21 | 4701.864 | 470.19 ± 298.21 |
| 7 | *Papio ursinus* | Chacma Baboon | 11679 | 1167.9 ± 320.89 | 11579.25 | 1157.93 ± 320.89 |
| 8 | *Canis lupus* | Grey Wolf | 5946 | 594.6 ± 194.55 | 5396 | 539.6 ± 194.55 |
| 9 | *Salvator rufescens* | Red Tegu | 5000 | 500 ± 1581.14 | 5000 | 500 ± 1581.14 |
| 10 | *Panthera pardus* | Leopard | 4832 | 483.2 ± 123.16 | 4691.5 | 469.15 ± 123.16 |
| 11 | *Caracal caracal* | Caracal | 3722 | 372.2 ± 153.62 | 3699 | 369.9 ± 153.62 |
| 12 | *Ursus arctos* | Brown Bear | 3611 | 361.1 ± 210.52 | 3439 | 343.9 ± 210.52 |
| 13 | *Meleagris ocellata* | Ocellated Turkey | 3483 | 348.3 ± 162.2 | 3364 | 336.4 ± 162.2 |
| 14 | *Chlorocebus pygerythrus* | Vervet Monkey | 3458 | 345.8 ± 166.55 | 3450 | 345 ± 166.55 |
| 15 | *Antilope cervicapra* | Blackbuck | 3250 | 325 ± 260.24 | 3249 | 324.9 ± 260.24 |
| 16 | *Capra sibirica* | Siberian Ibex | 3145 | 314.5 ± 158.69 | 3145 | 314.5 ± 158.69 |
| 17 | *Lynx canadensis* | Canada Lynx | 2397 | 239.7 ± 148.05 | 1879.75 | 187.98 ± 148.05 |
| 18 | *Kobus leche* | Lechwe | 1830 | 183 ± 53.13 | 1819 | 181.9 ± 53.13 |
| 19 | *Puma concolor* | Puma | 1669 | 166.9 ± 74.39 | 1642 | 164.2 ± 74.39 |
| 20 | *Ovis canadensis* | Bighorn Sheep | 1400 | 140 ± 35.94 | 1400 | 140 ± 35.94 |
| 21 | *Panthera leo* | Lion | 1357 | 135.7 ± 57.57 | 1191 | 119.1 ± 57.57 |

| | | | | | | |
|---|---|---|---|---|---|---|
| 22 | *Civettictis civetta* | African Civet | 1305 | 130.5 ± 48 | 1305 | 130.5 ± 48 |
| 23 | *Damaliscus pygargus pygargus* | Bontebok | 1293 | 129.3 ± 73.73 | 1293 | 129.3 ± 73.73 |
| 24 | *Ceratotherium simum simum* | Southern White Rhinoceros | 1254 | 125.4 ± 48.86 | 1219.75 | 121.98 ± 48.86 |
| 25 | *Leptailurus serval* | Serval | 1182 | 118.2 ± 51.13 | 1113.75 | 111.38 ± 51.13 |
| 26 | *Papio cynocephalus* | Yellow Baboon | 1174 | 117.4 ± 37.75 | 1173 | 117.3 ± 37.75 |
| 27 | *Capra hircus aegagrus* | Wild Goat | 1168 | 116.8 ± 51.1 | 1167 | 116.7 ± 51.1 |
| 28 | *Philantomba monticola* | Blue Duiker | 1168 | 116.8 ± 53.1 | 1168 | 116.8 ± 53.1 |
| 29 | *Alligator mississippiensis* | American Alligator | 858 | 85.8 ± 173.5 | 772 | 77.2 ± 173.5 |
| 30 | *Mellivora capensis* | Honey Badger | 851 | 85.1 ± 29.75 | 849 | 84.9 ± 29.75 |
| 31 | *Acinonyx jubatus* | Cheetah | 730 | 73 ± 22.15 | 723 | 72.3 ± 22.15 |
| 32 | *Crax rubra* | Great Curassow | 699 | 69.9 ± 36.73 | 692 | 69.2 ± 36.73 |
| 33 | *Ovis polii* | Marco Polo Argali | 653 | 65.3 ± 60.51 | 653 | 65.3 ± 60.51 |
| 34 | *Ovis ammon* | Wild Sheep | 632 | 63.2 ± 44.56 | 632 | 63.2 ± 44.56 |
| 35 | *Lontra canadensis* | North American River Otter | 624 | 62.4 ± 166.39 | 557 | 55.7 ± 166.39 |
| 36 | *Capra caucasica* | Western Tur | 512 | 51.2 ± 43.46 | 512 | 51.2 ± 43.46 |
| 37 | *Lynx rufus* | Bobcat | 415 | 41.5 ± 18.69 | 414 | 41.4 ± 18.69 |
| 38 | *Ursus maritimus* | Polar Bear | 414 | 41.4 ± 33.72 | 280.5 | 28.05 ± 33.72 |
| 39 | *Penelope purpurascens* | Crested Guan | 402 | 40.2 ± 23.51 | 401 | 40.1 ± 23.51 |
| 40 | *Proteles cristata* | Aardwolf | 401 | 40.1 ± 11.52 | 401 | 40.1 ± 11.52 |
| 41 | *Ammotragus lervia* | Aoudad | 400 | 40 ± 22.69 | 400 | 40 ± 22.69 |
| 42 | *Python brongersmai* | Brongersma's Short-tailed Python | 400 | 40 ± 126.49 | 400 | 40 ± 126.49 |
| 43 | *Felis lybica* | Afro-Asiatic Wildcat | 385 | 38.5 ± 22.96 | 384 | 38.4 ± 22.96 |
| 44 | *Papio anubis* | Olive Baboon | 375 | 37.5 ± 13.84 | 373.5 | 37.35 ± 13.84 |
| 45 | *Equus zebra* | Mountain Zebra | 357 | 35.7 ± 112.19 | 353 | 35.3 ± 112.19 |
| 46 | *Puma concolor couguar* | Florida Puma | 351 | 35.1 ± 79.67 | 279 | 27.9 ± 79.67 |

| | | | | | | |
|---|---|---|---|---|---|---|
| 47 | *Nasua narica* | White-nosed Coati | 342 | 34.2 ± 19.48 | 342 | 34.2 ± 19.48 |
| 48 | *Felis silvestris* | European Wildcat | 320 | 32 ± 20.1 | 317 | 31.7 ± 20.1 |
| 49 | *Stigmochelys pardalis* | Leopard Tortoise | 301 | 30.1 ± 94.83 | 1 | 0.1 ± 94.83 |
| 50 | *Odobenus rosmarus* | Walrus | 197 | 19.7 ± 15.22 | 67 | 6.7 ± 15.22 |
| 51 | *Damaliscus pygargus* | Blesbok | 177 | 17.7 ± 19.12 | 177 | 17.7 ± 19.12 |
| 52 | *Ovis darwini (Ovis ammon darwini)* | Gobi Argali | 171 | 17.1 ± 12.91 | 171 | 17.1 ± 12.91 |
| 53 | *Ortalis vetula* | Plain Chachalaca | 160 | 16 ± 9.72 | 160 | 16 ± 9.72 |
| 54 | *Canis aureus* | Common Jackal | 155 | 15.5 ± 5.36 | 155 | 15.5 ± 5.36 |
| 55 | *Dendrocygna autumnalis* | Black-bellied Whistling-duck | 139 | 13.9 ± 27.09 | 139 | 13.9 ± 27.09 |
| 56 | *Cephalophus dorsalis* | Bay Duiker | 124 | 12.4 ± 9.75 | 124 | 12.4 ± 9.75 |
| 57 | *Dendrocygna bicolor* | Fulvous Whistling-duck | 122 | 12.2 ± 25.6 | 122 | 12.2 ± 25.6 |
| 58 | *Colobus guereza* | Guereza | 117 | 11.7 ± 6.67 | 117 | 11.7 ± 6.67 |
| 59 | *Ovis bochariensis (Ovis v. bochariensis)* | Bukhara Urial | 107 | 10.7 ± 9.07 | 107 | 10.7 ± 9.07 |
| 60 | *Ceratotherium simum* | White Rhinoceros | 104 | 10.4 ± 6.8 | 97.25 | 9.73 ± 6.8 |
| 61 | *Malayopython reticulatus* | Reticulated Python | 100 | 10 ± 31.62 | 100 | 10 ± 31.62 |
| 62 | *Pseudois nayaur* | Blue Sheep | 98 | 9.8 ± 7.89 | 98 | 9.8 ± 7.89 |
| 63 | *Ovis cycloceros (O. v. cycloceros)* | Afghan Urial | 86 | 8.6 ± 18.73 | 86 | 8.6 ± 18.73 |
| 64 | *Capra falconeri* | Markhor | 83 | 8.3 ± 4.45 | 83 | 8.3 ± 4.45 |
| 65 | *Dasyprocta punctata* | Central American Agouti | 83 | 8.3 ± 6.02 | 83 | 8.3 ± 6.02 |
| 66 | *Equus zebra zebra* | Cape Mountain Zebra | 82 | 8.2 ± 11.19 | 79 | 7.9 ± 11.19 |
| 67 | *Capra falconeri heptneri* | Markhor | 79 | 7.9 ± 6.3 | 79 | 7.9 ± 6.3 |
| 68 | *Axis porcinus* | Hog Deer | 78 | 7.8 ± 7.21 | 78 | 7.8 ± 7.21 |
| 69 | *Papio hamadryas* | Hamadryas Baboon | 75 | 7.5 ± 4.53 | 75 | 7.5 ± 4.53 |
| 70 | *Cephalophus silvicultor* | Yellow-backed Duiker | 73 | 7.3 ± 4.42 | 73 | 7.3 ± 4.42 |
| 71 | *Capra hircus* | Goat | 64 | 6.4 ± 18.2 | 64 | 6.4 ± 18.2 |

| | | | | | | |
|---|---|---|---|---|---|---|
| 72 | *Ovis cycloceros cycloceros (O.v. cycloceros)* | Afghan Urial (subsp. cycloceros) | 63 | 6.3 ± 11.38 | 63 | 6.3 ± 11.38 |
| 73 | *Oryx dammah* | Scimitar-horned Oryx | 60 | 6 ± 5.56 | 60 | 6 ± 5.56 |
| 74 | *Ovis aries* | Urial | 55 | 5.5 ± 10.86 | 55 | 5.5 ± 10.86 |
| 75 | *Python sebae* | Central African Rock Python | 52 | 5.2 ± 4.39 | 52 | 5.2 ± 4.39 |
| 76 | *Ovis punjabiensis (Ovis v. punjabensis)* | Punjab Urial | 51 | 5.1 ± 6.87 | 51 | 5.1 ± 6.87 |
| 77 | *Sarkidiornis melanotos* | Knob-billed Duck | 51 | 5.1 ± 3.9 | 51 | 5.1 ± 3.9 |
| 78 | *Diceros bicornis* | Black Rhinoceros | 43 | 4.3 ± 3.09 | 43 | 4.3 ± 3.09 |
| 79 | *Philantomba maxwellii* | Maxwell's Duiker | 42 | 4.2 ± 4.37 | 42 | 4.2 ± 4.37 |
| 80 | *Lynx lynx* | Eurasian lynx | 41 | 4.1 ± 3.54 | 41 | 4.1 ± 3.54 |
| 81 | *Bubalus arnee* | Wild Water Buffalo | 35 | 3.5 ± 8.81 | 9 | 0.9 ± 8.81 |
| 82 | *Loxodonta cyclotis* | African Forest Elephant | 34 | 3.4 ± 5.06 | 23 | 2.3 ± 5.06 |
| 83 | *Hyaena hyaena* | Striped Hyena | 33 | 3.3 ± 3.33 | 33 | 3.3 ± 3.33 |
| 84 | *Theropithecus gelada* | Gelada | 30 | 3 ± 1.83 | 30 | 3 ± 1.83 |
| 85 | *Erythrocebus patas* | Patas Monkey | 29 | 2.9 ± 3.31 | 29 | 2.9 ± 3.31 |
| 86 | *Chlorocebus aethiops* | Grivet Monkey | 25 | 2.5 ± 4.28 | 25 | 2.5 ± 4.28 |
| 87 | *Arctocephalus pusillus* | Afro-Australian Fur Seal | 24 | 2.4 ± 4.06 | 19 | 1.9 ± 4.06 |
| 88 | *Cuniculus paca* | Paca | 18 | 1.8 ± 1.55 | 18 | 1.8 ± 1.55 |
| 89 | *Mazama temama* | Central American Red Brocket | 16 | 1.6 ± 4.72 | 16 | 1.6 ± 4.72 |
| 90 | *Canis lupus monstrabilis* | Texas Grey Wolf | 15 | 1.5 ± 3.44 | 15 | 1.5 ± 3.44 |
| 91 | *Ovis severtzovi (O.a. severtzovi)* | Severtzov's Argali | 15 | 1.5 ± 1.08 | 15 | 1.5 ± 1.08 |
| 92 | *Antigone canadensis (Grus canadensis)* | Sandhill Crane | 14 | 1.4 ± 1.84 | 14 | 1.4 ± 1.84 |
| 93 | *Bison bison athabascae* | Wood Bison | 14 | 1.4 ± 2.55 | 13 | 1.3 ± 2.55 |
| 94 | *Cercopithecus mitis* | Blue Monkey | 14 | 1.4 ± 1.35 | 14 | 1.4 ± 1.35 |

| | | | | | | |
|---|---|---|---|---|---|---|
| 95 | *Otolemur crassicaudatus* | Thick-tailed Greater Galago | 14 | 1.4 ± 1.78 | 14 | 1.4 ± 1.78 |
| 96 | *Ovis cycloceros arkal (O.v. arkal)* | Afghan Urial (subsp. arkal) | 14 | 1.4 ± 3.5 | 14 | 1.4 ± 3.5 |
| 97 | *Cephalophus zebra* | Zebra Duiker | 13 | 1.3 ± 4.11 | 13 | 1.3 ± 4.11 |
| 98 | *Crocodylus porosus* | Saltwater Crocodile | 12 | 1.2 ± 1.48 | 12 | 1.2 ± 1.48 |
| 99 | *Necrosyrtes monachus* | Hooded Vulture | 12 | 1.2 ± 2.57 | 12 | 1.2 ± 2.57 |
| 100 | *Ursus thibetanus* | Asiatic Black Bear | 11 | 1.1 ± 2.47 | 11 | 1.1 ± 2.47 |
| 101 | *Varanus albigularis* | White-throated Monitor | 11 | 1.1 ± 1.97 | 11 | 1.1 ± 1.97 |
| 102 | *Papio papio* | Guinea Baboon | 8 | 0.8 ± 2.53 | 8 | 0.8 ± 2.53 |
| 103 | *Tauraco livingstonii* | Livingstone's Turaco | 8 | 0.8 ± 2.53 | 8 | 0.8 ± 2.53 |
| 104 | *Haliaeetus vocifer* | African Fish-eagle | 7 | 0.7 ± 2.21 | 7 | 0.7 ± 2.21 |
| 105 | *Ovis vignei* | Urial | 7 | 0.7 ± 1.49 | 7 | 0.7 ± 1.49 |
| 106 | *Athene cunicularia* | Burrowing Owl | 6 | 0.6 ± 1.9 | 6 | 0.6 ± 1.9 |
| 107 | *Bison bison* | American Bison | 6 | 0.6 ± 1.9 | 5 | 0.5 ± 1.9 |
| 108 | *Cervus elaphus* | Red Deer | 6 | 0.6 ± 1.9 | 6 | 0.6 ± 1.9 |
| 109 | *Circaetus fasciolatus* | Southern Banded Snake-eagle | 6 | 0.6 ± 1.9 | 6 | 0.6 ± 1.9 |
| 110 | *Galago moholi* | Southern Lesser Galago | 6 | 0.6 ± 1.26 | 6 | 0.6 ± 1.26 |
| 111 | *Pavo cristatus* | Indian Peafowl | 6 | 0.6 ± 0.97 | 6 | 0.6 ± 0.97 |
| 112 | *Strix woodfordii* | African Wood-owl | 6 | 0.6 ± 1.9 | 6 | 0.6 ± 1.9 |
| 113 | *Capra falconeri megaceros* | Straight-horned Markhor | 5 | 0.5 ± 0.97 | 5 | 0.5 ± 0.97 |
| 114 | *Cephalophus ogilbyi* | Ogilby's Duiker | 5 | 0.5 ± 1.08 | 5 | 0.5 ± 1.08 |
| 115 | *Chlorocebus tantalus* | Tantalus Monkey | 5 | 0.5 ± 1.58 | 5 | 0.5 ± 1.58 |
| 116 | *Glaucidium capense* | African Barred Owlet | 5 | 0.5 ± 1.58 | 5 | 0.5 ± 1.58 |
| 117 | *Gyps rueppellii* | Rüppell's Vulture | 5 | 0.5 ± 0.97 | 5 | 0.5 ± 0.97 |
| 118 | *Moschus moschiferus* | Siberian Musk Deer | 5 | 0.5 ± 0.97 | 5 | 0.5 ± 0.97 |
| 119 | *Poicephalus cryptoxanthus* | Brown-headed Parrot | 5 | 0.5 ± 1.58 | 5 | 0.5 ± 1.58 |

| | | | | | | |
|---|---|---|---|---|---|---|
| 120 | *Tauraco porphyreolophus* | Purple-crested Turaco | 5 | 0.5 ± 0.85 | 5 | 0.5 ± 0.85 |
| 121 | *Accipiter gentilis* | Northern Goshawk | 4 | 0.4 ± 1.26 | 4 | 0.4 ± 1.26 |
| 122 | *Hystrix cristata* | Crested Porcupine | 4 | 0.4 ± 0.7 | 4 | 0.4 ± 0.7 |
| 123 | *Oryx leucoryx* | Arabian Oryx | 4 | 0.4 ± 0.84 | 4 | 0.4 ± 0.84 |
| 124 | *Ovis gmelini* | Mouflon | 4 | 0.4 ± 0.7 | 4 | 0.4 ± 0.7 |
| 125 | *Sagittarius serpentarius* | Secretarybird | 4 | 0.4 ± 0.7 | 4 | 0.4 ± 0.7 |
| 126 | *Trigonoceps occipitalis* | White-headed Vulture | 4 | 0.4 ± 1.26 | 4 | 0.4 ± 1.26 |
| 127 | *Aquila chrysaetos* | Golden Eagle | 3 | 0.3 ± 0.95 | 3 | 0.3 ± 0.95 |
| 128 | *Ardeotis kori* | Kori Bustard | 3 | 0.3 ± 0.48 | 3 | 0.3 ± 0.48 |
| 129 | *Gyps africanus* | White-backed Vulture | 3 | 0.3 ± 0.95 | 3 | 0.3 ± 0.95 |
| 130 | *Macrochelys temminckii* | Alligator Snapping Turtle | 3 | 0.3 ± 0.95 | 3 | 0.3 ± 0.95 |
| 131 | *Neotis denhami* | Denham's Bustard | 3 | 0.3 ± 0.67 | 3 | 0.3 ± 0.67 |
| 132 | *Phoeniconaias minor* | Lesser Flamingo | 3 | 0.3 ± 0.95 | 3 | 0.3 ± 0.95 |
| 133 | *Poicephalus fuscicollis* | Brown-necked Parrot | 3 | 0.3 ± 0.95 | 3 | 0.3 ± 0.95 |
| 134 | *Sarcogyps calvus* | Red-headed Vulture | 3 | 0.3 ± 0.95 | 3 | 0.3 ± 0.95 |
| 135 | *Ursus arctos isabellinus* | Himalayan Brown Bear | 3 | 0.3 ± 0.67 | 3 | 0.3 ± 0.67 |
| 136 | *Apalone mutica* | Smooth Softshell Turtle | 2 | 0.2 ± 0.63 | 2 | 0.2 ± 0.63 |
| 137 | *Boselaphus tragocamelus* | Nilgai | 2 | 0.2 ± 0.42 | 2 | 0.2 ± 0.42 |
| 138 | *Buteo augur* | Augur Buzzard | 2 | 0.2 ± 0.63 | 2 | 0.2 ± 0.63 |
| 139 | *Caiman crocodilus* | Spectacled Caiman | 2 | 0.2 ± 0.63 | 2 | 0.2 ± 0.63 |
| 140 | *Chlorocebus sabaeus* | Green Monkey | 2 | 0.2 ± 0.63 | 2 | 0.2 ± 0.63 |
| 141 | *Ciconia nigra* | Black Stork | 2 | 0.2 ± 0.63 | 2 | 0.2 ± 0.63 |
| 142 | *Colibri delphinae* | Brown Violetear | 2 | 0.2 ± 0.63 | 2 | 0.2 ± 0.63 |
| 143 | *Galago senegalensis* | Northern Lesser Galago | 2 | 0.2 ± 0.63 | 2 | 0.2 ± 0.63 |
| 144 | *Gypohierax angolensis* | Palm-nut Vulture | 2 | 0.2 ± 0.63 | 2 | 0.2 ± 0.63 |
| 145 | *Hippotragus niger* | Sable Antelope | 2 | 0.2 ± 0.63 | 1 | 0.1 ± 0.63 |

| | | | | | | |
|---|---|---|---|---|---|---|
| 146 | *Hydrictis maculicollis* | Spotted-necked Otter | 2 | 0.2 ± 0.63 | 2 | 0.2 ± 0.63 |
| 147 | *Lutra lutra* | Eurasian Otter | 2 | 0.2 ± 0.63 | 2 | 0.2 ± 0.63 |
| 148 | *Miopithecus talapoin* | Southern Talapoin Monkey | 2 | 0.2 ± 0.63 | 2 | 0.2 ± 0.63 |
| 149 | *Polemaetus bellicosus* | Martial Eagle | 2 | 0.2 ± 0.42 | 2 | 0.2 ± 0.42 |
| 150 | *Polyboroides typus* | African Harrier-hawk | 2 | 0.2 ± 0.63 | 2 | 0.2 ± 0.63 |
| 151 | *Syrmaticus reevesii* | Reeves's Pheasant | 2 | 0.2 ± 0.63 | 2 | 0.2 ± 0.63 |
| 152 | *Terathopius ecaudatus* | Bateleur | 2 | 0.2 ± 0.63 | 2 | 0.2 ± 0.63 |
| 153 | *Torgos tracheliotus* | Lappet-faced Vulture | 2 | 0.2 ± 0.42 | 2 | 0.2 ± 0.42 |
| 154 | *Varanus niloticus* | Nile Monitor | 2 | 0.2 ± 0.42 | 2 | 0.2 ± 0.42 |
| 155 | *Accipiter melanoleucus* | Black Sparrowhawk | 1 | 0.1 ± 0.32 | 1 | 0.1 ± 0.32 |
| 156 | *Accipiter striatus* | Sharp-shinned Hawk | 1 | 0.1 ± 0.32 | 1 | 0.1 ± 0.32 |
| 157 | *Aonyx capensis* | African Clawless Otter | 1 | 0.1 ± 0.32 | 1 | 0.1 ± 0.32 |
| 158 | *Aquila rapax* | Tawny Eagle | 1 | 0.1 ± 0.32 | 1 | 0.1 ± 0.32 |
| 159 | *Ara chloropterus* | Red-and-green Macaw | 1 | 0.1 ± 0.32 | 1 | 0.1 ± 0.32 |
| 160 | *Asio capensis* | Marsh Owl | 1 | 0.1 ± 0.32 | 1 | 0.1 ± 0.32 |
| 161 | *Balearica regulorum* | Grey Crowned Crane | 1 | 0.1 ± 0.32 | 1 | 0.1 ± 0.32 |
| 162 | *Branta canadensis leucopareia* | Aleutian Canada Goose | 1 | 0.1 ± 0.32 | 1 | 0.1 ± 0.32 |
| 163 | *Buteo buteo* | Eurasian Buzzard | 1 | 0.1 ± 0.32 | 1 | 0.1 ± 0.32 |
| 164 | *Caiman crocodilus crocodilus* | South American Spectacled Caiman | 1 | 0.1 ± 0.32 | 1 | 0.1 ± 0.32 |
| 165 | *Canis lupus irremotus* | Northern Rocky Mountain Wolf | 1 | 0.1 ± 0.32 | 1 | 0.1 ± 0.32 |
| 166 | *Centrochelys sulcata* | African Spurred Tortoise | 1 | 0.1 ± 0.32 | 1 | 0.1 ± 0.32 |
| 167 | *Cercopithecus neglectus* | De Brazza's Monkey | 1 | 0.1 ± 0.32 | 1 | 0.1 ± 0.32 |
| 168 | *Cervus elaphus bactrianus (C. hanglu bactrianus)* | Bactrian Red Deer | 1 | 0.1 ± 0.32 | 1 | 0.1 ± 0.32 |
| 169 | *Chelonia mydas* | Green Turtle | 1 | 0.1 ± 0.32 | 1 | 0.1 ± 0.32 |

| | | | | | | |
|---|---|---|---|---|---|---|
| 170 | *Chelydra serpentina* | Snapping Turtle | 1 | 0.1 ± 0.32 | 1 | 0.1 ± 0.32 |
| 171 | *Circaetus pectoralis* | Black-chested Snake-eagle | 1 | 0.1 ± 0.32 | 1 | 0.1 ± 0.32 |
| 172 | *Crotalus durissus* | Cascabel Rattlesnake | 1 | 0.1 ± 0.32 | 1 | 0.1 ± 0.32 |
| 173 | *Eunectes murinus* | Green Anaconda | 1 | 0.1 ± 0.32 | 1 | 0.1 ± 0.32 |
| 174 | *Eupodotis senegalensis* | White-bellied Bustard | 1 | 0.1 ± 0.32 | 1 | 0.1 ± 0.32 |
| 175 | *Falco chicquera* | Red-headed Falcon | 1 | 0.1 ± 0.32 | 1 | 0.1 ± 0.32 |
| 176 | *Gazella bennettii* | Chinkara | 1 | 0.1 ± 0.32 | 1 | 0.1 ± 0.32 |
| 177 | *Haliaeetus leucocephalus* | Bald Eagle | 1 | 0.1 ± 0.32 | 0 | 0 ± 0.32 |
| 178 | *Lissotis melanogaster* | Black-bellied Bustard | 1 | 0.1 ± 0.32 | 1 | 0.1 ± 0.32 |
| 179 | *Lophaetus occipitalis* | Long-crested Eagle | 1 | 0.1 ± 0.32 | 1 | 0.1 ± 0.32 |
| 180 | *Manis tricuspis* | White-bellied Pangolin | 1 | 0.1 ± 0.32 | 1 | 0.1 ± 0.32 |
| 181 | *Melierax canorus* | Pale-Chanting Goshawk | 1 | 0.1 ± 0.32 | 1 | 0.1 ± 0.32 |
| 182 | *Melierax poliopterus* | Eastern Chanting-goshawk | 1 | 0.1 ± 0.32 | 1 | 0.1 ± 0.32 |
| 183 | *Micronisus gabar* | Gabar Goshawk | 1 | 0.1 ± 0.32 | 1 | 0.1 ± 0.32 |
| 184 | *Mustela erminea* | Stoat | 1 | 0.1 ± 0.32 | 1 | 0.1 ± 0.32 |
| 185 | *Odocoileus virginianus* | White-tailed Deer | 1 | 0.1 ± 0.32 | 1 | 0.1 ± 0.32 |
| 186 | *Phoenicopterus roseus* | Greater Flamingo | 1 | 0.1 ± 0.32 | 1 | 0.1 ± 0.32 |
| 187 | *Phoenicopterus ruber* | American Flamingo | 1 | 0.1 ± 0.32 | 1 | 0.1 ± 0.32 |
| 188 | *Polihierax semitorquatus* | African Pygmy-falcon | 1 | 0.1 ± 0.32 | 1 | 0.1 ± 0.32 |
| 189 | *Python natalensis* | Southern African Rock Python | 1 | 0.1 ± 0.32 | 1 | 0.1 ± 0.32 |
| 190 | *Rucervus duvaucelii* | Swamp Deer | 1 | 0.1 ± 0.32 | 1 | 0.1 ± 0.32 |
| 191 | *Saguinus midas* | Golden-handed Tamarin | 1 | 0.1 ± 0.32 | 1 | 0.1 ± 0.32 |
| 192 | *Saimiri sciureus* | Guianan Squirrel Monkey | 1 | 0.1 ± 0.32 | 1 | 0.1 ± 0.32 |
| 193 | *Stephanoaetus coronatus* | Crowned Eagle | 1 | 0.1 ± 0.32 | 0 | 0 ± 0.32 |
| 194 | *Tauraco macrorhynchus* | Yellow-billed Turaco | 1 | 0.1 ± 0.32 | 1 | 0.1 ± 0.32 |

| | | | | | | |
|---|---|---|---|---|---|---|
| 195 | *Ursus americanus emmonsii* | Emmon's Black Bear | 1 | 0.1 ± 0.32 | 1 | 0.1 ± 0.32 |
| 196 | *Varanus salvator* | Common Water Monitor | 1 | 0.1 ± 0.32 | 1 | 0.1 ± 0.32 |
| 197 | *Viverra civettina* | Malabar Civet | 1 | 0.1 ± 0.32 | 1 | 0.1 ± 0.32 |
| 198 | *Vulpes vulpes* | Red Fox | 1 | 0.1 ± 0.32 | 1 | 0.1 ± 0.32 |
| **Totals** | | | **211,295** | | **154,514** | |

**Table S8. Estimated number of hunting trophies and WOEs from CITES-listed species imported by countries in the Global North (2015-2024) that have enacted or are considering import bans on hunting trophies, and other countries, ranked by number of trophies, and the number of trophies and WOEs that would be affected by proposed bans.**

| Country | Trophies | WOEs | Trophies affected | WOEs affected |
|---|---|---|---|---|
| United States | 102,999 | 73576.36 | 27196 | 22254.53 |
| Germany | 9733 | 8060.07 | 2209 | 1306.07 |
| Spain | 5297 | 4290.82 | 2289 | 1385.32 |
| France | 4110 | 2888.89 | 3879 | 2657.89 |
| Canada | 3349 | 2994.28 | 275 | 85.38 |
| Italy | 2408 | 1534.53 | 2299 | 1424.52 |
| Australia | 1666 | 1454 | 2 | 2 |
| Poland | 1514 | 1432.19 | 376 | 310 |
| United Kingdom | 1105 | 838.87 | 1033 | 766.87 |
| Belgium | 937 | 798.19 | 307 | 207.19 |
| Finland | 923 | 782 | 137 | 124.53 |
| Netherlands | 482 | 424.25 | 41 | 37 |
| TOTAL | 134,523 (64%) | 99,074.45 (64%) | 40043 (19%) | 30561.3 (31%) |
| Other countries | 76,772 (36%) | 55,439.55 (36%) | | |
| TOTAL | 211,295 | 154514 | | |

**Tables S9-S20. Species that are affected by enacted bans or would be affected by proposed import bans are presented in bold font. Trade in species in the period 2015-2024 that would be affected by these bans are highlighted in gray. Bans may not affect all trade in some species because different populations in are included in different Annexes of the EU Wildlife Trade Regulations.**

**Table S9. CITES-listed species imported to Australia as hunting trophies (2015-2024; *n* = 52), estimated number of trophies, mean (± SD) number of trophies imported/year, WOEs imported, and mean (± SD) WOEs imported/year, ranked by trophies.**

| Rank | Scientific name | Common name | Trophies | Mean + SD trophies/year | WOEs | Mean + SD WOEs/year |
|---|---|---|---|---|---|---|
| 1 | *Ursus americanus* | American Black Bear | 387 | 38.7 ± 21.55 | 293 | 29.3 ± 13.86 |
| 2 | *Papio ursinus* | Chacma Baboon | 173 | 17.3 ± 9.27 | 173 | 17.3 ± 9.27 |
| 3 | *Equus zebra hartmannae* | Hartmann's Mountain Zebra | 136 | 13.6 ± 6.54 | 134 | 13.4 ± 6.72 |
| 4 | *Chlorocebus pygerythrus* | Vervet Monkey | 126 | 12.6 ± 13.94 | 126 | 12.6 ± 13.94 |
| 5 | *Hippopotamus amphibius* | Common Hippopotamus | 120 | 12 ± 7.3 | 25.98 | 2.6 ± 2.05 |
| 6 | *Caracal caracal* | Caracal | 79 | 7.9 ± 3.84 | 79 | 7.9 ± 3.84 |
| 7 | *Giraffa camelopardalis* | Giraffe | 65 | 6.5 ± 5.91 | 63 | 6.3 ± 5.74 |
| 8 | *Ursus arctos* | Brown Bear | 62 | 6.2 ± 5.16 | 61 | 6.1 ± 5.22 |
| 9 | *Puma concolor* | Puma | 58 | 5.8 ± 4.8 | 57 | 5.7 ± 4.6 |
| 10 | *Capra sibirica* | Siberian Ibex | 55 | 5.5 ± 7.14 | 55 | 5.5 ± 7.14 |
| 11 | *Canis lupus* | Grey Wolf | 52 | 5.2 ± 3.55 | 47 | 4.7 ± 3.02 |
| 12 | *Antilope cervicapra* | Blackbuck | 37 | 3.7 ± 5.14 | 37 | 3.7 ± 5.14 |
| 13 | *Lynx canadensis* | Canada Lynx | 36 | 3.6 ± 4.45 | 36 | 3.6 ± 4.45 |
| 14 | *Crocodylus niloticus* | Nile Crocodile | 32 | 3.2 ± 2.66 | 32 | 3.2 ± 2.66 |
| 15 | *Loxodonta africana* | African Savanna Elephant | 26 | 2.6 ± 4.01 | 17.41 | 1.74 ± 2.55 |
| 16 | *Kobus leche* | Lechwe | 25 | 2.5 ± 3.21 | 25 | 2.5 ± 3.21 |
| 17 | *Damaliscus pygargus pygargus* | Bontebok | 19 | 1.9 ± 1.79 | 19 | 1.9 ± 1.79 |
| 18 | *Lynx rufus* | Bobcat | 18 | 1.8 ± 1.48 | 17 | 1.7 ± 1.42 |
| 19 | *Mellivora capensis* | Honey Badger | 17 | 1.7 ± 1.49 | 17 | 1.7 ± 1.49 |
| 20 | *Civettictis civetta* | African Civet | 15 | 1.5 ± 1.51 | 15 | 1.5 ± 1.51 |
| 21 | *Puma concolor couguar* | Florida Puma | 14 | 1.4 ± 3.78 | 14 | 1.4 ± 3.78 |
| 22 | *Ammotragus lervia* | Aoudad | 13 | 1.3 ± 1.42 | 13 | 1.3 ± 1.42 |
| 23 | *Ursus maritimus* | Polar Bear | 12 | 1.2 ± 1.32 | 9 | 0.9 ± 0.99 |
| 24 | *Capra hircus aegagrus (Capra aegagrus)* | Wild Goat | 11 | 1.1 ± 1.29 | 11 | 1.1 ± 1.29 |

| | | | | | | |
|---|---|---|---|---|---|---|
| x | *Leptailurus serval* | Serval | 10 | 1 ± 1.15 | 10 | 1 ± 1.15 |
| 26 | *Alligator mississippiensis* | American Alligator | 9 | 0.9 ± 1.29 | 9 | 0.9 ± 1.29 |
| 27 | *Philantomba monticola* | Blue Duiker | 9 | 0.9 ± 2.23 | 9 | 0.9 ± 2.23 |
| 28 | *Ceratotherium simum simum* | Southern White Rhinoceros | 8 | 0.8 ± 0.63 | 8 | 0.8 ± 0.63 |
| 29 | *Proteles cristata* | Aardwolf | 4 | 0.4 ± 0.52 | 4 | 0.4 ± 0.52 |
| 30 | *Axis porcinus* | Hog Deer | 3 | 0.3 ± 0.67 | 3 | 0.3 ± 0.67 |
| 31 | *Felis lybica* | Afro-Asiatic Wildcat | 3 | 0.3 ± 0.48 | 3 | 0.3 ± 0.48 |
| 32 | *Ovis ammon* | Wild Sheep | 3 | 0.3 ± 0.67 | 3 | 0.3 ± 0.67 |
| 33 | *Papio cynocephalus* | Yellow Baboon | 3 | 0.3 ± 0.48 | 3 | 0.3 ± 0.48 |
| 34 | *Chlorocebus aethiops* | Grivet Monkey | 2 | 0.2 ± 0.63 | 2 | 0.2 ± 0.63 |
| 35 | *Damaliscus pygargus* | Blesbok | 2 | 0.2 ± 0.42 | 2 | 0.2 ± 0.42 |
| 36 | *Equus zebra zebra* | Cape Mountain Zebra | 2 | 0.2 ± 0.63 | 2 | 0.2 ± 0.63 |
| 37 | *Otolemur crassicaudatus* | Thick-tailed Greater Galago | 2 | 0.2 ± 0.63 | 2 | 0.2 ± 0.63 |
| 38 | *Ovis polii* | Marco Polo Argali | 2 | 0.2 ± 0.63 | 2 | 0.2 ± 0.63 |
| 39 | ***Panthera leo*** | **Lion** | 2 | 0.2 ± 0.42 | 2 | 0.2 ± 0.42 |
| 40 | *Panthera pardus* | Leopard | 2 | 0.2 ± 0.42 | 2 | 0.2 ± 0.42 |
| 41 | *Canis aureus* | Common Jackal | 1 | 0.1 ± 0.32 | 1 | 0.1 ± 0.32 |
| 42 | *Capra caucasica* | Western Tur | 1 | 0.1 ± 0.32 | 1 | 0.1 ± 0.32 |
| 43 | *Capra falconeri* | Markhor | 1 | 0.1 ± 0.32 | 1 | 0.1 ± 0.32 |
| 44 | *Chelydra serpentina* | Snapping Turtle | 1 | 0.1 ± 0.32 | 1 | 0.1 ± 0.32 |
| 45 | *Colobus guereza* | Guereza | 1 | 0.1 ± 0.32 | 1 | 0.1 ± 0.32 |
| 46 | *Felis silvestris* | European Wildcat | 1 | 0.1 ± 0.32 | 1 | 0.1 ± 0.32 |
| 47 | *Lontra canadensis* | North American River Otter | 1 | 0.1 ± 0.32 | 1 | 0.1 ± 0.32 |
| 48 | *Oryx leucoryx* | Arabian Oryx | 1 | 0.1 ± 0.32 | 1 | 0.1 ± 0.32 |
| 49 | *Ovis aries* | Urial | 1 | 0.1 ± 0.32 | 1 | 0.1 ± 0.32 |
| 50 | *Papio anubis* | Olive Baboon | 1 | 0.1 ± 0.32 | 1 | 0.1 ± 0.32 |
| 51 | *Pseudois nayaur* | Blue Sheep | 1 | 0.1 ± 0.32 | 1 | 0.1 ± 0.32 |
| 52 | *Ursus americanus emmonsii* | Emmon's Black Bear | 1 | 0.1 ± 0.32 | 1 | 0.1 ± 0.32 |
| **Totals** | | | **1666** | | **1454** | |

**Table S10. CITES-listed species imported to Belgium as hunting trophies (2015-2024; *n* = 58), estimated number of trophies, mean (± SD) number of trophies imported/year, WOEs imported, and mean (± SD) WOEs imported/year, ranked by trophies. Asterix indicates populations of species split between Annex A and B of the EU WTRs.**

| Rank | Scientific name | Common name | Trophies | Mean + SD trophies/year | WOEs | Mean + SD WOEs/year |
|---|---|---|---|---|---|---|
| 1 | *Equus zebra hartmannae* | Hartmann's Mountain Zebra | 132 | 13.2 ± 5.18 | 132 | 13.2 ± 5.18 |
| 2 | ***Loxodonta africana**** | **African Savanna Elephant** | 93 | 9.3 ± 10.58 | 47.94 | 4.79 ± 4.68 |
| 3 | ***Hippopotamus amphibius*** | **Common Hippopotamus** | 91 | 9.1 ± 11.39 | 45.23 | 4.52 ± 5.58 |
| 4 | *Ursus americanus* | American Black Bear | 71 | 7.1 ± 8.29 | 43 | 4.3 ± 4.69 |
| 5 | *Antilope cervicapra* | Blackbuck | 63 | 6.3 ± 7.89 | 63 | 6.3 ± 7.89 |
| 6 | *Papio ursinus* | Chacma Baboon | 51 | 5.1 ± 2.77 | 51 | 5.1 ± 2.77 |
| 7 | *Capra sibirica* | Siberian Ibex | 43 | 4.3 ± 3.56 | 43 | 4.3 ± 3.56 |
| 8 | *Giraffa camelopardalis* | Giraffe | 33 | 3.3 ± 4.57 | 32 | 3.2 ± 4.54 |
| 9 | *Chlorocebus pygerythrus* | Vervet Monkey | 29 | 2.9 ± 2.92 | 29 | 2.9 ± 2.92 |
| 10 | ***Crocodylus niloticus**** | **Nile Crocodile** | 29 | 2.9 ± 1.37 | 29 | 2.9 ± 1.37 |
| 11 | ***Ursus arctos*** | **Brown Bear** | 28 | 2.8 ± 1.99 | 28 | 2.8 ± 1.99 |
| 12 | ***Panthera pardus*** | **Leopard** | 27 | 2.7 ± 1.89 | 27 | 2.7 ± 1.89 |
| 13 | ***Panthera leo**** | **Lion** | 26 | 2.6 ± 2.72 | 24 | 2.4 ± 2.22 |
| 14 | ***Caracal caracal**** | **Caracal** | 24 | 2.4 ± 1.35 | 24 | 2.4 ± 1.35 |
| 15 | *Capra hircus aegagrus (Capra aegagrus)* | Wild Goat | 19 | 1.9 ± 2.08 | 19 | 1.9 ± 2.08 |
| 16 | *Odobenus rosmarus* | Walrus | 14 | 1.4 ± 4.43 | 4 | 0.4 ± 1.26 |
| 17 | *Civettictis civetta* | African Civet | 12 | 1.2 ± 1.62 | 12 | 1.2 ± 1.62 |
| 18 | *Leptailurus serval* | Serval | 12 | 1.2 ± 0.92 | 12 | 1.2 ± 0.92 |
| 19 | ***Ursus maritimus*** | **Polar Bear** | 12 | 1.2 ± 1.99 | 8 | 0.8 ± 1.4 |
| 20 | *Capra caucasica* | Western Tur | 10 | 1 ± 2.21 | 10 | 1 ± 2.21 |
| 21 | ***Acinonyx jubatus*** | **Cheetah** | 9 | 0.9 ± 1.1 | 9 | 0.9 ± 1.1 |
| 22 | *Papio cynocephalus* | Yellow Baboon | 9 | 0.9 ± 0.99 | 9 | 0.9 ± 0.99 |
| 23 | *Felis lybica* | Afro-Asiatic Wildcat | 7 | 0.7 ± 1.06 | 7 | 0.7 ± 1.06 |
| 24 | *Kobus leche* | Lechwe | 6 | 0.6 ± 1.26 | 6 | 0.6 ± 1.26 |
| 25 | *Mellivora capensis* | Honey Badger | 6 | 0.6 ± 0.97 | 6 | 0.6 ± 0.97 |
| 26 | *Papio anubis* | Olive Baboon | 6 | 0.6 ± 1.07 | 6 | 0.6 ± 1.07 |

| | | | | | | |
|---|---|---|---|---|---|---|
| 27 | *Philantomba monticola* | Blue Duiker | 6 | 0.6 ± 0.52 | 6 | 0.6 ± 0.52 |
| 28 | ***Canis lupus**** | **Grey Wolf** | 5 | 0.5 ± 1.27 | 2 | 0.2 ± 0.42 |
| 29 | *Damaliscus pygargus pygargus* | Bontebok | 5 | 0.5 ± 0.97 | 5 | 0.5 ± 0.97 |
| 30 | ***Puma concolor**** | **Puma** | 5 | 0.5 ± 1.08 | 5 | 0.5 ± 1.08 |
| 31 | *Cephalophus silvicultor* | Yellow-backed Duiker | 4 | 0.4 ± 0.52 | 4 | 0.4 ± 0.52 |
| 32 | ***Ceratotherium simum simum*** | **Southern White Rhinoceros** | 4 | 0.4 ± 0.52 | 4 | 0.4 ± 0.52 |
| 33 | *Lynx canadensis* | Canada Lynx | 4 | 0.4 ± 0.84 | 4 | 0.4 ± 0.84 |
| 34 | *Cephalophus dorsalis* | Bay Duiker | 3 | 0.3 ± 0.67 | 3 | 0.3 ± 0.67 |
| 35 | *Equus zebra zebra* | Cape Mountain Zebra | 3 | 0.3 ± 0.95 | 3 | 0.3 ± 0.95 |
| 36 | ***Felis silvestris*** | **European Wildcat** | 3 | 0.3 ± 0.67 | 3 | 0.3 ± 0.67 |
| 37 | *Lynx rufus* | Bobcat | 3 | 0.3 ± 0.67 | 3 | 0.3 ± 0.67 |
| 38 | ***Ovis ammon*** | **Wild Sheep** | 3 | 0.3 ± 0.67 | 3 | 0.3 ± 0.67 |
| 39 | *Pseudois nayaur* | Blue Sheep | 3 | 0.3 ± 0.48 | 3 | 0.3 ± 0.48 |
| 40 | *Colobus guereza* | Guereza | 2 | 0.2 ± 0.42 | 2 | 0.2 ± 0.42 |
| 41 | *Erythrocebus patas* | Patas Monkey | 2 | 0.2 ± 0.42 | 2 | 0.2 ± 0.42 |
| 42 | *Gyps rueppellii* | Rüppell's Vulture | 2 | 0.2 ± 0.42 | 2 | 0.2 ± 0.42 |
| 43 | ***Oryx dammah*** | **Scimitar-horned Oryx** | 2 | 0.2 ± 0.42 | 2 | 0.2 ± 0.42 |
| 44 | *Ovis cycloceros (O. v. cycloceros)* | Afghan Urial | 2 | 0.2 ± 0.42 | 2 | 0.2 ± 0.42 |
| 45 | *Alligator mississippiensis* | American Alligator | 1 | 0.1 ± 0.32 | 1 | 0.1 ± 0.32 |
| 46 | *Canis aureus* | Common Jackal | 1 | 0.1 ± 0.32 | 1 | 0.1 ± 0.32 |
| 47 | ***Capra falconeri*** | **Markhor** | 1 | 0.1 ± 0.32 | 1 | 0.1 ± 0.32 |
| 48 | *Crax rubra* | Great Curassow | 1 | 0.1 ± 0.32 | 1 | 0.1 ± 0.32 |
| 49 | *Hyaena hyaena* | Striped Hyena | 1 | 0.1 ± 0.32 | 1 | 0.1 ± 0.32 |
| 50 | ***Hystrix cristata*** | **Crested Porcupine** | 1 | 0.1 ± 0.32 | 1 | 0.1 ± 0.32 |
| 51 | *Meleagris ocellata* | Ocellated Turkey | 1 | 0.1 ± 0.32 | 1 | 0.1 ± 0.32 |
| 52 | *Ovis aries* | Urial | 1 | 0.1 ± 0.32 | 1 | 0.1 ± 0.32 |
| 53 | *Ovis bochariensis (Ovis v. bochariensis)* | Bukhara Urial | 1 | 0.1 ± 0.32 | 1 | 0.1 ± 0.32 |
| 54 | ***Ovis darwini (Ovis ammon darwini)*** | **Gobi Argali** | 1 | 0.1 ± 0.32 | 1 | 0.1 ± 0.32 |
| 55 | ***Ovis polii*** | **Marco Polo Argali** | 1 | 0.1 ± 0.32 | 1 | 0.1 ± 0.32 |
| 56 | *Ovis punjabiensis (Ovis v. punjabensis)* | Punjab Urial | 1 | 0.1 ± 0.32 | 1 | 0.1 ± 0.32 |
| 57 | *Proteles cristata* | Aardwolf | 1 | 0.1 ± 0.32 | 1 | 0.1 ± 0.32 |

| | | | | | | |
|---|---|---|---|---|---|---|
| 58 | *Varanus niloticus* | Nile Monitor | 1 | 0.1 ± 0.32 | 1 | 0.1 ± 0.32 |
| **Totals** | | | **937** | | **798.19** | |

**Table S11. CITES-listed species imported to Canada as hunting trophies (2015-2024; *n* = 65), estimated number of trophies, mean (± SD) number of trophies imported/year, WOEs imported, and mean (± SD) WOEs imported/year, ranked by trophies.**

| Rank | Scientific name | Common name | Trophies | Mean + SD trophies/year | WOEs | Mean + SD WOEs/year |
|---|---|---|---|---|---|---|
| 1 | *Alligator mississippiensis* | American Alligator | 735 | 73.5 ± 170.94 | 670 | 67 ± 169.64 |
| 2 | *Equus zebra hartmannae* | Hartmann's Mountain Zebra | 287 | 28.7 ± 15.24 | 281 | 28.1 ± 15.71 |
| 3 | *Papio ursinus* | Chacma Baboon | 271 | 27.1 ± 12.48 | 271 | 27.1 ± 12.48 |
| 4 | ***Loxodonta africana*** | **African Savanna Elephant** | 243 | 24.3 ± 32.81 | 56.38 | 5.64 ± 3.36 |
| 5 | *Hippopotamus amphibius* | Common Hippopotamus | 154 | 15.4 ± 17.17 | 77.89 | 7.79 ± 4.95 |
| 6 | *Antilope cervicapra* | Blackbuck | 146 | 14.6 ± 13.38 | 146 | 14.6 ± 13.38 |
| 7 | *Ursus arctos* | Brown Bear | 127 | 12.7 ± 8.55 | 121 | 12.1 ± 8.54 |
| 8 | *Ovis polii* | Marco Polo Argali | 120 | 12 ± 21.63 | 120 | 12 ± 21.63 |
| 9 | *Giraffa camelopardalis* | Giraffe | 118 | 11.8 ± 14.86 | 114 | 11.4 ± 14.8 |
| 10 | *Capra sibirica* | Siberian Ibex | 110 | 11 ± 6.93 | 110 | 11 ± 6.93 |
| 11 | *Caracal caracal* | Caracal | 92 | 9.2 ± 5.14 | 92 | 9.2 ± 5.14 |
| 12 | *Chlorocebus pygerythrus* | Vervet Monkey | 89 | 8.9 ± 5.78 | 89 | 8.9 ± 5.78 |
| 13 | *Ovis canadensis* | Bighorn Sheep | 89 | 8.9 ± 4.23 | 89 | 8.9 ± 4.23 |
| 14 | *Panthera pardus* | Leopard | 72 | 7.2 ± 2.97 | 71 | 7.1 ± 2.73 |
| 15 | *Crocodylus niloticus* | Nile Crocodile | 71 | 7.1 ± 3.31 | 71 | 7.1 ± 3.31 |
| 16 | *Ovis ammon* | Wild Sheep | 69 | 6.9 ± 7.45 | 69 | 6.9 ± 7.45 |
| 17 | *Ammotragus lervia* | Aoudad | 58 | 5.8 ± 7.45 | 58 | 5.8 ± 7.45 |
| 18 | *Ursus americanus* | American Black Bear | 46 | 4.6 ± 3.13 | 41 | 4.1 ± 3.21 |
| 19 | *Capra hircus aegagrus* | Capra hircus aegagrus (Capra aegagrus) | 40 | 4 ± 3.16 | 40 | 4 ± 3.16 |
| 20 | ***Ceratotherium simum simum*** | **Southern White Rhinoceros** | 32 | 3.2 ± 3.65 | 29 | 2.9 ± 3.31 |
| 21 | *Panthera leo* | Lion | 32 | 3.2 ± 2.04 | 32 | 3.2 ± 2.04 |
| 22 | *Damaliscus pygargus pygargus* | Bontebok | 30 | 3 ± 3.43 | 30 | 3 ± 3.43 |
| 23 | *Acinonyx jubatus* | Cheetah | 28 | 2.8 ± 2.2 | 28 | 2.8 ± 2.2 |
| 24 | *Kobus leche* | Lechwe | 26 | 2.6 ± 1.71 | 26 | 2.6 ± 1.71 |
| 25 | *Leptailurus serval* | Serval | 25 | 2.5 ± 1.18 | 25 | 2.5 ± 1.18 |
| 26 | *Civettictis civetta* | African Civet | 21 | 2.1 ± 1.73 | 21 | 2.1 ± 1.73 |
| 27 | *Philantomba monticola* | Blue Duiker | 19 | 1.9 ± 1.79 | 19 | 1.9 ± 1.79 |

| | | | | | | |
|---|---|---|---|---|---|---|
| 28 | *Meleagris ocellata* | Ocellated Turkey | 18 | 1.8 ± 1.87 | 18 | 1.8 ± 1.87 |
| 29 | *Mellivora capensis* | Honey Badger | 15 | 1.5 ± 2.01 | 15 | 1.5 ± 2.01 |
| 30 | *Canis lupus* | Grey Wolf | 13 | 1.3 ± 0.95 | 13 | 1.3 ± 0.95 |
| 31 | *Capra caucasica* | Western Tur | 13 | 1.3 ± 1.89 | 13 | 1.3 ± 1.89 |
| 32 | *Felis silvestris* | European Wildcat | 12 | 1.2 ± 1.32 | 12 | 1.2 ± 1.32 |
| 33 | *Ovis darwini (Ovis ammon darwini)* | Gobi Argali | 12 | 1.2 ± 2.82 | 12 | 1.2 ± 2.82 |
| 34 | *Proteles cristata* | Aardwolf | 10 | 1 ± 1.05 | 10 | 1 ± 1.05 |
| 35 | *Papio cynocephalus* | Yellow Baboon | 9 | 0.9 ± 1.73 | 9 | 0.9 ± 1.73 |
| 36 | *Equus zebra zebra* | Cape Mountain Zebra | 7 | 0.7 ± 1.89 | 7 | 0.7 ± 1.89 |
| 37 | *Puma concolor* | Puma | 7 | 0.7 ± 0.95 | 6 | 0.6 ± 0.97 |
| 38 | *Capra falconeri* | Markhor | 6 | 0.6 ± 0.7 | 6 | 0.6 ± 0.7 |
| 39 | *Capra falconeri heptneri* | Markhor | 6 | 0.6 ± 0.97 | 6 | 0.6 ± 0.97 |
| 40 | *Crax rubra* | Great Curassow | 6 | 0.6 ± 1.26 | 6 | 0.6 ± 1.26 |
| 41 | *Felis lybica* | Afro-Asiatic Wildcat | 6 | 0.6 ± 0.7 | 6 | 0.6 ± 0.7 |
| 42 | *Oryx dammah* | Scimitar-horned Oryx | 6 | 0.6 ± 1.07 | 6 | 0.6 ± 1.07 |
| 43 | *Pseudois nayaur* | Blue Sheep | 6 | 0.6 ± 0.7 | 6 | 0.6 ± 0.7 |
| 44 | *Ovis bochariensis (Ovis v. bochariensis)* | Bukhara Urial | 5 | 0.5 ± 0.71 | 5 | 0.5 ± 0.71 |
| 45 | *Penelope purpurascens* | Crested Guan | 5 | 0.5 ± 1.08 | 5 | 0.5 ± 1.08 |
| 46 | *Lynx rufus* | Bobcat | 4 | 0.4 ± 0.84 | 4 | 0.4 ± 0.84 |
| 47 | *Damaliscus pygargus* | Blesbok | 3 | 0.3 ± 0.95 | 3 | 0.3 ± 0.95 |
| 48 | *Macrochelys temminckii* | Alligator Snapping Turtle | 3 | 0.3 ± 0.95 | 3 | 0.3 ± 0.95 |
| 49 | *Ovis severtzovi (O.a. severtzovi)* | Severtzov's Argali | 3 | 0.3 ± 0.67 | 3 | 0.3 ± 0.67 |
| 50 | *Apalone mutica* | Smooth Softshell Turtle | 2 | 0.2 ± 0.63 | 2 | 0.2 ± 0.63 |
| 51 | *Canis aureus* | Common Jackal | 2 | 0.2 ± 0.42 | 2 | 0.2 ± 0.42 |
| 52 | *Colobus guereza* | Guereza | 2 | 0.2 ± 0.63 | 2 | 0.2 ± 0.63 |
| 53 | *Lontra canadensis* | North American River Otter | 2 | 0.2 ± 0.63 | 2 | 0.2 ± 0.63 |
| 54 | *Lynx canadensis* | Canada Lynx | 2 | 0.2 ± 0.63 | 1 | 0.1 ± 0.32 |
| 55 | *Nasua narica* | White-nosed Coati | 2 | 0.2 ± 0.63 | 2 | 0.2 ± 0.63 |
| 56 | *Neotis denhami* | Denham's Bustard | 2 | 0.2 ± 0.63 | 2 | 0.2 ± 0.63 |
| 57 | *Ovis aries* | Urial | 2 | 0.2 ± 0.42 | 2 | 0.2 ± 0.42 |
| 58 | *Canis lupus irremotus* | Northern Rocky Mountain Wolf | 1 | 0.1 ± 0.32 | 1 | 0.1 ± 0.32 |

| | | | | | | |
|---|---|---|---|---|---|---|
| 59 | *Ceratotherium simum* | White Rhinoceros | 1 | 0.1 ± 0.32 | 1 | 0.1 ± 0.32 |
| 60 | *Cuniculus paca* | Paca | 1 | 0.1 ± 0.32 | 1 | 0.1 ± 0.32 |
| 61 | *Dasyprocta punctata* | Central American Agouti | 1 | 0.1 ± 0.32 | 1 | 0.1 ± 0.32 |
| 62 | *Ortalis vetula* | Plain Chachalaca | 1 | 0.1 ± 0.32 | 1 | 0.1 ± 0.32 |
| 63 | *Ovis cycloceros (O. v. cycloceros)* | Afghan Urial | 1 | 0.1 ± 0.32 | 1 | 0.1 ± 0.32 |
| 64 | *Papio anubis* | Olive Baboon | 1 | 0.1 ± 0.32 | 1 | 0.1 ± 0.32 |
| 65 | *Papio hamadryas* | Hamadryas Baboon | 1 | 0.1 ± 0.32 | 1 | 0.1 ± 0.32 |
| **Totals** | | | **3349** | | **2994.28** | |

**Table S12. CITES-listed species imported to Finland as hunting trophies (2015-2024; *n* = 32), estimated number of trophies, mean (± SD) number of trophies imported/year, WOEs imported, and mean (± SD) WOEs imported/year, ranked by trophies. Asterix indicates populations of species split between Annex A and B of the EU WTRs.**

| Rank | Scientific name | Common name | Trophies | Mean + SD trophies/year | WOEs | Mean + SD WOEs/year |
|---|---|---|---|---|---|---|
| 1 | *Ursus americanus* | American Black Bear | 447 | 44.7 ± 28.96 | 320.5 | 32.05 ± 21.95 |
| 2 | *Equus zebra hartmannae* | Hartmann's Mountain Zebra | 111 | 11.1 ± 9.65 | 110 | 11 ± 9.53 |
| 3 | *Papio ursinus* | Chacma Baboon | 89 | 8.9 ± 5.51 | 89 | 8.9 ± 5.51 |
| 4 | ***Ursus arctos*** | **Brown Bear** | 75 | 7.5 ± 12.88 | 75 | 7.5 ± 12.88 |
| 5 | *Chlorocebus pygerythrus* | Vervet Monkey | 31 | 3.1 ± 3.78 | 31 | 3.1 ± 3.78 |
| 6 | ***Caracal caracal**** | **Caracal** | 29 | 2.9 ± 3.98 | 29 | 2.9 ± 3.98 |
| 7 | ***Hippopotamus amphibius*** | **Common Hippopotamus** | 24 | 2.4 ± 4.01 | 12 | 1.2 ± 2.2 |
| 8 | ***Crocodylus niloticus**** | **Nile Crocodile** | 18 | 1.8 ± 1.48 | 18 | 1.8 ± 1.48 |
| 9 | ***Canis lupus**** | **Grey Wolf** | 10 | 1 ± 1.49 | 10 | 1 ± 1.49 |
| 10 | *Giraffa camelopardalis* | Giraffe | 10 | 1 ± 1.33 | 9 | 0.9 ± 1.1 |
| 11 | ***Loxodonta africana**** | **African Savanna Elephant** | 10 | 1 ± 1.41 | 9.53 | 0.95 ± 1.38 |
| 12 | *Capra sibirica* | Siberian Ibex | 9 | 0.9 ± 1.52 | 9 | 0.9 ± 1.52 |
| 13 | ***Acinonyx jubatus*** | **Cheetah** | 7 | 0.7 ± 0.82 | 7 | 0.7 ± 0.82 |
| 14 | *Damaliscus pygargus pygargus* | Bontebok | 7 | 0.7 ± 1.34 | 7 | 0.7 ± 1.34 |
| 15 | ***Panthera pardus*** | **Leopard** | 7 | 0.7 ± 1.06 | 7 | 0.7 ± 1.06 |
| 16 | *Antilope cervicapra* | Blackbuck | 6 | 0.6 ± 0.97 | 6 | 0.6 ± 0.97 |
| 17 | *Papio anubis* | Olive Baboon | 5 | 0.5 ± 1.58 | 5 | 0.5 ± 1.58 |
| 18 | *Alligator mississippiensis* | American Alligator | 4 | 0.4 ± 0.7 | 4 | 0.4 ± 0.7 |
| 19 | *Ovis ammon* | Wild Sheep | 4 | 0.4 ± 0.97 | 4 | 0.4 ± 0.97 |
| 20 | *Philantomba monticola* | Blue Duiker | 4 | 0.4 ± 0.7 | 4 | 0.4 ± 0.7 |
| 21 | *Kobus leche* | Lechwe | 3 | 0.3 ± 0.67 | 3 | 0.3 ± 0.67 |
| 22 | *Capra caucasica* | Western Tur | 2 | 0.2 ± 0.42 | 2 | 0.2 ± 0.42 |
| 23 | ***Ceratotherium simum simum**** | **Southern White Rhinoceros** | 2 | 0.2 ± 0.42 | 2 | 0.2 ± 0.42 |
| 24 | *Capra hircus aegagrus (Capra aegagrus)* | Wild Goat | 1 | 0.1 ± 0.32 | 1 | 0.1 ± 0.32 |
| 25 | *Civettictis civetta* | African Civet | 1 | 0.1 ± 0.32 | 1 | 0.1 ± 0.32 |
| 26 | *Damaliscus pygargus* | Blesbok | 1 | 0.1 ± 0.32 | 1 | 0.1 ± 0.32 |

| | | | | | | |
|---|---|---|---|---|---|---|
| 27 | *Equus zebra zebra* | Cape Mountain Zebra | 1 | 0.1 ± 0.32 | 1 | 0.1 ± 0.32 |
| 28 | *Felis lybica* | Afro-Asiatic Wildcat | 1 | 0.1 ± 0.32 | 1 | 0.1 ± 0.32 |
| 29 | ***Felis silvestris*** | **European Wildcat** | 1 | 0.1 ± 0.32 | 1 | 0.1 ± 0.32 |
| 30 | *Leptailurus serval* | Serval | 1 | 0.1 ± 0.32 | 1 | 0.1 ± 0.32 |
| 31 | *Mellivora capensis* | Honey Badger | 1 | 0.1 ± 0.32 | 1 | 0.1 ± 0.32 |
| 32 | ***Panthera leo**** | **Lion** | 1 | 0.1 ± 0.32 | 1 | 0.1 ± 0.32 |
| **Totals** | | | **923** | | **782** | |

**Table S13. CITES-listed species imported to France as hunting trophies (2015-2024; *n* = 72), estimated number of trophies, mean (± SD) number of trophies imported/year, WOEs imported, and mean (± SD) WOEs imported/year, ranked by trophies.**

| Rank | Scientific name | Common name | Trophies | Mean + SD trophies/year | WOEs | Mean + SD WOEs/year |
|---|---|---|---|---|---|---|
| 1 | ***Ursus americanus*** | **American Black Bear** | 1028 | 102.8 ± 41.44 | 558.5 | 55.85 ± 26.14 |
| 2 | ***Hippopotamus amphibius*** | **Common Hippopotamus** | 672 | 67.2 ± 34.77 | 124.83 | 12.48 ± 4.01 |
| 3 | ***Equus zebra hartmannae*** | **Hartmann's Mountain Zebra** | 501 | 50.1 ± 19.38 | 494 | 49.4 ± 18.5 |
| 4 | ***Loxodonta africana*** | **African Savanna Elephant** | 284 | 28.4 ± 28.18 | 134.06 | 13.41 ± 5.04 |
| 5 | ***Papio ursinus*** | **Chacma Baboon** | 209 | 20.9 ± 6.66 | 201 | 20.1 ± 6.17 |
| 6 | ***Panthera pardus*** | **Leopard** | 181 | 18.1 ± 6.82 | 180 | 18 ± 6.82 |
| 7 | ***Crocodylus niloticus*** | **Nile Crocodile** | 137 | 13.7 ± 4.97 | 137 | 13.7 ± 4.97 |
| 8 | *Capra sibirica* | Siberian Ibex | 106 | 10.6 ± 6.95 | 106 | 10.6 ± 6.95 |
| 9 | ***Ursus arctos*** | **Brown Bear** | 102 | 10.2 ± 11.22 | 102 | 10.2 ± 11.22 |
| 10 | ***Acinonyx jubatus*** | **Cheetah** | 90 | 9 ± 4.57 | 88 | 8.8 ± 4.32 |
| 11 | ***Caracal caracal*** | **Caracal** | 80 | 8 ± 3.74 | 80 | 8 ± 3.74 |
| 12 | *Antilope cervicapra* | Blackbuck | 65 | 6.5 ± 8.09 | 65 | 6.5 ± 8.09 |
| 13 | ***Giraffa camelopardalis*** | **Giraffe** | 63 | 6.3 ± 7.07 | 58 | 5.8 ± 6.44 |
| 14 | ***Canis lupus*** | **Grey Wolf** | 62 | 6.2 ± 4.49 | 50 | 5 ± 4 |
| 15 | ***Papio anubis*** | **Olive Baboon** | 43 | 4.3 ± 3.02 | 41.5 | 4.15 ± 2.79 |
| 16 | ***Kobus leche*** | **Lechwe** | 38 | 3.8 ± 2.82 | 38 | 3.8 ± 2.82 |
| 17 | ***Chlorocebus pygerythrus*** | **Vervet Monkey** | 37 | 3.7 ± 2.63 | 37 | 3.7 ± 2.63 |
| 18 | ***Papio cynocephalus*** | **Yellow Baboon** | 35 | 3.5 ± 2.55 | 35 | 3.5 ± 2.55 |
| 19 | ***Puma concolor*** | **Puma** | 30 | 3 ± 3.68 | 29 | 2.9 ± 3.67 |
| 20 | *Capra hircus aegagrus (Capra aegagrus)* | Wild Goat | 29 | 2.9 ± 2.85 | 29 | 2.9 ± 2.85 |
| 21 | ***Ceratotherium simum simum*** | **Southern White Rhinoceros** | 29 | 2.9 ± 3.03 | 29 | 2.9 ± 3.03 |
| 22 | ***Damaliscus pygargus pygargus*** | **Bontebok** | 22 | 2.2 ± 3.01 | 22 | 2.2 ± 3.01 |
| 23 | ***Ovis polii*** | **Marco Polo Argali** | 22 | 2.2 ± 3.01 | 22 | 2.2 ± 3.01 |
| 24 | ***Philantomba monticola*** | **Blue Duiker** | 20 | 2 ± 2 | 20 | 2 ± 2 |
| 25 | ***Capra caucasica*** | **Western Tur** | 16 | 1.6 ± 2.5 | 16 | 1.6 ± 2.5 |
| 26 | ***Ammotragus lervia*** | **Aoudad** | 14 | 1.4 ± 2.55 | 14 | 1.4 ± 2.55 |
| 27 | *Civettictis civetta* | African Civet | 14 | 1.4 ± 1.26 | 14 | 1.4 ± 1.26 |

| | | | | | | |
|---|---|---|---|---|---|---|
| 28 | ***Ursus maritimus*** | **Polar Bear** | 14 | 1.4 ± 2.88 | 10 | 1 ± 1.94 |
| 29 | ***Cephalophus dorsalis*** | **Bay Duiker** | 13 | 1.3 ± 1.7 | 13 | 1.3 ± 1.7 |
| 30 | ***Ovis ammon*** | **Wild Sheep** | 13 | 1.3 ± 1.57 | 13 | 1.3 ± 1.57 |
| 31 | ***Panthera leo*** | **Lion** | 13 | 1.3 ± 3.43 | 13 | 1.3 ± 3.43 |
| 32 | ***Leptailurus serval*** | **Serval** | 12 | 1.2 ± 2.15 | 12 | 1.2 ± 2.15 |
| 33 | ***Loxodonta cyclotis*** | **African Forest Elephant** | 12 | 1.2 ± 3.79 | 1 | 0.1 ± 0.32 |
| 34 | ***Lynx canadensis*** | **Canada Lynx** | 10 | 1 ± 1.89 | 10 | 1 ± 1.89 |
| 35 | ***Felis silvestris*** | **European Wildcat** | 6 | 0.6 ± 1.35 | 6 | 0.6 ± 1.35 |
| 36 | ***Lynx rufus*** | **Bobcat** | 6 | 0.6 ± 0.97 | 6 | 0.6 ± 0.97 |
| 37 | *Mellivora capensis* | Honey Badger | 6 | 0.6 ± 0.7 | 6 | 0.6 ± 0.7 |
| 38 | ***Python sebae*** | **Central African Rock Python** | 6 | 0.6 ± 0.52 | 6 | 0.6 ± 0.52 |
| 39 | ***Cephalophus silvicultor*** | **Yellow-backed Duiker** | 5 | 0.5 ± 0.97 | 5 | 0.5 ± 0.97 |
| 40 | ***Ceratotherium simum*** | **White Rhinoceros** | 4 | 0.4 ± 0.7 | 4 | 0.4 ± 0.7 |
| 41 | ***Colobus guereza*** | **Guereza** | 4 | 0.4 ± 0.7 | 4 | 0.4 ± 0.7 |
| 42 | *Hyaena hyaena* | Striped Hyena | 4 | 0.4 ± 1.26 | 4 | 0.4 ± 1.26 |
| 43 | ***Lynx lynx*** | **Eurasian lynx** | 4 | 0.4 ± 0.84 | 4 | 0.4 ± 0.84 |
| 44 | *Canis aureus* | Common Jackal | 3 | 0.3 ± 0.67 | 3 | 0.3 ± 0.67 |
| 45 | ***Capra falconeri heptneri*** | **Markhor** | 3 | 0.3 ± 0.48 | 3 | 0.3 ± 0.48 |
| 46 | ***Ovis cycloceros (O. v. cycloceros)*** | **Afghan Urial** | 3 | 0.3 ± 0.95 | 3 | 0.3 ± 0.95 |
| 47 | ***Ovis cycloceros arkal (O.v. arkal)*** | **Afghan Urial (subsp. arkal)** | 3 | 0.3 ± 0.95 | 3 | 0.3 ± 0.95 |
| 48 | ***Papio hamadryas*** | **Hamadryas Baboon** | 3 | 0.3 ± 0.67 | 3 | 0.3 ± 0.67 |
| 49 | ***Alligator mississippiensis*** | **American Alligator** | 2 | 0.2 ± 0.42 | 2 | 0.2 ± 0.42 |
| 50 | ***Chlorocebus tantalus*** | **Tantalus Monkey** | 2 | 0.2 ± 0.63 | 2 | 0.2 ± 0.63 |
| 51 | ***Diceros bicornis*** | **Black Rhinoceros** | 2 | 0.2 ± 0.63 | 2 | 0.2 ± 0.63 |
| 52 | ***Erythrocebus patas*** | **Patas Monkey** | 2 | 0.2 ± 0.63 | 2 | 0.2 ± 0.63 |
| 53 | ***Felis lybica*** | **Afro-Asiatic Wildcat** | 2 | 0.2 ± 0.42 | 2 | 0.2 ± 0.42 |
| 54 | ***Miopithecus talapoin*** | **Southern Talapoin Monkey** | 2 | 0.2 ± 0.63 | 2 | 0.2 ± 0.63 |
| 55 | ***Moschus moschiferus*** | **Siberian Musk Deer** | 2 | 0.2 ± 0.63 | 2 | 0.2 ± 0.63 |
| 56 | ***Odobenus rosmarus*** | **Walrus** | 2 | 0.2 ± 0.63 | 1 | 0.1 ± 0.32 |
| 57 | *Pseudois nayaur* | Blue Sheep | 2 | 0.2 ± 0.42 | 2 | 0.2 ± 0.42 |
| 58 | ***Sarkidiornis melanotos*** | **Knob-billed Duck** | 2 | 0.2 ± 0.42 | 2 | 0.2 ± 0.42 |

| | | | | | | |
|---|---|---|---|---|---|---|
| 59 | ***Capra falconeri*** | **Markhor** | 1 | 0.1 ± 0.32 | 1 | 0.1 ± 0.32 |
| 60 | ***Centrochelys sulcata*** | **African Spurred Tortoise** | 1 | 0.1 ± 0.32 | 1 | 0.1 ± 0.32 |
| 61 | ***Chelonia mydas*** | **Green Turtle** | 1 | 0.1 ± 0.32 | 1 | 0.1 ± 0.32 |
| 62 | *Damaliscus pygargus* | Blesbok | 1 | 0.1 ± 0.32 | 1 | 0.1 ± 0.32 |
| 63 | ***Equus zebra zebra*** | **Cape Mountain Zebra** | 1 | 0.1 ± 0.32 | 1 | 0.1 ± 0.32 |
| 64 | ***Neotis denhami*** | **Denham's Bustard** | 1 | 0.1 ± 0.32 | 1 | 0.1 ± 0.32 |
| 65 | ***Ovis bochariensis (Ovis v. bochariensis)*** | **Bukhara Urial** | 1 | 0.1 ± 0.32 | 1 | 0.1 ± 0.32 |
| 66 | ***Ovis canadensis*** | **Bighorn Sheep** | 1 | 0.1 ± 0.32 | 1 | 0.1 ± 0.32 |
| 67 | ***Ovis darwini (Ovis ammon darwini)*** | **Gobi Argali** | 1 | 0.1 ± 0.32 | 1 | 0.1 ± 0.32 |
| 68 | *Proteles cristata* | Aardwolf | 1 | 0.1 ± 0.32 | 1 | 0.1 ± 0.32 |
| 69 | ***Puma concolor couguar*** | **Florida Puma** | 1 | 0.1 ± 0.32 | 1 | 0.1 ± 0.32 |
| 70 | ***Stephanoaetus coronatus*** | **Crowned Eagle** | 1 | 0.1 ± 0.32 | 0 | 0 ± 0 |
| 71 | ***Ursus arctos isabellinus*** | **Himalayan Brown Bear** | 1 | 0.1 ± 0.32 | 1 | 0.1 ± 0.32 |
| 72 | ***Varanus albigularis*** | **White-throated Monitor** | 1 | 0.1 ± 0.32 | 1 | 0.1 ± 0.32 |
| **Totals** | | | **4110** | | **2888.89** | |

**Table S14. CITES-listed species imported to Germany as hunting trophies (2015-2024; *n* = 73), estimated number of trophies, mean (± SD) number of trophies imported/year, WOEs imported, and mean (± SD) WOEs imported/year, ranked by trophies. Asterix indicates populations of species split between Annex A and B of the EU WTRs.**

| Rank | Scientific name | Common name | Trophies | Mean + SD trophies/year | WOEs | Mean + SD WOEs/year |
|---|---|---|---|---|---|---|
| 1 | *Equus zebra hartmannae* | Hartmann's Mountain Zebra | 2779 | 277.9 ± 69.97 | 2768.5 | 276.85 ± 69.97 |
| 2 | *Papio ursinus* | Chacma Baboon | 1674 | 167.4 ± 53.58 | 1640 | 164 ± 53.58 |
| 3 | *Ursus americanus* | American Black Bear | 861 | 86.1 ± 33.4 | 675 | 67.5 ± 33.4 |
| 4 | ***Crocodylus niloticus**** | **Nile Crocodile** | 748 | 74.8 ± 158.08 | 242.5 | 24.25 ± 158.08 |
| 5 | ***Loxodonta africana**** | **African Savanna Elephant** | 677 | 67.7 ± 63.71 | 242.75 | 24.28 ± 63.71 |
| 6 | ***Hippopotamus amphibius*** | **Common Hippopotamus** | 668 | 66.8 ± 32.86 | 245.32 | 24.53 ± 32.86 |
| 7 | *Giraffa camelopardalis* | Giraffe | 278 | 27.8 ± 29.59 | 262 | 26.2 ± 29.59 |
| 8 | *Capra sibirica* | Siberian Ibex | 252 | 25.2 ± 14.86 | 252 | 25.2 ± 14.86 |
| 9 | ***Panthera pardus*** | **Leopard** | 234 | 23.4 ± 7.78 | 232 | 23.2 ± 7.78 |
| 10 | ***Canis lupus**** | **Grey Wolf** | 184 | 18.4 ± 10.11 | 167 | 16.7 ± 10.11 |
| 11 | ***Caracal caracal**** | **Caracal** | 181 | 18.1 ± 6.4 | 180 | 18 ± 6.4 |
| 12 | ***Ursus arctos*** | **Brown Bear** | 142 | 14.2 ± 12.63 | 141 | 14.1 ± 12.63 |
| 13 | *Kobus leche* | Lechwe | 100 | 10 ± 5.31 | 99 | 9.9 ± 5.31 |
| 14 | ***Acinonyx jubatus*** | **Cheetah** | 78 | 7.8 ± 3.91 | 78 | 7.8 ± 3.91 |
| 15 | *Chlorocebus pygerythrus* | Vervet Monkey | 78 | 7.8 ± 6.37 | 77 | 7.7 ± 6.37 |
| 16 | *Antilope cervicapra* | Blackbuck | 76 | 7.6 ± 7.41 | 76 | 7.6 ± 7.41 |
| 17 | *Civettictis civetta* | African Civet | 61 | 6.1 ± 3.78 | 61 | 6.1 ± 3.78 |
| 18 | ***Puma concolor**** | **Puma** | 60 | 6 ± 6.83 | 59 | 5.9 ± 6.83 |
| 19 | ***Panthera leo**** | **Lion** | 50 | 5 ± 2.26 | 50 | 5 ± 2.26 |
| 20 | *Lynx canadensis* | Canada Lynx | 47 | 4.7 ± 3.43 | 47 | 4.7 ± 3.43 |
| 21 | ***Ovis polii*** | **Marco Polo Argali** | 39 | 3.9 ± 3.96 | 39 | 3.9 ± 3.96 |
| 22 | ***Ursus maritimus*** | **Polar Bear** | 39 | 3.9 ± 8.37 | 13 | 1.3 ± 8.37 |
| 23 | ***Ceratotherium simum simum**** | **Southern White Rhinoceros** | 38 | 3.8 ± 3.26 | 38 | 3.8 ± 3.26 |
| 24 | *Capra hircus aegagrus (Capra aegagrus)* | Wild Goat | 36 | 3.6 ± 3.89 | 36 | 3.6 ± 3.89 |
| 25 | *Leptailurus serval* | Serval | 32 | 3.2 ± 1.87 | 32 | 3.2 ± 1.87 |
| 26 | *Mellivora capensis* | Honey Badger | 29 | 2.9 ± 2.38 | 29 | 2.9 ± 2.38 |

| | | | | | | |
|---|---|---|---|---|---|---|
| 27 | *Capra caucasica* | Western Tur | 25 | 2.5 ± 4.01 | 25 | 2.5 ± 4.01 |
| 28 | *Philantomba monticola* | Blue Duiker | 25 | 2.5 ± 2.22 | 25 | 2.5 ± 2.22 |
| 29 | *Papio cynocephalus* | Yellow Baboon | 23 | 2.3 ± 1.89 | 23 | 2.3 ± 1.89 |
| 30 | *Odobenus rosmarus* | Walrus | 20 | 2 ± 3.89 | 6 | 0.6 ± 3.89 |
| 31 | ***Ovis ammon*** | **Wild Sheep** | 20 | 2 ± 2.75 | 20 | 2 ± 2.75 |
| 32 | *Damaliscus pygargus pygargus* | Bontebok | 15 | 1.5 ± 1.65 | 15 | 1.5 ± 1.65 |
| 33 | *Alligator mississippiensis* | American Alligator | 13 | 1.3 ± 1.06 | 13 | 1.3 ± 1.06 |
| 34 | *Papio anubis* | Olive Baboon | 11 | 1.1 ± 1.52 | 11 | 1.1 ± 1.52 |
| 35 | *Ammotragus lervia* | Aoudad | 10 | 1 ± 1.25 | 10 | 1 ± 1.25 |
| 36 | *Felis lybica* | Afro-Asiatic Wildcat | 10 | 1 ± 0.82 | 10 | 1 ± 0.82 |
| 37 | *Proteles cristata* | Aardwolf | 10 | 1 ± 1.25 | 10 | 1 ± 1.25 |
| 38 | ***Ovis darwini (Ovis ammon darwini)*** | **Gobi Argali** | 9 | 0.9 ± 1.6 | 9 | 0.9 ± 1.6 |
| 39 | *Puma concolor couguar* | Florida Puma | 8 | 0.8 ± 1.93 | 8 | 0.8 ± 1.93 |
| 40 | ***Felis silvestris*** | **European Wildcat** | 7 | 0.7 ± 1.49 | 7 | 0.7 ± 1.49 |
| 41 | *Cephalophus dorsalis* | Bay Duiker | 6 | 0.6 ± 0.97 | 6 | 0.6 ± 0.97 |
| 42 | ***Ceratotherium simum*** | **White Rhinoceros** | 5 | 0.5 ± 0.85 | 5 | 0.5 ± 0.85 |
| 43 | *Damaliscus pygargus* | Blesbok | 5 | 0.5 ± 0.97 | 5 | 0.5 ± 0.97 |
| 44 | ***Diceros bicornis*** | **Black Rhinoceros** | 5 | 0.5 ± 0.97 | 5 | 0.5 ± 0.97 |
| 45 | *Hyaena hyaena* | Striped Hyena | 5 | 0.5 ± 0.97 | 5 | 0.5 ± 0.97 |
| 46 | *Lynx rufus* | Bobcat | 5 | 0.5 ± 0.85 | 5 | 0.5 ± 0.85 |
| 47 | *Ovis bochariensis (Ovis v. bochariensis)* | Bukhara Urial | 5 | 0.5 ± 0.53 | 5 | 0.5 ± 0.53 |
| 48 | ***Capra falconeri*** | **Markhor** | 4 | 0.4 ± 0.7 | 4 | 0.4 ± 0.7 |
| 49 | *Pseudois nayaur* | Blue Sheep | 4 | 0.4 ± 0.7 | 4 | 0.4 ± 0.7 |
| 50 | ***Capra falconeri heptneri*** | **Markhor** | 3 | 0.3 ± 0.48 | 3 | 0.3 ± 0.48 |
| 51 | *Cephalophus silvicultor* | Yellow-backed Duiker | 3 | 0.3 ± 0.95 | 3 | 0.3 ± 0.95 |
| 52 | *Colobus guereza* | Guereza | 3 | 0.3 ± 0.48 | 3 | 0.3 ± 0.48 |
| 53 | *Equus zebra zebra* | Cape Mountain Zebra | 3 | 0.3 ± 0.67 | 3 | 0.3 ± 0.67 |
| 54 | ***Lynx lynx*** | **Eurasian lynx** | 3 | 0.3 ± 0.67 | 3 | 0.3 ± 0.67 |
| 55 | ***Ovis vignei*** | **Urial** | 3 | 0.3 ± 0.95 | 3 | 0.3 ± 0.95 |
| 56 | *Canis aureus* | Common Jackal | 2 | 0.2 ± 0.42 | 2 | 0.2 ± 0.42 |
| 57 | *Lontra canadensis* | North American River Otter | 2 | 0.2 ± 0.42 | 2 | 0.2 ± 0.42 |

| | | | | | | |
|---|---|---|---|---|---|---|
| 58 | *Meleagris ocellata* | Ocellated Turkey | 2 | 0.2 ± 0.63 | 2 | 0.2 ± 0.63 |
| 59 | *Ovis canadensis* | Bighorn Sheep | 2 | 0.2 ± 0.42 | 2 | 0.2 ± 0.42 |
| 60 | *Ovis cycloceros cycloceros (O.v. cycloceros)* | Afghan Urial (subsp. cycloceros) | 2 | 0.2 ± 0.42 | 2 | 0.2 ± 0.42 |
| 61 | *Ovis punjabiensis (Ovis v. punjabensis)* | Punjab Urial | 2 | 0.2 ± 0.42 | 2 | 0.2 ± 0.42 |
| 62 | *Ardeotis kori* | Kori Bustard | 1 | 0.1 ± 0.32 | 1 | 0.1 ± 0.32 |
| 63 | *Axis porcinus* | Hog Deer | 1 | 0.1 ± 0.32 | 1 | 0.1 ± 0.32 |
| 64 | *Cercopithecus mitis* | Blue Monkey | 1 | 0.1 ± 0.32 | 1 | 0.1 ± 0.32 |
| 65 | *Chlorocebus aethiops* | Grivet Monkey | 1 | 0.1 ± 0.32 | 1 | 0.1 ± 0.32 |
| 66 | *Erythrocebus patas* | Patas Monkey | 1 | 0.1 ± 0.32 | 1 | 0.1 ± 0.32 |
| 67 | ***Oryx dammah*** | **Scimitar-horned Oryx** | 1 | 0.1 ± 0.32 | 1 | 0.1 ± 0.32 |
| 68 | *Ovis cycloceros (O. v. cycloceros)* | Afghan Urial | 1 | 0.1 ± 0.32 | 1 | 0.1 ± 0.32 |
| 69 | *Papio hamadryas* | Hamadryas Baboon | 1 | 0.1 ± 0.32 | 1 | 0.1 ± 0.32 |
| 70 | *Python sebae* | Central African Rock Python | 1 | 0.1 ± 0.32 | 1 | 0.1 ± 0.32 |
| 71 | *Tauraco porphyreolophus* | Purple-crested Turaco | 1 | 0.1 ± 0.32 | 1 | 0.1 ± 0.32 |
| 72 | *Theropithecus gelada* | Gelada | 1 | 0.1 ± 0.32 | 1 | 0.1 ± 0.32 |
| 73 | *Varanus niloticus* | Nile Monitor | 1 | 0.1 ± 0.32 | 1 | 0.1 ± 0.32 |
| **Totals** | | | **9733** | | **8060.07** | |

**Table S15. CITES-listed species imported to Italy as hunting trophies (2015-2024; *n* = 56), estimated number of trophies, mean (± SD) number of trophies imported/year, WOEs imported, and mean (± SD) WOEs imported/year, ranked by trophies.**

| Rank | Scientific name | Common name | Trophies | Mean + SD trophies/year | WOEs | Mean + SD WOEs/year |
|---|---|---|---|---|---|---|
| 1 | ***Crocodylus niloticus*** | **Nile Crocodile** | 1202 | 120.2 ± 210.61 | 685 | 68.5 ± 117.48 |
| 2 | ***Hippopotamus amphibius*** | **Common Hippopotamus** | 279 | 27.9 ± 32.86 | 50.45 | 5.05 ± 3.65 |
| 3 | ***Loxodonta africana*** | **African Savanna Elephant** | 185 | 18.5 ± 13.11 | 98.07 | 9.81 ± 8.15 |
| 4 | ***Equus zebra hartmannae*** | **Hartmann's Mountain Zebra** | 146 | 14.6 ± 6.36 | 146 | 14.6 ± 6.36 |
| 5 | ***Ursus americanus*** | **American Black Bear** | 84 | 8.4 ± 10.92 | 57 | 5.7 ± 6.6 |
| 6 | ***Papio ursinus*** | **Chacma Baboon** | 55 | 5.5 ± 3.54 | 51 | 5.1 ± 2.56 |
| 7 | ***Panthera pardus*** | **Leopard** | 53 | 5.3 ± 4.35 | 53 | 5.3 ± 4.35 |
| 8 | *Capra sibirica* | Siberian Ibex | 40 | 4 ± 3.94 | 40 | 4 ± 3.94 |
| 9 | *Capra hircus aegagrus (Capra aegagrus)* | Wild Goat | 28 | 2.8 ± 3.33 | 28 | 2.8 ± 3.33 |
| 10 | ***Giraffa camelopardalis*** | **Giraffe** | 27 | 2.7 ± 2.95 | 24 | 2.4 ± 2.72 |
| 11 | ***Capra caucasica*** | **Western Tur** | 24 | 2.4 ± 3.89 | 24 | 2.4 ± 3.89 |
| 12 | *Antilope cervicapra* | Blackbuck | 21 | 2.1 ± 4.25 | 21 | 2.1 ± 4.25 |
| 13 | ***Panthera leo*** | **Lion** | 21 | 2.1 ± 2.51 | 21 | 2.1 ± 2.51 |
| 14 | ***Puma concolor*** | **Puma** | 21 | 2.1 ± 2.13 | 21 | 2.1 ± 2.13 |
| 15 | ***Kobus leche*** | **Lechwe** | 20 | 2 ± 1.76 | 20 | 2 ± 1.76 |
| 16 | ***Ovis ammon*** | **Wild Sheep** | 17 | 1.7 ± 2 | 17 | 1.7 ± 2 |
| 17 | ***Caracal caracal*** | **Caracal** | 16 | 1.6 ± 2.07 | 16 | 1.6 ± 2.07 |
| 18 | ***Ceratotherium simum simum*** | **Southern White Rhinoceros** | 14 | 1.4 ± 2.5 | 14 | 1.4 ± 2.5 |
| 19 | ***Papio cynocephalus*** | **Yellow Baboon** | 14 | 1.4 ± 1.58 | 14 | 1.4 ± 1.58 |
| 20 | *Civettictis civetta* | African Civet | 13 | 1.3 ± 1.49 | 13 | 1.3 ± 1.49 |
| 21 | ***Damaliscus pygargus pygargus*** | **Bontebok** | 12 | 1.2 ± 1.14 | 12 | 1.2 ± 1.14 |
| 22 | ***Ursus arctos*** | **Brown Bear** | 10 | 1 ± 1.15 | 10 | 1 ± 1.15 |
| 23 | ***Ursus maritimus*** | **Polar Bear** | 9 | 0.9 ± 1.52 | 7 | 0.7 ± 1.16 |
| 24 | ***Ovis polii*** | **Marco Polo Argali** | 8 | 0.8 ± 1.14 | 8 | 0.8 ± 1.14 |
| 25 | ***Papio anubis*** | **Olive Baboon** | 7 | 0.7 ± 1.57 | 7 | 0.7 ± 1.57 |
| 26 | ***Acinonyx jubatus*** | **Cheetah** | 6 | 0.6 ± 0.7 | 6 | 0.6 ± 0.7 |
| 27 | ***Puma concolor couguar*** | **Florida Puma** | 6 | 0.6 ± 1.35 | 6 | 0.6 ± 1.35 |

| | | | | | | |
|---|---|---|---|---|---|---|
| 28 | ***Canis lupus*** | **Grey Wolf** | 5 | 0.5 ± 0.85 | 3 | 0.3 ± 0.67 |
| 29 | ***Chlorocebus pygerythrus*** | **Vervet Monkey** | 5 | 0.5 ± 0.53 | 5 | 0.5 ± 0.53 |
| 30 | ***Felis silvestris*** | **European Wildcat** | 5 | 0.5 ± 0.85 | 5 | 0.5 ± 0.85 |
| 31 | *Odobenus rosmarus* | Walrus | 5 | 0.5 ± 1.58 | 2 | 0.2 ± 0.63 |
| 32 | ***Philantomba monticola*** | **Blue Duiker** | 5 | 0.5 ± 0.85 | 5 | 0.5 ± 0.85 |
| 33 | ***Ammotragus lervia*** | **Aoudad** | 4 | 0.4 ± 0.7 | 4 | 0.4 ± 0.7 |
| 34 | *Damaliscus pygargus* | Blesbok | 4 | 0.4 ± 0.97 | 4 | 0.4 ± 0.97 |
| 35 | ***Lynx canadensis*** | **Canada Lynx** | 4 | 0.4 ± 0.84 | 4 | 0.4 ± 0.84 |
| 36 | ***Lynx rufus*** | **Bobcat** | 4 | 0.4 ± 0.84 | 4 | 0.4 ± 0.84 |
| 37 | ***Ovis cycloceros (O. v. cycloceros)*** | **Afghan Urial** | 4 | 0.4 ± 0.84 | 4 | 0.4 ± 0.84 |
| 38 | ***Alligator mississippiensis*** | **American Alligator** | 2 | 0.2 ± 0.42 | 2 | 0.2 ± 0.42 |
| 39 | ***Cephalophus silvicultor*** | **Yellow-backed Duiker** | 2 | 0.2 ± 0.63 | 2 | 0.2 ± 0.63 |
| 40 | ***Leptailurus serval*** | **Serval** | 2 | 0.2 ± 0.42 | 2 | 0.2 ± 0.42 |
| 41 | ***Lynx lynx*** | **Eurasian lynx** | 2 | 0.2 ± 0.63 | 2 | 0.2 ± 0.63 |
| 42 | ***Ovis darwini (Ovis ammon darwini)*** | **Gobi Argali** | 2 | 0.2 ± 0.63 | 2 | 0.2 ± 0.63 |
| 43 | ***Ovis punjabiensis (Ovis v. punjabensis)*** | **Punjab Urial** | 2 | 0.2 ± 0.63 | 2 | 0.2 ± 0.63 |
| 44 | ***Capra falconeri*** | **Markhor** | 1 | 0.1 ± 0.32 | 1 | 0.1 ± 0.32 |
| 45 | ***Capra falconeri heptneri*** | **Markhor** | 1 | 0.1 ± 0.32 | 1 | 0.1 ± 0.32 |
| 46 | ***Ceratotherium simum*** | **White Rhinoceros** | 1 | 0.1 ± 0.32 | 1 | 0.1 ± 0.32 |
| 47 | ***Colobus guereza*** | **Guereza** | 1 | 0.1 ± 0.32 | 1 | 0.1 ± 0.32 |
| 48 | ***Diceros bicornis*** | **Black Rhinoceros** | 1 | 0.1 ± 0.32 | 1 | 0.1 ± 0.32 |
| 49 | ***Oryx dammah*** | **Scimitar-horned Oryx** | 1 | 0.1 ± 0.32 | 1 | 0.1 ± 0.32 |
| 50 | ***Oryx leucoryx*** | **Arabian Oryx** | 1 | 0.1 ± 0.32 | 1 | 0.1 ± 0.32 |
| 51 | ***Ovis canadensis*** | **Bighorn Sheep** | 1 | 0.1 ± 0.32 | 1 | 0.1 ± 0.32 |
| 52 | ***Ovis cycloceros cycloceros (O.v. cycloceros)*** | **Afghan Urial (subsp. cycloceros)** | 1 | 0.1 ± 0.32 | 1 | 0.1 ± 0.32 |
| 53 | ***Papio hamadryas*** | **Hamadryas Baboon** | 1 | 0.1 ± 0.32 | 1 | 0.1 ± 0.32 |
| 54 | *Proteles cristata* | Aardwolf | 1 | 0.1 ± 0.32 | 1 | 0.1 ± 0.32 |
| 55 | *Pseudois nayaur* | Blue Sheep | 1 | 0.1 ± 0.32 | 1 | 0.1 ± 0.32 |
| 56 | *Python sebae* | Central African Rock Python | 1 | 0.1 ± 0.32 | 1 | 0.1 ± 0.32 |
| **Totals** | | | **2408** | | **1534.53** | |

**Table S16. CITES-listed species imported to the Netherlands as hunting trophies (2015-2024; *n* = 38), estimated number of trophies, mean (± SD) number of trophies imported/year, WOEs imported, and mean (± SD) WOEs imported/year, ranked by trophies. Asterix indicates populations of species split between Annex A and B of the EU WTRs.**

| Rank | Scientific name | Common name | Trophies | Mean + SD trophies/year | WOEs | Mean + SD WOEs/year |
|---|---|---|---|---|---|---|
| 1 | *Ursus americanus* | American Black Bear | 156 | 15.6 ± 12.1 | 110 | 11 ± 9.18 |
| 2 | *Equus zebra hartmannae* | Hartmann's Mountain Zebra | 66 | 6.6 ± 3.57 | 65.25 | 6.53 ± 3.37 |
| 3 | *Papio ursinus* | Chacma Baboon | 44 | 4.4 ± 1.9 | 44 | 4.4 ± 1.9 |
| 4 | *Capra sibirica* | Siberian Ibex | 26 | 2.6 ± 1.84 | 26 | 2.6 ± 1.84 |
| 5 | *Damaliscus pygargus pygargus* | Bontebok | 23 | 2.3 ± 3.43 | 23 | 2.3 ± 3.43 |
| 6 | *Giraffa camelopardalis* | Giraffe | 16 | 1.6 ± 2.22 | 16 | 1.6 ± 2.22 |
| 7 | ***Caracal caracal**** | **Caracal** | 14 | 1.4 ± 1.58 | 13 | 1.3 ± 1.64 |
| 8 | *Leptailurus serval* | Serval | 12 | 1.2 ± 1.23 | 12 | 1.2 ± 1.23 |
| 9 | ***Canis lupus**** | **Grey Wolf** | 10 | 1 ± 1.41 | 9 | 0.9 ± 1.37 |
| 10 | *Lynx canadensis* | Canada Lynx | 10 | 1 ± 2 | 10 | 1 ± 2 |
| 11 | *Lynx rufus* | Bobcat | 10 | 1 ± 1.94 | 10 | 1 ± 1.94 |
| 12 | ***Panthera pardus*** | **Leopard** | 10 | 1 ± 1.76 | 10 | 1 ± 1.76 |
| 13 | *Alligator mississippiensis* | American Alligator | 9 | 0.9 ± 2.18 | 3 | 0.3 ± 0.48 |
| 14 | *Capra caucasica* | Western Tur | 7 | 0.7 ± 1.34 | 7 | 0.7 ± 1.34 |
| 15 | *Antilope cervicapra* | Blackbuck | 6 | 0.6 ± 1.35 | 6 | 0.6 ± 1.35 |
| 16 | *Capra hircus aegagrus (Capra aegagrus)* | Wild Goat | 5 | 0.5 ± 0.71 | 5 | 0.5 ± 0.71 |
| 17 | *Civettictis civetta* | African Civet | 5 | 0.5 ± 0.85 | 5 | 0.5 ± 0.85 |
| 18 | ***Crocodylus niloticus**** | **Nile Crocodile** | 5 | 0.5 ± 1.27 | 5 | 0.5 ± 1.27 |
| 19 | ***Hippopotamus amphibius*** | **Common Hippopotamus** | 5 | 0.5 ± 0.97 | 5 | 0.5 ± 0.97 |
| 20 | ***Loxodonta africana**** | **African Savanna Elephant** | 5 | 0.5 ± 0.97 | 3 | 0.3 ± 0.48 |
| 21 | ***Ursus arctos*** | **Brown Bear** | 5 | 0.5 ± 0.85 | 4 | 0.4 ± 0.7 |
| 22 | *Felis lybica* | Afro-Asiatic Wildcat | 4 | 0.4 ± 0.84 | 4 | 0.4 ± 0.84 |
| 23 | *Kobus leche* | Lechwe | 4 | 0.4 ± 0.7 | 4 | 0.4 ± 0.7 |
| 24 | ***Puma concolor**** | **Puma** | 4 | 0.4 ± 0.84 | 4 | 0.4 ± 0.84 |
| 25 | *Damaliscus pygargus* | Blesbok | 3 | 0.3 ± 0.67 | 3 | 0.3 ± 0.67 |
| 26 | *Papio cynocephalus* | Yellow Baboon | 3 | 0.3 ± 0.67 | 3 | 0.3 ± 0.67 |

| | | | | | | |
|---|---|---|---|---|---|---|
| 27 | *Hyaena hyaena* | Striped Hyena | 2 | 0.2 ± 0.63 | 2 | 0.2 ± 0.63 |
| 28 | *Puma concolor couguar* | Florida Puma | 2 | 0.2 ± 0.63 | 2 | 0.2 ± 0.63 |
| 29 | ***Ursus maritimus*** | **Polar Bear** | 2 | 0.2 ± 0.63 | 2 | 0.2 ± 0.63 |
| 30 | *Ammotragus lervia* | Aoudad | 1 | 0.1 ± 0.32 | 1 | 0.1 ± 0.32 |
| 31 | *Canis aureus* | Common Jackal | 1 | 0.1 ± 0.32 | 1 | 0.1 ± 0.32 |
| 32 | ***Felis silvestris*** | **European Wildcat** | 1 | 0.1 ± 0.32 | 1 | 0.1 ± 0.32 |
| 33 | ***Lynx lynx*** | **Eurasian lynx** | 1 | 0.1 ± 0.32 | 1 | 0.1 ± 0.32 |
| 34 | *Mellivora capensis* | Honey Badger | 1 | 0.1 ± 0.32 | 1 | 0.1 ± 0.32 |
| 35 | *Otolemur crassicaudatus* | Thick-tailed Greater Galago | 1 | 0.1 ± 0.32 | 1 | 0.1 ± 0.32 |
| 36 | ***Ovis darwini (Ovis ammon darwini)*** | **Gobi Argali** | 1 | 0.1 ± 0.32 | 1 | 0.1 ± 0.32 |
| 37 | ***Panthera leo**** | **Lion** | 1 | 0.1 ± 0.32 | 1 | 0.1 ± 0.32 |
| 38 | *Pseudois nayaur* | Blue Sheep | 1 | 0.1 ± 0.32 | 1 | 0.1 ± 0.32 |
| **Totals** | | | **482** | | **424.25** | |

**Table S17. CITES-listed species imported to Poland as hunting trophies (2015-2024; *n* = 57), estimated number of trophies, mean (± SD) number of trophies imported/year, WOEs imported, and mean (± SD) WOEs imported/year, ranked by trophies. Asterix indicates populations of species split between Annex A and B of the EU WTRs.**

| Rank | Scientific name | Common name | Trophies | Mean + SD trophies/year | WOEs | Mean + SD WOEs/year |
|---|---|---|---|---|---|---|
| 1 | *Ursus americanus* | American Black Bear | 343 | 34.3 ± 24.73 | 316 | 31.6 ± 24.01 |
| 2 | ***Ursus arctos*** | **Brown Bear** | 139 | 13.9 ± 20.89 | 139 | 13.9 ± 20.89 |
| 3 | *Papio ursinus* | Chacma Baboon | 120 | 12 ± 6.9 | 119 | 11.9 ± 6.94 |
| 4 | *Giraffa camelopardalis* | Giraffe | 98 | 9.8 ± 12.99 | 87 | 8.7 ± 11.43 |
| 5 | *Equus zebra hartmannae* | Hartmann's Mountain Zebra | 82 | 8.2 ± 2.94 | 80 | 8 ± 2.79 |
| 6 | ***Crocodylus niloticus**** | **Nile Crocodile** | 72 | 7.2 ± 4.34 | 72 | 7.2 ± 4.34 |
| 7 | ***Panthera pardus*** | **Leopard** | 56 | 5.6 ± 3.98 | 56 | 5.6 ± 3.98 |
| 8 | *Chlorocebus pygerythrus* | Vervet Monkey | 55 | 5.5 ± 5.6 | 55 | 5.5 ± 5.6 |
| 9 | ***Loxodonta africana**** | **African Savanna Elephant** | 52 | 5.2 ± 4.69 | 35.19 | 3.52 ± 2.47 |
| 10 | ***Caracal caracal**** | **Caracal** | 43 | 4.3 ± 3.86 | 43 | 4.3 ± 3.86 |
| 11 | *Capra sibirica* | Siberian Ibex | 41 | 4.1 ± 3.28 | 41 | 4.1 ± 3.28 |
| 12 | ***Puma concolor**** | **Puma** | 37 | 3.7 ± 3.65 | 36 | 3.6 ± 3.75 |
| 13 | ***Acinonyx jubatus*** | **Cheetah** | 36 | 3.6 ± 3.24 | 36 | 3.6 ± 3.24 |
| 14 | ***Hippopotamus amphibius*** | **Common Hippopotamus** | 35 | 3.5 ± 4.6 | 23.99 | 2.4 ± 1.65 |
| 15 | *Leptailurus serval* | Serval | 32 | 3.2 ± 4.05 | 32 | 3.2 ± 4.05 |
| 16 | *Lynx canadensis* | Canada Lynx | 30 | 3 ± 3.4 | 30 | 3 ± 3.4 |
| 17 | ***Canis lupus**** | **Grey Wolf** | 24 | 2.4 ± 3.5 | 21 | 2.1 ± 3.57 |
| 18 | ***Ceratotherium simum simum**** | **Southern White Rhinoceros** | 22 | 2.2 ± 2.04 | 22 | 2.2 ± 2.04 |
| 19 | *Civettictis civetta* | African Civet | 16 | 1.6 ± 1.43 | 16 | 1.6 ± 1.43 |
| 20 | *Kobus leche* | Lechwe | 16 | 1.6 ± 1.58 | 13 | 1.3 ± 1.34 |
| 21 | *Mellivora capensis* | Honey Badger | 14 | 1.4 ± 1.17 | 14 | 1.4 ± 1.17 |
| 22 | *Damaliscus pygargus pygargus* | Bontebok | 12 | 1.2 ± 1.23 | 12 | 1.2 ± 1.23 |
| 23 | ***Panthera leo**** | **Lion** | 11 | 1.1 ± 1.66 | 11 | 1.1 ± 1.66 |
| 24 | *Philantomba monticola* | Blue Duiker | 11 | 1.1 ± 1.2 | 11 | 1.1 ± 1.2 |
| 25 | *Puma concolor couguar* | Florida Puma | 11 | 1.1 ± 2.33 | 11 | 1.1 ± 2.33 |

| | | | | | | |
|---|---|---|---|---|---|---|
| 26 | *Felis lybica* | Afro-Asiatic Wildcat | 10 | 1 ± 1.15 | 10 | 1 ± 1.15 |
| 27 | *Antilope cervicapra* | Blackbuck | 7 | 0.7 ± 1.25 | 7 | 0.7 ± 1.25 |
| 28 | *Odobenus rosmarus* | Walrus | 7 | 0.7 ± 2.21 | 1 | 0.1 ± 0.32 |
| 29 | *Otolemur crassicaudatus* | Thick-tailed Greater Galago | 7 | 0.7 ± 1.34 | 7 | 0.7 ± 1.34 |
| 30 | *Proteles cristata* | Aardwolf | 6 | 0.6 ± 0.52 | 6 | 0.6 ± 0.52 |
| 31 | *Capra hircus aegagrus (Capra aegagrus)* | Wild Goat | 5 | 0.5 ± 0.71 | 5 | 0.5 ± 0.71 |
| 32 | ***Ceratotherium simum**** | **White Rhinoceros** | 4 | 0.4 ± 0.7 | 4 | 0.4 ± 0.7 |
| 33 | *Lynx rufus* | Bobcat | 4 | 0.4 ± 0.97 | 4 | 0.4 ± 0.97 |
| 34 | *Meleagris ocellata* | Ocellated Turkey | 4 | 0.4 ± 1.26 | 4 | 0.4 ± 1.26 |
| 35 | ***Ovis ammon*** | **Wild Sheep** | 4 | 0.4 ± 0.7 | 4 | 0.4 ± 0.7 |
| 36 | ***Ovis polii*** | **Marco Polo Argali** | 4 | 0.4 ± 0.52 | 4 | 0.4 ± 0.52 |
| 37 | *Papio cynocephalus* | Yellow Baboon | 4 | 0.4 ± 0.7 | 4 | 0.4 ± 0.7 |
| 38 | *Tauraco porphyreolophus* | Purple-crested Turaco | 4 | 0.4 ± 0.84 | 4 | 0.4 ± 0.84 |
| 39 | *Ammotragus lervia* | Aoudad | 3 | 0.3 ± 0.67 | 3 | 0.3 ± 0.67 |
| 40 | *Capra caucasica* | Western Tur | 3 | 0.3 ± 0.67 | 3 | 0.3 ± 0.67 |
| 41 | *Cephalophus dorsalis* | Bay Duiker | 3 | 0.3 ± 0.48 | 3 | 0.3 ± 0.48 |
| 42 | ***Felis silvestris*** | **European Wildcat** | 3 | 0.3 ± 0.48 | 3 | 0.3 ± 0.48 |
| 43 | *Papio anubis* | Olive Baboon | 3 | 0.3 ± 0.48 | 3 | 0.3 ± 0.48 |
| 44 | *Crax rubra* | Great Curassow | 2 | 0.2 ± 0.63 | 2 | 0.2 ± 0.63 |
| 45 | *Galago senegalensis* | Northern Lesser Galago | 2 | 0.2 ± 0.63 | 2 | 0.2 ± 0.63 |
| 46 | ***Loxodonta cyclotis*** | **African Forest Elephant** | 2 | 0.2 ± 0.63 | 2 | 0.2 ± 0.63 |
| 47 | ***Lynx lynx*** | **Eurasian lynx** | 2 | 0.2 ± 0.63 | 2 | 0.2 ± 0.63 |
| 48 | ***Oryx dammah*** | **Scimitar-horned Oryx** | 2 | 0.2 ± 0.42 | 2 | 0.2 ± 0.42 |
| 49 | *Ovis cycloceros (O. v. cycloceros)* | Afghan Urial | 2 | 0.2 ± 0.63 | 2 | 0.2 ± 0.63 |
| 50 | ***Ursus maritimus*** | **Polar Bear** | 2 | 0.2 ± 0.63 | 2 | 0.2 ± 0.63 |
| 51 | *Alligator mississippiensis* | American Alligator | 1 | 0.1 ± 0.32 | 1 | 0.1 ± 0.32 |
| 52 | *Damaliscus pygargus* | Blesbok | 1 | 0.1 ± 0.32 | 1 | 0.1 ± 0.32 |
| 53 | *Equus zebra zebra* | Cape Mountain Zebra | 1 | 0.1 ± 0.32 | 1 | 0.1 ± 0.32 |
| 54 | *Lissotis melanogaster* | Black-bellied Bustard | 1 | 0.1 ± 0.32 | 1 | 0.1 ± 0.32 |
| 55 | *Lophaetus occipitalis* | Long-crested Eagle | 1 | 0.1 ± 0.32 | 1 | 0.1 ± 0.32 |
| 56 | *Polemaetus bellicosus* | Martial Eagle | 1 | 0.1 ± 0.32 | 1 | 0.1 ± 0.32 |

| | | | | | | |
|---|---|---|---|---|---|---|
| 57 | *Sarkidiornis melanotos* | Knob-billed Duck | 1 | 0.1 ± 0.32 | 1 | 0.1 ± 0.32 |
| **Totals** | | | **1514** | | **1432.19** | |

**Table S18. CITES-listed species imported to Spain as hunting trophies (2015-2024; *n* = 65), estimated number of trophies, mean (± SD) number of trophies imported/year, WOEs imported, and mean (± SD) WOEs imported/year, ranked by trophies. Asterix indicates populations of species split between Annex A and B of the EU WTRs.**

| Rank | Scientific name | Common name | Trophies | Mean + SD trophies/year | WOEs | Mean + SD WOEs/year |
|---|---|---|---|---|---|---|
| 1 | ***Hippopotamus amphibius*** | **Common Hippopotamus** | 851 | 85.1 ± 47.35 | 226.607 | 22.66 ± 9.38 |
| 2 | ***Loxodonta africana**** | **African Savanna Elephant** | 559 | 55.9 ± 46.07 | 301.714 | 30.17 ± 26.14 |
| 3 | *Equus zebra hartmannae* | Hartmann's Mountain Zebra | 436 | 43.6 ± 11.88 | 433 | 43.3 ± 11.94 |
| 4 | *Papio ursinus* | Chacma Baboon | 390 | 39 ± 12 | 390 | 39 ± 12 |
| 5 | *Capra sibirica* | Siberian Ibex | 359 | 35.9 ± 21.42 | 359 | 35.9 ± 21.42 |
| 6 | *Ursus americanus* | American Black Bear | 352 | 35.2 ± 30.54 | 267.5 | 26.75 ± 20.96 |
| 7 | *Antilope cervicapra* | Blackbuck | 208 | 20.8 ± 18.25 | 208 | 20.8 ± 18.25 |
| 8 | ***Giraffa camelopardalis*** | **Giraffe** | 196 | 19.6 ± 20.33 | 187 | 18.7 ± 19.86 |
| 9 | ***Caracal caracal**** | **Caracal** | 191 | 19.1 ± 9.9 | 191 | 19.1 ± 9.9 |
| 10 | ***Crocodylus niloticus**** | **Nile Crocodile** | 167 | 16.7 ± 8.41 | 167 | 16.7 ± 8.41 |
| 11 | ***Panthera pardus*** | **Leopard** | 151 | 15.1 ± 5.57 | 151 | 15.1 ± 5.57 |
| 12 | ***Ursus arctos*** | **Brown Bear** | 131 | 13.1 ± 20.71 | 123 | 12.3 ± 20.86 |
| 13 | *Capra hircus aegagrus (Capra aegagrus)* | Wild Goat | 117 | 11.7 ± 5.7 | 117 | 11.7 ± 5.7 |
| 14 | *Chlorocebus pygerythrus* | Vervet Monkey | 112 | 11.2 ± 7.98 | 112 | 11.2 ± 7.98 |
| 15 | *Civettictis civetta* | African Civet | 89 | 8.9 ± 4.82 | 89 | 8.9 ± 4.82 |
| 16 | *Damaliscus pygargus pygargus* | Bontebok | 88 | 8.8 ± 7.04 | 88 | 8.8 ± 7.04 |
| 17 | *Capra caucasica* | Western Tur | 81 | 8.1 ± 7.69 | 81 | 8.1 ± 7.69 |
| 18 | ***Ovis ammon*** | **Wild Sheep** | 68 | 6.8 ± 6.2 | 68 | 6.8 ± 6.2 |
| 19 | ***Acinonyx jubatus*** | **Cheetah** | 63 | 6.3 ± 4.27 | 63 | 6.3 ± 4.27 |
| 20 | ***Ovis polii*** | **Marco Polo Argali** | 50 | 5 ± 6.2 | 50 | 5 ± 6.2 |
| 21 | ***Panthera leo**** | **Lion** | 50 | 5 ± 1.76 | 50 | 5 ± 1.76 |
| 22 | *Philantomba monticola* | Blue Duiker | 50 | 5 ± 2.98 | 50 | 5 ± 2.98 |
| 23 | *Papio cynocephalus* | Yellow Baboon | 44 | 4.4 ± 2.01 | 44 | 4.4 ± 2.01 |
| 24 | ***Ceratotherium simum simum**** | **Southern White Rhinoceros** | 43 | 4.3 ± 2.87 | 43 | 4.3 ± 2.87 |
| 25 | *Leptailurus serval* | Serval | 43 | 4.3 ± 1.64 | 43 | 4.3 ± 1.64 |
| 26 | *Mellivora capensis* | Honey Badger | 40 | 4 ± 2.36 | 40 | 4 ± 2.36 |

| | | | | | | |
|---|---|---|---|---|---|---|
| 27 | *Kobus leche* | Lechwe | 38 | 3.8 ± 2.74 | 38 | 3.8 ± 2.74 |
| 28 | ***Ursus maritimus*** | **Polar Bear** | 37 | 3.7 ± 2.31 | 33 | 3.3 ± 1.83 |
| 29 | ***Canis lupus**** | **Grey Wolf** | 36 | 3.6 ± 3.03 | 35 | 3.5 ± 2.84 |
| 30 | *Felis lybica* | Afro-Asiatic Wildcat | 28 | 2.8 ± 2.15 | 28 | 2.8 ± 2.15 |
| 31 | *Odobenus rosmarus* | Walrus | 23 | 2.3 ± 3.95 | 8 | 0.8 ± 1.4 |
| 32 | *Equus zebra zebra* | Cape Mountain Zebra | 16 | 1.6 ± 3.34 | 16 | 1.6 ± 3.34 |
| 33 | ***Felis silvestris*** | **European Wildcat** | 16 | 1.6 ± 1.51 | 16 | 1.6 ± 1.51 |
| 34 | ***Puma concolor**** | **Puma** | 13 | 1.3 ± 2.06 | 13 | 1.3 ± 2.06 |
| 35 | *Ammotragus lervia* | Aoudad | 11 | 1.1 ± 0.99 | 11 | 1.1 ± 0.99 |
| 36 | *Pseudois nayaur* | Blue Sheep | 11 | 1.1 ± 1.29 | 11 | 1.1 ± 1.29 |
| 37 | *Ovis punjabiensis (Ovis v. punjabensis)* | Punjab Urial | 10 | 1 ± 1.25 | 10 | 1 ± 1.25 |
| 38 | *Papio anubis* | Olive Baboon | 10 | 1 ± 0.82 | 10 | 1 ± 0.82 |
| 39 | *Lynx rufus* | Bobcat | 9 | 0.9 ± 1.2 | 9 | 0.9 ± 1.2 |
| 40 | ***Oryx dammah*** | **Scimitar-horned Oryx** | 9 | 0.9 ± 1.1 | 9 | 0.9 ± 1.1 |
| 41 | *Damaliscus pygargus* | Blesbok | 8 | 0.8 ± 1.62 | 8 | 0.8 ± 1.62 |
| 42 | ***Ceratotherium simum**** | **White Rhinoceros** | 7 | 0.7 ± 1.06 | 7 | 0.7 ± 1.06 |
| 43 | *Ovis aries* | Urial | 7 | 0.7 ± 1.64 | 7 | 0.7 ± 1.64 |
| 44 | *Canis aureus* | Common Jackal | 6 | 0.6 ± 1.26 | 6 | 0.6 ± 1.26 |
| 45 | ***Capra falconeri*** | **Marhor** | 6 | 0.6 ± 1.07 | 6 | 0.6 ± 1.07 |
| 46 | ***Capra falconeri heptneri*** | **Markhor** | 6 | 0.6 ± 0.84 | 6 | 0.6 ± 0.84 |
| 47 | *Cephalophus silvicultor* | Yellow-backed Duiker | 6 | 0.6 ± 0.84 | 6 | 0.6 ± 0.84 |
| 48 | *Lynx canadensis* | Canada Lynx | 6 | 0.6 ± 1.35 | 6 | 0.6 ± 1.35 |
| 49 | *Ovis cycloceros (O. v. cycloceros)* | Afghan Urial | 6 | 0.6 ± 1.35 | 6 | 0.6 ± 1.35 |
| 50 | *Alligator mississippiensis* | American Alligator | 5 | 0.5 ± 0.71 | 5 | 0.5 ± 0.71 |
| 51 | *Cephalophus dorsalis* | Bay Duiker | 5 | 0.5 ± 0.85 | 5 | 0.5 ± 0.85 |
| 52 | *Ovis cycloceros cycloceros (O.v. cycloceros)* | Afghan Urial (subsp. cycloceros) | 4 | 0.4 ± 0.84 | 4 | 0.4 ± 0.84 |
| 53 | ***Ovis darwini (Ovis ammon darwini)*** | **Gobi Argali** | 4 | 0.4 ± 0.7 | 4 | 0.4 ± 0.7 |
| 54 | *Proteles cristata* | Aardwolf | 4 | 0.4 ± 0.97 | 4 | 0.4 ± 0.97 |
| 55 | *Python sebae* | Central African Rock Python | 4 | 0.4 ± 0.7 | 4 | 0.4 ± 0.7 |
| 56 | *Ovis bochariensis (Ovis v. bochariensis)* | Bukhara Urial | 3 | 0.3 ± 0.95 | 3 | 0.3 ± 0.95 |
| 57 | ***Ovis severtzovi (O.a. severtzovi)*** | **Severtzov's Argali** | 3 | 0.3 ± 0.67 | 3 | 0.3 ± 0.67 |

| | | | | | | |
|---|---|---|---|---|---|---|
| 58 | ***Lynx lynx*** | **Eurasian lynx** | 2 | 0.2 ± 0.42 | 2 | 0.2 ± 0.42 |
| 59 | *Ovis cycloceros arkal (O.v. arkal)* | Afghan Urial (subsp. arkal) | 2 | 0.2 ± 0.63 | 2 | 0.2 ± 0.63 |
| 60 | *Varanus albigularis* | White-throated Monitor | 2 | 0.2 ± 0.63 | 2 | 0.2 ± 0.63 |
| 61 | *Cephalophus ogilbyi* | Ogilby's Duiker | 1 | 0.1 ± 0.32 | 1 | 0.1 ± 0.32 |
| 62 | *Colobus guereza* | Guereza | 1 | 0.1 ± 0.32 | 1 | 0.1 ± 0.32 |
| 63 | ***Diceros bicornis*** | **Black Rhinoceros** | 1 | 0.1 ± 0.32 | 1 | 0.1 ± 0.32 |
| 64 | *Ovis canadensis* | Bighorn Sheep | 1 | 0.1 ± 0.32 | 1 | 0.1 ± 0.32 |
| 65 | *Philantomba maxwellii* | Maxwell's Duiker | 1 | 0.1 ± 0.32 | 1 | 0.1 ± 0.32 |
| **Totals** | | | **5297** | | **4290.82** | |

**Table S19. CITES-listed species imported to the United Kingdom as hunting trophies (2015-2024; *n* = 46), estimated number of trophies, mean (± SD) number of trophies imported/year, WOEs imported, and mean (± SD) WOEs imported/year, ranked by trophies.**

| Rank | Scientific name | Common name | Trophies | Mean + SD trophies/year | WOEs | Mean + SD WOEs/year |
|---|---|---|---|---|---|---|
| 1 | ***Ursus americanus*** | **American Black Bear** | 200 | 20 ± 15.57 | 86 | 8.6 ± 6.02 |
| 2 | ***Papio ursinus*** | **Chacma Baboon** | 129 | 12.9 ± 9.75 | 129 | 12.9 ± 9.75 |
| 3 | ***Giraffa camelopardalis*** | **Giraffe** | 113 | 11.3 ± 15.14 | 68 | 6.8 ± 7.9 |
| 4 | ***Equus zebra hartmannae*** | **Hartmann's Mountain Zebra** | 101 | 10.1 ± 6.47 | 94 | 9.4 ± 6.13 |
| 5 | ***Loxodonta africana*** | **African Savanna Elephant** | 97 | 9.7 ± 11.58 | 38.288 | 3.83 ± 3.02 |
| 6 | ***Hippopotamus amphibius*** | **Common Hippopotamus** | 59 | 5.9 ± 4.98 | 22.577 | 2.26 ± 1.24 |
| 7 | ***Chlorocebus pygerythrus*** | **Vervet Monkey** | 52 | 5.2 ± 3.26 | 52 | 5.2 ± 3.26 |
| 8 | ***Crocodylus niloticus*** | **Nile Crocodile** | 52 | 5.2 ± 4.49 | 51 | 5.1 ± 4.2 |
| 9 | ***Caracal caracal*** | **Caracal** | 49 | 4.9 ± 3.96 | 49 | 4.9 ± 3.96 |
| 10 | *Capra sibirica* | Siberian Ibex | 34 | 3.4 ± 4.12 | 34 | 3.4 ± 4.12 |
| 11 | ***Panthera pardus*** | **Leopard** | 27 | 2.7 ± 1.49 | 27 | 2.7 ± 1.49 |
| 12 | ***Ursus arctos*** | **Brown Bear** | 27 | 2.7 ± 2.41 | 27 | 2.7 ± 2.41 |
| 13 | ***Leptailurus serval*** | **Serval** | 18 | 1.8 ± 1.69 | 18 | 1.8 ± 1.69 |
| 14 | ***Damaliscus pygargus pygargus*** | **Bontebok** | 15 | 1.5 ± 1.72 | 15 | 1.5 ± 1.72 |
| 15 | *Antilope cervicapra* | Blackbuck | 14 | 1.4 ± 1.96 | 14 | 1.4 ± 1.96 |
| 16 | ***Kobus leche*** | **Lechwe** | 14 | 1.4 ± 1.65 | 14 | 1.4 ± 1.65 |
| 17 | ***Canis lupus*** | **Grey Wolf** | 13 | 1.3 ± 1.7 | 11 | 1.1 ± 1.6 |
| 18 | *Capra hircus aegagrus (Capra aegagrus)* | Wild Goat | 8 | 0.8 ± 0.92 | 8 | 0.8 ± 0.92 |
| 19 | ***Puma concolor*** | **Puma** | 8 | 0.8 ± 1.69 | 8 | 0.8 ± 1.69 |
| 20 | ***Papio cynocephalus*** | **Yellow Baboon** | 7 | 0.7 ± 0.82 | 7 | 0.7 ± 0.82 |
| 21 | ***Alligator mississippiensis*** | **American Alligator** | 6 | 0.6 ± 0.84 | 6 | 0.6 ± 0.84 |
| 22 | *Civettictis civetta* | African Civet | 6 | 0.6 ± 0.52 | 6 | 0.6 ± 0.52 |
| 23 | ***Papio anubis*** | **Olive Baboon** | 6 | 0.6 ± 1.07 | 6 | 0.6 ± 1.07 |
| 24 | ***Philantomba monticola*** | **Blue Duiker** | 6 | 0.6 ± 0.84 | 6 | 0.6 ± 0.84 |
| 25 | *Mellivora capensis* | Honey Badger | 4 | 0.4 ± 0.52 | 4 | 0.4 ± 0.52 |
| 26 | ***Panthera leo*** | **Lion** | 4 | 0.4 ± 0.52 | 4 | 0.4 ± 0.52 |

| | | | | | | |
|---|---|---|---|---|---|---|
| 27 | ***Ammotragus lervia*** | **Aoudad** | 3 | 0.3 ± 0.67 | 3 | 0.3 ± 0.67 |
| 28 | ***Ceratotherium simum simum*** | **Southern White Rhinoceros** | 3 | 0.3 ± 0.48 | 3 | 0.3 ± 0.48 |
| 29 | *Damaliscus pygargus* | Blesbok | 3 | 0.3 ± 0.67 | 3 | 0.3 ± 0.67 |
| 30 | ***Ovis ammon*** | **Wild Sheep** | 3 | 0.3 ± 0.48 | 3 | 0.3 ± 0.48 |
| 31 | ***Ovis canadensis*** | **Bighorn Sheep** | 3 | 0.3 ± 0.48 | 3 | 0.3 ± 0.48 |
| 32 | ***Capra caucasica*** | **Western Tur** | 2 | 0.2 ± 0.42 | 2 | 0.2 ± 0.42 |
| 33 | ***Cephalophus dorsalis*** | **Bay Duiker** | 2 | 0.2 ± 0.63 | 2 | 0.2 ± 0.63 |
| 34 | ***Lynx canadensis*** | **Canada Lynx** | 2 | 0.2 ± 0.63 | 2 | 0.2 ± 0.63 |
| 35 | ***Lynx rufus*** | **Bobcat** | 2 | 0.2 ± 0.42 | 2 | 0.2 ± 0.42 |
| 36 | *Pseudois nayaur* | Blue Sheep | 2 | 0.2 ± 0.63 | 2 | 0.2 ± 0.63 |
| 37 | ***Ursus maritimus*** | **Polar Bear** | 2 | 0.2 ± 0.63 | 2 | 0.2 ± 0.63 |
| 38 | ***Acinonyx jubatus*** | **Cheetah** | 1 | 0.1 ± 0.32 | 1 | 0.1 ± 0.32 |
| 39 | ***Equus zebra zebra*** | **Cape Mountain Zebra** | 1 | 0.1 ± 0.32 | 1 | 0.1 ± 0.32 |
| 40 | ***Haliaeetus leucocephalus*** | **Bald Eagle** | 1 | 0.1 ± 0.32 | 0 | 0 ± 0 |
| 41 | *Hyaena hyaena* | Striped Hyena | 1 | 0.1 ± 0.32 | 1 | 0.1 ± 0.32 |
| 42 | ***Lynx lynx*** | **Eurasian lynx** | 1 | 0.1 ± 0.32 | 1 | 0.1 ± 0.32 |
| 43 | ***Odobenus rosmarus*** | **Walrus** | 1 | 0.1 ± 0.32 | 0 | 0 ± 0 |
| 44 | ***Ovis cycloceros (O. v. cycloceros)*** | **Afghan Urial** | 1 | 0.1 ± 0.32 | 1 | 0.1 ± 0.32 |
| 45 | ***Ovis polii*** | **Marco Polo Argali** | 1 | 0.1 ± 0.32 | 1 | 0.1 ± 0.32 |
| 46 | *Proteles cristata* | Aardwolf | 1 | 0.1 ± 0.32 | 1 | 0.1 ± 0.32 |
| **Totals** | | | **1105** | | **838.87** | |

**Table S20. CITES-listed species imported to the United States as hunting trophies (2015-2024) (*n* = 177), estimated number of trophies, mean (± SD) number of trophies imported/year, WOEs imported, and mean (± SD) WOEs imported/year, ranked by trophies.**

| Rank | Scientific name | Common name | Trophies | Mean + SD trophies/year | WOEs | Mean + SD WOEs/year |
|---|---|---|---|---|---|---|
| 1 | *Ursus americanus* | American Black Bear | 31832 | 3183.2 ± 1529.9 | 16194.75 | 1619.48 ± 670.16 |
| 2 | *Giraffa camelopardalis* | Giraffe | 9234 | 923.4 ± 1757.64 | 3449 | 344.9 ± 389.03 |
| 3 | ***Crocodylus niloticus*** | **Nile Crocodile** | 6155 | 615.5 ± 885.8 | 3330.5 | 333.05 ± 192.66 |
| 4 | *Papio ursinus* | Chacma Baboon | 5895 | 589.5 ± 184.23 | 5870 | 587 ± 182.8 |
| 5 | *Hippopotamus amphibius* | Common Hippopotamus | 5063 | 506.3 ± 132.74 | 2367.077 | 236.71 ± 59.64 |
| 6 | ***Equus zebra hartmannae*** | **Hartmann's Mountain Zebra** | 5036 | 503.6 ± 131 | 4938 | 493.8 ± 129.41 |
| 7 | ***Canis lupus*** | **Grey Wolf** | 4972 | 497.2 ± 187.71 | 4525 | 452.5 ± 158.66 |
| 8 | *Meleagris ocellata* | Ocellated Turkey | 3454 | 345.4 ± 161.06 | 3335 | 333.5 ± 147.31 |
| 9 | ***Panthera pardus*** | **Leopard** | 2551 | 255.1 ± 92.66 | 2508.5 | 250.85 ± 82.6 |
| 10 | *Chlorocebus pygerythrus* | Vervet Monkey | 2323 | 232.3 ± 112.85 | 2323 | 232.3 ± 112.85 |
| 11 | ***Lynx canadensis*** | **Canada Lynx** | 2134 | 213.4 ± 145.95 | 1618.75 | 161.88 ± 64.31 |
| 12 | *Antilope cervicapra* | Blackbuck | 2051 | 205.1 ± 171.71 | 2051 | 205.1 ± 171.71 |
| 13 | *Caracal caracal* | Caracal | 1951 | 195.1 ± 91.16 | 1947 | 194.7 ± 90.86 |
| 14 | ***Loxodonta africana*** | **African Savanna Elephant** | 1894 | 189.4 ± 174.67 | 998.532 | 99.85 ± 74.66 |
| 15 | ***Ursus arctos*** | **Brown Bear** | 1739 | 173.9 ± 97.73 | 1654 | 165.4 ± 92.97 |
| 16 | *Ovis canadensis* | Bighorn Sheep | 1276 | 127.6 ± 36.32 | 1276 | 127.6 ± 36.32 |
| 17 | *Puma concolor* | Puma | 1160 | 116 ± 56.06 | 1150 | 115 ± 55.81 |
| 18 | *Capra sibirica* | Siberian Ibex | 998 | 99.8 ± 53.51 | 998 | 99.8 ± 53.51 |
| 19 | ***Kobus leche*** | **Lechwe** | 988 | 98.8 ± 36.42 | 978 | 97.8 ± 36.68 |
| 20 | *Civettictis civetta* | African Civet | 815 | 81.5 ± 32.11 | 815 | 81.5 ± 32.11 |
| 21 | *Leptailurus serval* | Serval | 788 | 78.8 ± 39.99 | 723.75 | 72.38 ± 34.94 |
| 22 | *Crax rubra* | Great Curassow | 688 | 68.8 ± 36.36 | 681 | 68.1 ± 36.33 |
| 23 | *Philantomba monticola* | Blue Duiker | 637 | 63.7 ± 29.77 | 637 | 63.7 ± 29.77 |
| 24 | *Capra hircus aegagrus (Capra aegagrus)* | Wild Goat | 614 | 61.4 ± 27.52 | 613 | 61.3 ± 27.6 |
| 25 | *Mellivora capensis* | Honey Badger | 603 | 60.3 ± 22.8 | 601 | 60.1 ± 22.77 |
| 26 | ***Damaliscus pygargus pygargus*** | **Bontebok** | 591 | 59.1 ± 46.77 | 591 | 59.1 ± 46.77 |

| | | | | | | |
|---|---|---|---|---|---|---|
| 27 | ***Ceratotherium simum simum*** | **Southern White Rhinoceros** | 584 | 58.4 ± 30.61 | 566.25 | 56.63 ± 29.41 |
| 28 | *Panthera leo* | Lion | 511 | 51.1 ± 24.85 | 491 | 49.1 ± 25.6 |
| 29 | *Papio cynocephalus* | Yellow Baboon | 488 | 48.8 ± 19.92 | 488 | 48.8 ± 19.92 |
| 30 | *Python brongersmai* | Brongersma's Short-tailed Python | 400 | 40 ± 126.49 | 400 | 40 ± 126.49 |
| 31 | *Penelope purpurascens* | Crested Guan | 395 | 39.5 ± 23.3 | 394 | 39.4 ± 23.26 |
| 32 | *Equus zebra* | Mountain Zebra | 349 | 34.9 ± 110.36 | 345 | 34.5 ± 109.1 |
| 33 | *Nasua narica* | White-nosed Coati | 338 | 33.8 ± 19.45 | 338 | 33.8 ± 19.45 |
| 34 | *Lynx rufus* | Bobcat | 273 | 27.3 ± 15.75 | 273 | 27.3 ± 15.75 |
| 35 | *Proteles cristata* | Aardwolf | 265 | 26.5 ± 10.5 | 265 | 26.5 ± 10.5 |
| 36 | ***Ovis ammon*** | **Wild Sheep** | 245 | 24.5 ± 26.44 | 245 | 24.5 ± 26.44 |
| 37 | *Felis lybica* | Afro-Asiatic Wildcat | 233 | 23.3 ± 15.14 | 232 | 23.2 ± 15.08 |
| 38 | *Papio anubis* | Olive Baboon | 195 | 19.5 ± 9.41 | 195 | 19.5 ± 9.41 |
| 39 | ***Puma concolor couguar*** | **Florida Puma** | 186 | 18.6 ± 41.23 | 184 | 18.4 ± 40.68 |
| 40 | *Capra caucasica* | Western Tur | 173 | 17.3 ± 18.86 | 173 | 17.3 ± 18.86 |
| 41 | *Felis silvestris* | European Wildcat | 169 | 16.9 ± 10.94 | 167 | 16.7 ± 10.54 |
| 42 | *Ammotragus lervia* | Aoudad | 164 | 16.4 ± 16.52 | 164 | 16.4 ± 16.52 |
| 43 | *Ortalis vetula* | Plain Chachalaca | 158 | 15.8 ± 9.66 | 158 | 15.8 ± 9.66 |
| 44 | *Ovis polii* | Marco Polo Argali | 153 | 15.3 ± 19.07 | 153 | 15.3 ± 19.07 |
| 45 | *Dendrocygna autumnalis* | Black-bellied Whistling-duck | 139 | 13.9 ± 27.09 | 139 | 13.9 ± 27.09 |
| 46 | *Dendrocygna bicolor* | Fulvous Whistling-duck | 122 | 12.2 ± 25.6 | 122 | 12.2 ± 25.6 |
| 47 | *Canis aureus* | Common Jackal | 108 | 10.8 ± 3.77 | 108 | 10.8 ± 3.77 |
| 48 | *Aquila chrysaetos* | Golden Eagle | 101 | 10.1 ± 20.31 | 101 | 10.1 ± 20.31 |
| 49 | *Malayopython reticulatus* | Reticulated Python | 100 | 10 ± 31.62 | 100 | 10 ± 31.62 |
| 50 | *Lontra canadensis* | North American River Otter | 93 | 9.3 ± 21.58 | 26 | 2.6 ± 3.27 |
| 51 | *Colobus guereza* | Guereza | 84 | 8.4 ± 4.86 | 84 | 8.4 ± 4.86 |
| 52 | *Dasyprocta punctata* | Central American Agouti | 82 | 8.2 ± 6.11 | 82 | 8.2 ± 6.11 |
| 53 | *Ovis darwini (Ovis ammon darwini)* | Gobi Argali | 81 | 8.1 ± 7.56 | 81 | 8.1 ± 7.56 |
| 54 | *Damaliscus pygargus* | Blesbok | 77 | 7.7 ± 9.18 | 77 | 7.7 ± 9.18 |
| 55 | *Ovis bochariensis (Ovis v. bochariensis)* | Bukhara Urial | 65 | 6.5 ± 5.78 | 65 | 6.5 ± 5.78 |
| 56 | *Cephalophus dorsalis* | Bay Duiker | 64 | 6.4 ± 5.48 | 64 | 6.4 ± 5.48 |
| 57 | *Papio hamadryas* | Hamadryas Baboon | 60 | 6 ± 4 | 60 | 6 ± 4 |

| | | | | | | |
|---|---|---|---|---|---|---|
| 58 | *Capra hircus* | Goat | 58 | 5.8 ± 18.34 | 58 | 5.8 ± 18.34 |
| 59 | *Axis porcinus* | Hog Deer | 55 | 5.5 ± 6.28 | 55 | 5.5 ± 6.28 |
| 60 | *Ovis cycloceros (O. v. cycloceros)* | Afghan Urial | 52 | 5.2 ± 11.32 | 52 | 5.2 ± 11.32 |
| 61 | *Pseudois nayaur* | Blue Sheep | 51 | 5.1 ± 3.63 | 51 | 5.1 ± 3.63 |
| 62 | *Capra falconeri heptneri* | Markhor | 49 | 4.9 ± 3.98 | 49 | 4.9 ± 3.98 |
| 63 | *Sarkidiornis melanotos* | Knob-billed Duck | 42 | 4.2 ± 3.97 | 42 | 4.2 ± 3.97 |
| 64 | *Capra falconeri* | Markhor | 41 | 4.1 ± 2.28 | 41 | 4.1 ± 2.28 |
| 65 | *Cephalophus silvicultor* | Yellow-backed Duiker | 38 | 3.8 ± 3.12 | 38 | 3.8 ± 3.12 |
| 66 | ***Acinonyx jubatus*** | **Cheetah** | 37 | 3.7 ± 3.2 | 37 | 3.7 ± 3.2 |
| 67 | *Ovis cycloceros cycloceros (O.v. cycloceros)* | Afghan Urial (subsp. cycloceros) | 36 | 3.6 ± 6.1 | 36 | 3.6 ± 6.1 |
| 68 | *Bubalus arnee* | Wild Water Buffalo | 35 | 3.5 ± 8.81 | 9 | 0.9 ± 1.91 |
| 69 | *Ovis aries* | Urial | 31 | 3.1 ± 6.64 | 31 | 3.1 ± 6.64 |
| 70 | *Python sebae* | Central African Rock Python | 31 | 3.1 ± 2.92 | 31 | 3.1 ± 2.92 |
| 71 | *Ceratotherium simum* | White Rhinoceros | 29 | 2.9 ± 2.92 | 28.25 | 2.83 ± 2.75 |
| 72 | *Ovis punjabiensis (Ovis v. punjabensis)* | Punjab Urial | 25 | 2.5 ± 4.06 | 25 | 2.5 ± 4.06 |
| 73 | ***Theropithecus gelada*** | **Gelada** | 23 | 2.3 ± 1.83 | 23 | 2.3 ± 1.83 |
| 74 | *Philantomba maxwellii* | Maxwell's Duiker | 20 | 2 ± 2.67 | 20 | 2 ± 2.67 |
| 75 | *Erythrocebus patas* | Patas Monkey | 18 | 1.8 ± 2.62 | 18 | 1.8 ± 2.62 |
| 76 | *Chlorocebus aethiops* | Grivet Monkey | 17 | 1.7 ± 2.83 | 17 | 1.7 ± 2.83 |
| 77 | *Cuniculus paca* | Paca | 16 | 1.6 ± 1.26 | 16 | 1.6 ± 1.26 |
| 78 | *Canis lupus monstrabilis* | Texas Grey Wolf | 15 | 1.5 ± 3.44 | 15 | 1.5 ± 3.44 |
| 79 | ***Equus zebra zebra*** | **Cape Mountain Zebra** | 15 | 1.5 ± 2.01 | 12 | 1.2 ± 1.4 |
| 80 | *Mazama temama* | Central American Red Brocket | 15 | 1.5 ± 4.4 | 15 | 1.5 ± 4.4 |
| 81 | *Antigone canadensis (Grus canadensis)* | Sandhill Crane | 14 | 1.4 ± 1.84 | 14 | 1.4 ± 1.84 |
| 82 | *Cephalophus zebra* | Zebra Duiker | 13 | 1.3 ± 4.11 | 13 | 1.3 ± 4.11 |
| 83 | *Cercopithecus mitis* | Blue Monkey | 12 | 1.2 ± 1.32 | 12 | 1.2 ± 1.32 |
| 84 | ***Crocodylus porosus*** | **Saltwater Crocodile** | 12 | 1.2 ± 1.48 | 12 | 1.2 ± 1.48 |
| 85 | *Hyaena hyaena* | Striped Hyena | 12 | 1.2 ± 1.48 | 12 | 1.2 ± 1.48 |
| 86 | *Necrosyrtes monachus* | Hooded Vulture | 12 | 1.2 ± 2.57 | 12 | 1.2 ± 2.57 |
| 87 | ***Bison bison athabascae*** | **Wood Bison** | 11 | 1.1 ± 2.02 | 10 | 1 ± 1.76 |
| 88 | *Lynx lynx* | Eurasian lynx | 9 | 0.9 ± 1.1 | 9 | 0.9 ± 1.1 |

| | | | | | | |
|---|---|---|---|---|---|---|
| 89 | *Ursus thibetanus* | Asiatic Black Bear | 9 | 0.9 ± 2.51 | 9 | 0.9 ± 2.51 |
| 90 | *Mazama temama cerasina* | Guatemalan Red Brocket | 8 | 0.8 ± 1.87 | 8 | 0.8 ± 1.87 |
| 91 | *Myiopsitta monachus* | Monk Parakeet | 8 | 0.8 ± 2.53 | 8 | 0.8 ± 2.53 |
| 92 | ***Oryx dammah*** | **Scimitar-horned Oryx** | 8 | 0.8 ± 1.87 | 8 | 0.8 ± 1.87 |
| 93 | *Papio papio* | Guinea Baboon | 8 | 0.8 ± 2.53 | 8 | 0.8 ± 2.53 |
| 94 | *Tauraco livingstonii* | Livingstone's Turaco | 8 | 0.8 ± 2.53 | 8 | 0.8 ± 2.53 |
| 95 | *Varanus albigularis* | White-throated Monitor | 8 | 0.8 ± 1.55 | 8 | 0.8 ± 1.55 |
| 96 | *Gyps africanus* | White-backed Vulture | 7 | 0.7 ± 2.21 | 7 | 0.7 ± 2.21 |
| 97 | *Haliaeetus vocifer* | African Fish-eagle | 7 | 0.7 ± 2.21 | 7 | 0.7 ± 2.21 |
| 98 | *Kaupifalco monogrammicus* | Lizard Buzzard | 7 | 0.7 ± 2.21 | 7 | 0.7 ± 2.21 |
| 99 | *Odobenus rosmarus* | Walrus | 7 | 0.7 ± 2.21 | 1 | 0.1 ± 0.32 |
| 100 | *Otolemur crassicaudatus* | Thick-tailed Greater Galago | 7 | 0.7 ± 1.34 | 7 | 0.7 ± 1.34 |
| 101 | *Athene cunicularia* | Burrowing Owl | 6 | 0.6 ± 1.9 | 6 | 0.6 ± 1.9 |
| 102 | *Bison bison* | American Bison | 6 | 0.6 ± 1.9 | 5 | 0.5 ± 1.58 |
| 103 | *Circaetus fasciolatus* | Southern Banded Snake-eagle | 6 | 0.6 ± 1.9 | 6 | 0.6 ± 1.9 |
| 104 | ***Diceros bicornis*** | **Black Rhinoceros** | 6 | 0.6 ± 0.97 | 6 | 0.6 ± 0.97 |
| 105 | *Galago moholi* | Southern Lesser Galago | 6 | 0.6 ± 1.26 | 6 | 0.6 ± 1.26 |
| 106 | *Strix woodfordii* | African Wood-owl | 6 | 0.6 ± 1.9 | 6 | 0.6 ± 1.9 |
| 107 | ***Capra falconeri megaceros*** | **Straight-horned Markhor** | 5 | 0.5 ± 0.97 | 5 | 0.5 ± 0.97 |
| 108 | *Glaucidium capense* | African Barred Owlet | 5 | 0.5 ± 1.58 | 5 | 0.5 ± 1.58 |
| 109 | *Pavo cristatus* | Indian Peafowl | 5 | 0.5 ± 0.97 | 5 | 0.5 ± 0.97 |
| 110 | *Poicephalus cryptoxanthus* | Brown-headed Parrot | 5 | 0.5 ± 1.58 | 5 | 0.5 ± 1.58 |
| 111 | *Cephalophus ogilbyi* | Ogilby's Duiker | 4 | 0.4 ± 0.97 | 4 | 0.4 ± 0.97 |
| 112 | *Tauraco porphyreolophus* | Purple-crested Turaco | 4 | 0.4 ± 0.84 | 4 | 0.4 ± 0.84 |
| 113 | *Trigonoceps occipitalis* | White-headed Vulture | 4 | 0.4 ± 1.26 | 4 | 0.4 ± 1.26 |
| 114 | *Chlorocebus tantalus* | Tantalus Monkey | 3 | 0.3 ± 0.95 | 3 | 0.3 ± 0.95 |
| 115 | *Gyps rueppellii* | Rüppell's Vulture | 3 | 0.3 ± 0.95 | 3 | 0.3 ± 0.95 |
| 116 | *Ovis cycloceros arkal (O.v. arkal)* | Afghan Urial (subsp. arkal) | 3 | 0.3 ± 0.95 | 3 | 0.3 ± 0.95 |
| 117 | *Phoeniconaias minor* | Lesser Flamingo | 3 | 0.3 ± 0.95 | 3 | 0.3 ± 0.95 |
| 118 | *Poicephalus fuscicollis* | Brown-necked Parrot | 3 | 0.3 ± 0.95 | 3 | 0.3 ± 0.95 |
| 119 | *Sagittarius serpentarius* | Secretarybird | 3 | 0.3 ± 0.67 | 3 | 0.3 ± 0.67 |

| | | | | | | |
|---|---|---|---|---|---|---|
| 120 | *Sarcogyps calvus* | Red-headed Vulture | 3 | 0.3 ± 0.95 | 3 | 0.3 ± 0.95 |
| 121 | *Boselaphus tragocamelus* | Nilgai | 2 | 0.2 ± 0.42 | 2 | 0.2 ± 0.42 |
| 122 | *Buteo augur* | Augur Buzzard | 2 | 0.2 ± 0.63 | 2 | 0.2 ± 0.63 |
| 123 | *Caiman crocodilus* | Spectacled Caiman | 2 | 0.2 ± 0.63 | 2 | 0.2 ± 0.63 |
| 124 | *Chlorocebus sabaeus* | Green Monkey | 2 | 0.2 ± 0.63 | 2 | 0.2 ± 0.63 |
| 125 | *Ciconia nigra* | Black Stork | 2 | 0.2 ± 0.63 | 2 | 0.2 ± 0.63 |
| 126 | *Colibri delphinae* | Brown Violetear | 2 | 0.2 ± 0.63 | 2 | 0.2 ± 0.63 |
| 127 | *Galago senegalensis* | Northern Lesser Galago | 2 | 0.2 ± 0.63 | 2 | 0.2 ± 0.63 |
| 128 | *Gypohierax angolensis* | Palm-nut Vulture | 2 | 0.2 ± 0.63 | 2 | 0.2 ± 0.63 |
| 129 | *Hydrictis maculicollis* | Spotted-necked Otter | 2 | 0.2 ± 0.63 | 2 | 0.2 ± 0.63 |
| 130 | *Loxodonta cyclotis* | African Forest Elephant | 2 | 0.2 ± 0.63 | 2 | 0.2 ± 0.63 |
| 131 | *Moschus moschiferus* | Siberian Musk Deer | 2 | 0.2 ± 0.42 | 2 | 0.2 ± 0.42 |
| 132 | *Odocoileus virginianus mayensis* | Guatemalan White-tailed Deer | 2 | 0.2 ± 0.63 | 2 | 0.2 ± 0.63 |
| 133 | *Ovis gmelini* | Mouflon | 2 | 0.2 ± 0.63 | 2 | 0.2 ± 0.63 |
| 134 | *Pecari tajacu* | Collared Pecary | 2 | 0.2 ± 0.42 | 2 | 0.2 ± 0.42 |
| 135 | *Perodicticus potto* | West African Potto | 2 | 0.2 ± 0.63 | 2 | 0.2 ± 0.63 |
| 136 | *Polyboroides typus* | African Harrier-hawk | 2 | 0.2 ± 0.63 | 2 | 0.2 ± 0.63 |
| 137 | *Syrmaticus reevesii* | Reeves's Pheasant | 2 | 0.2 ± 0.63 | 2 | 0.2 ± 0.63 |
| 138 | *Terathopius ecaudatus* | Bateleur | 2 | 0.2 ± 0.63 | 2 | 0.2 ± 0.63 |
| 139 | *Ursus americanus emmonsii* | Emmon's Black Bear | 2 | 0.2 ± 0.42 | 2 | 0.2 ± 0.42 |
| 140 | ***Ursus maritimus*** | **Polar Bear** | 2 | 0.2 ± 0.63 | 2 | 0.2 ± 0.63 |
| 141 | *Accipiter melanoleucus* | Black Sparrowhawk | 1 | 0.1 ± 0.32 | 1 | 0.1 ± 0.32 |
| 142 | ***Alligator mississippiensis*** | **American Alligator** | 1 | 0.1 ± 0.32 | 1 | 0.1 ± 0.32 |
| 143 | *Aonyx capensis* | African Clawless Otter | 1 | 0.1 ± 0.32 | 1 | 0.1 ± 0.32 |
| 144 | *Aquila rapax* | Tawny Eagle | 1 | 0.1 ± 0.32 | 1 | 0.1 ± 0.32 |
| 145 | *Ardeotis kori* | Kori Bustard | 1 | 0.1 ± 0.32 | 1 | 0.1 ± 0.32 |
| 146 | *Asio capensis* | Marsh Owl | 1 | 0.1 ± 0.32 | 1 | 0.1 ± 0.32 |
| 147 | *Balearica regulorum* | Grey Crowned Crane | 1 | 0.1 ± 0.32 | 0 | 0 ± 0 |
| 148 | *Branta canadensis leucopareia* | Aleutian Canada Goose | 1 | 0.1 ± 0.32 | 1 | 0.1 ± 0.32 |
| 149 | *Buteo buteo* | Eurasian Buzzard | 1 | 0.1 ± 0.32 | 1 | 0.1 ± 0.32 |
| 150 | ***Caiman crocodilus crocodilus*** | **South American Spectacled Caiman** | 1 | 0.1 ± 0.32 | 1 | 0.1 ± 0.32 |

| | | | | | | |
|---|---|---|---|---|---|---|
| 151 | *Cercopithecus neglectus* | De Brazza's Monkey | 1 | 0.1 ± 0.32 | 1 | 0.1 ± 0.32 |
| 152 | *Circaetus pectoralis* | Black-chested Snake-eagle | 1 | 0.1 ± 0.32 | 1 | 0.1 ± 0.32 |
| 153 | *Crotalus durissus* | Cascabel Rattlesnake | 1 | 0.1 ± 0.32 | 1 | 0.1 ± 0.32 |
| 154 | *Eunectes murinus* | Green Anaconda | 1 | 0.1 ± 0.32 | 1 | 0.1 ± 0.32 |
| 155 | *Eupodotis senegalensis* | White-bellied Bustard | 1 | 0.1 ± 0.32 | 1 | 0.1 ± 0.32 |
| 156 | *Falco chicquera* | Red-headed Falcon | 1 | 0.1 ± 0.32 | 1 | 0.1 ± 0.32 |
| 157 | *Gazella bennettii* | Chinkara | 1 | 0.1 ± 0.32 | 1 | 0.1 ± 0.32 |
| 158 | *Lissotis melanogaster* | Black-bellied Bustard | 1 | 0.1 ± 0.32 | 1 | 0.1 ± 0.32 |
| 159 | *Lophaetus occipitalis* | Long-crested Eagle | 1 | 0.1 ± 0.32 | 1 | 0.1 ± 0.32 |
| 160 | *Manis tricuspis* | White-bellied Pangolin | 1 | 0.1 ± 0.32 | 1 | 0.1 ± 0.32 |
| 161 | *Melierax poliopterus* | Eastern Chanting-goshawk | 1 | 0.1 ± 0.32 | 1 | 0.1 ± 0.32 |
| 162 | *Micronisus gabar* | Gabar Goshawk | 1 | 0.1 ± 0.32 | 1 | 0.1 ± 0.32 |
| 163 | *Mustela erminea* | Stoat | 1 | 0.1 ± 0.32 | 1 | 0.1 ± 0.32 |
| 164 | *Odocoileus virginianus* | White-tailed Deer | 1 | 0.1 ± 0.32 | 1 | 0.1 ± 0.32 |
| 165 | *Phoenicopterus roseus* | Greater Flamingo | 1 | 0.1 ± 0.32 | 0 | 0 ± 0 |
| 166 | *Phoenicopterus ruber* | American Flamingo | 1 | 0.1 ± 0.32 | 1 | 0.1 ± 0.32 |
| 167 | *Polemaetus bellicosus* | Martial Eagle | 1 | 0.1 ± 0.32 | 1 | 0.1 ± 0.32 |
| 168 | *Polihierax semitorquatus* | African Pygmy-falcon | 1 | 0.1 ± 0.32 | 1 | 0.1 ± 0.32 |
| 169 | *Python regius* | Ball Python | 1 | 0.1 ± 0.32 | 1 | 0.1 ± 0.32 |
| 170 | *Saguinus midas* | Golden-handed Tamarin | 1 | 0.1 ± 0.32 | 1 | 0.1 ± 0.32 |
| 171 | *Saimiri sciureus* | Guianan Squirrel Monkey | 1 | 0.1 ± 0.32 | 1 | 0.1 ± 0.32 |
| 172 | *Stigmochelys pardalis* | Leopard Tortoise | 1 | 0.1 ± 0.32 | 1 | 0.1 ± 0.32 |
| 173 | *Tayassu pecari* | White-lipped Peccary | 1 | 0.1 ± 0.32 | 1 | 0.1 ± 0.32 |
| 174 | *Torgos tracheliotus* | Lappet-faced Vulture | 1 | 0.1 ± 0.32 | 1 | 0.1 ± 0.32 |
| 175 | *Varanus salvator* | Common Water Monitor | 1 | 0.1 ± 0.32 | 1 | 0.1 ± 0.32 |
| 176 | *Viverra civettina* | Malabar Civet | 1 | 0.1 ± 0.32 | 1 | 0.1 ± 0.32 |
| 177 | *Vulpes vulpes* | Red Fox | 1 | 0.1 ± 0.32 | 1 | 0.1 ± 0.32 |
| **Totals** | | | **102,999** | | **73,576.36** | |

**Supplementary Data 1. Population status of CITES-listed species traded internationally as hunting trophies (2015-2024) in exporting countries.** Species ranked by number of trophies.

**Taxonomic notes:**

- Exports of the African Savanna elephant (*Loxodonta africana*) from Cameroon may refer to the African Forest elephant (*Loxodonta cyclotis*) as Cameroon has populations of both species are per the Red List.
- Exports of *Felis silvestris* from Ethiopia and Namibia should be considered as *F. lybica* on the Red List due to taxonomic differences between the CITES trade database and the Red List.
- Exports of *Canis aureus* from Cameroon, Ethiopia, Chad and Tanzania should be treated as *C. lupaster* on the Red List due to taxonomic differences between the CITES trade database and the Red List.
- Exports of *Papio cynocephalus* from South Africa likely refer to *P. ursinus* and these records reflect a CITES nomenclatural issue.
- Trade records for *Philantomba maxwelli* likely refer *P. monticola; P. maxwelli* was formerly a synonym.
- Exports of *Papio hamadryas* from South Africa likely refer to *P. ursinus*.
- It is unlikely exports of *Chlorocebus aethiops* from South Africa and Zimbabwe refer to this species and reflect a CITES nomenclatural issue.
- Exports of *Papio anubis* from South Africa likely reflect a CITES nomenclatural issue.

**Abbreviations:** AE = United Arab Emirates, AM = Armenia, AR = Argentina, AU = Australia, AZ = Azerbaijan, BF = Burkina Faso, BG = Bulgaria, BJ = Benin, BW = Botswana, CA = Canada, CF = Central African Republic, CG = Congo, CM = Cameroon, CZ = Czech Republic, DE = Germany, EE = Estonia, ES = Spain, ET = Ethiopia, FI = Finland, GB = United Kingdom of Great Britain and Northern Ireland, GH = Ghana, HR = Croatia, HU = Hungary, ID = Indonesia, IR = Iran (Islamic Republic of), KG = Kyrgyzstan, KZ = Kazakhstan, LR = Liberia, LT = Lithuania, LV = Latvia, MA = Morocco, MK = North Macedonia, MN = Mongolia, MX = Mexico, MY = Malaysia, MZ = Mozambique, NA = Namibia, NP = Nepal, NZ = New Zealand, PK = Pakistan, PY = Paraguay, RO = Romania, RS = Serbia, RU = Russian Federation, SE = Sweden, SI = Slovenia, SN = Senegal, SR = Suriname, TD = Chad, TJ = Tajikistan, TR = Türkiye, TZ = United Republic of Tanzania, UG = Uganda, US = United States of America, UZ = Uzbekistan, ZA = South Africa, ZM = Zambia, ZW = Zimbabwe.

[see MS Excel file attached].
